# Global tree encoding of atlas-scale single-cell genomics

**DOI:** 10.64898/2026.08.31.747971

**Authors:** Brett Kiyota, Chaehyeon Lee, Haoyang Yao, Nozomu Yachie

**Affiliations:** School of Biomedical Engineering, The University of British Columbia, Vancouver, BC, Canada; Premium Research Institute for Human Metaverse Medicine (WPI-PRIMe), The University of Osaka, Suita, Osaka, Japan; Research Center for Advanced Science and Technology, The University of Tokyo, Tokyo, Japan; CREST Cell Control Program, Japan Science and Technology Agency, Core Research for Evolutional Science and Technology, Kawaguchi-shi, Saitama, Japan; The McMillan Multiscale Human Program, the Canadian Institute for Advanced Research, Toronto, Canada

## Abstract

The rapid expansion of single-cell genomic datasets has led to the compilation of biological resources comprising hundreds of millions of cells across tissues, developmental stages, and disease states. This has underscored the need for scalable and interpretable data representations that preserve the complex relationships and multi-scale organization of cellular states, while remaining computationally tractable at atlas scale. Existing approaches based on discrete abstractions have enabled cell annotation, clustering, and trajectory inference, but are often optimized for local inference tasks and may obscure continuous cellular relationships and multi-resolution structure within complex transcriptional and other genomic landscapes. Moreover, increasing dataset sizes often require information-reduction strategies such as random downsampling, limiting the resolution of rare cell populations and heterogeneous cellular states. Here, we present MILK, a scalable computational framework that organizes high-dimensional single-cell populations into unified tree representations. Across large-scale transcriptomic atlases, MILK enables representative subsampling with preserved information, supporting the tractable application of existing algorithms for tasks including deep generative model training and foundation model benchmarking. Additionally, MILK enables holistic, multi-resolution analyses that capture global developmental trajectories, characterize disease-associated cellular perturbations across tissues, and facilitate comparison of transcriptional programs across species within a coherent hierarchical framework. Together, these results establish the hierarchical organization of biological data as a scalable and unifying representation of cellular identity, enabling integrative analysis of single-cell genomic data across diverse contexts.

## INTRODUCTION

Advances in single-cell genomics technologies^1–3^ have enabled the comprehensive profiling of complex biological systems at unprecedented resolution and scale. We are currently facing a deluge of information across diverse tissues^4^, developmental stages^5^, and disease contexts^6^. This has motivated organized efforts to compile and harmonize the molecular readouts of hundreds of millions of cells, as exemplified by resources such as the Chan-Zuckerberg CELLxGENE Discover Census^7^, Human Cell Atlas^8^, and Arc Virtual Cell Atlas^9^. As these resources rapidly expand toward billions of cells through initiatives such as the CZI Billion Cells Project^10^, it marks a shift in the central bottleneck of single-cell biology from data acquisition to the efficient extraction of interpretable information from high-dimensional molecular measurements. However, despite their promise for deeper understanding of biological processes, the global insights obtained from these vast resources have been limited, suggesting that much of this rich information remains underutilized^11^.

Single-cell RNA sequencing (scRNA-seq) is a methodology that quantifies expression across thousands of genes, enabling populations of cells to be represented as point clouds in a high-dimensional vector space^12,13^. Despite being impacted by a multitude of technical biases^14^ and stochastic biological factors^15–17^, transcriptomic profiling has been instrumental in characterizing the heterogeneity of cellular states and their variation across biological contexts in time and space^18,19^. While the diverse array of analytical tools developed for these data has shaped our understanding of biological processes, extracting holistic insights from complex transcriptional landscapes remains a major challenge.

Cells in multicellular systems often occupy overlapping or closely related states, forming continuous differentiation trajectories shaped by cell lineage and intercellular communication^20,21^. Such cellular states are inherently structured across multiple scales, reflecting the nested organization of biological systems^22^. However, current single-cell embeddings and model-learning frameworks are largely developed without explicitly accounting for this structure. In practice, different analytical choices can yield fragmented and sometimes discrepant views of cell identity^1^. Current workflows frequently discretize cellular states into distinct subpopulations for subsequent analysis in terms of their distributions^23^, cell-state transitions^20^, and genetic programs^24^. Interpretation is often further guided by lower-dimensional projections, such as t-SNE (t-distributed stochastic neighbor embedding)^25^ or UMAP (uniform manifold approximation and projection)^26^, which prioritize local structure but are not guaranteed to preserve global relationships. Collectively, these approaches facilitate human interpretation but can obscure intermediate- and long-range dependencies, limiting our ability to capture the multi-scale organization of biological systems. There is a fundamental mismatch between the high-dimensional, continuous nature of cellular states and the representations used to encode them.

Tree-like structures provide a fundamental language for representing biology across scales. Evolutionary diversification is conventionally summarized by phylogenies, in which species and molecular sequences branch through time as lineages diversify under changing environments. Development of multicellular organisms is also rooted in branching processes, in which a single fertilized egg gives rise to cellular lineages through iterative cell divisions, organized by gene regulatory programs and cell–cell interactions that produce progressively specialized cell states. Thus, both cell lineage and cell-state diversification can be viewed, at least in part, through hierarchical tree structures that capture major aspects of development. Because genome evolution and genome-encoded regulatory programs ultimately shape gene expression and other molecular phenotypes, single-cell molecular profiles can be regarded as intermediate readouts of these underlying evolutionary and developmental processes. Consistent with this view, recent work has shown that tree representations of single-cell transcriptomes can support biological interpretation, noise regularization and multi-scale exploration^27^. However, existing tree-based approaches have yet to provide a general framework for interrogating atlas-scale single-cell resources comprising millions to over tens of millions of cells across diverse datasets, biological contexts and metadata annotations. In particular, it remains unclear which biological signals can be decoded from global tree representations, how such structures can support quantitative metadata-driven analyses, and how they can be used to complement increasingly complex foundation-model embeddings of single-cell atlases.

The rapid growth of single-cell data has also facilitated the integration of gene expression profiles from diverse datasets into a shared space. While such meta-analyses enable identification of emergent biological patterns, they present additional technical challenges, most notably batch effects, or the technical variation that arises from differences in experimental protocols, sample handling, and sequencing technologies^28–30^. A growing class of batch normalization methods has been developed to address these systematic discrepancies^31^. Some align datasets by identifying shared cellular states across batches^32,33^, whereas others employ probabilistic or generative models to learn the underlying distribution that separates technical variation from biological signal^34,35^. More recently, foundation models trained on large-scale scRNA-seq data have exhibited promise in learning low-dimensional, generalizable embeddings that implicitly integrate data across batches and experimental conditions^36–38^. However, these approaches remain difficult to systematically evaluate at scale^31^, particularly as the number of independent datasets grows.

While access to more information can potentially uncover underappreciated biological phenomena, the computing costs of analyzing millions of cells become prohibitive for widely used algorithms^13,30^. Computationally, this often necessitates the use of heuristic approximations^13,39^ and modeling assumptions that can distort biological information, often sacrificing the granularity desired from the rich molecular data. Another common practice involves the downsampling of data to computationally tractable sizes^4,13,19,40^, which can lead to substantial loss of information and biases in population structure. These scaling pressures introduce an additional layer of challenges that can hinder analyses targeting more integrative and comprehensive understanding of population dynamics.

More broadly, foundation models have rapidly emerged as an effective paradigm for leveraging the abundance of single-cell information available. The process of learning latent representations that capture functional signatures in transcriptomic profiles underpins their utility across a broad range of predictive inference tasks^36–38,41,42^. However, understanding how predictions are made often remains a challenge^43^, limiting the mechanistic interpretability of downstream biological insights. Furthermore, these approaches often require substantial computational resources for model training, excluding their accessibility to research groups with rich infrastructure^44^.

A scalable approach for inferring hierarchical representations of large-scale cell populations, should such a framework be achievable, would provide several advantages. First, it could generate balanced representations of cell populations within an intuitive tree-based structure, while simultaneously encoding complex biological signals. Second, by linking similar cells to representative exemplars, it would enable equitable data compression while preserving underlying structure and rare populations, which could support tractable global analyses and model training. Third, mathematical measures defined on tree topology, together with associated metadata labels, would enable data-driven analyses across multiple biological resolutions and provide a principled framework for assessing data integration methods, while requiring fewer tunable parameters and assumptions. Finally, such a hierarchical representation may provide a natural framework for contextualizing and interpreting predictions derived from foundation models or additional modalities of single-cell information.

Here, we present MILK (multi-resolution integration of large-scale and high-dimensional kernel information), a method that captures hierarchical representations of cell populations at unprecedented scales to enable both new biological observations and systematic assessment of single-cell analysis frameworks (**Fig. 1**). Although broadly applicable to high-dimensional datasets, we applied MILK extensively to large-scale scRNA-seq atlases and show that “transcriptional trees” constructed by MILK can efficiently compress data while preserving biological information (**Fig. 2**), capture developmental cell differentiation trajectories (**Fig. 3**), benchmark single-cell foundation models (**Fig. 4**), globally contextualize cells across complex disease contexts (**Fig. 5**), and support comparative analyses of transcriptional program evolution across diverse cell types (**Fig. 6**).

**Fig. 1.**
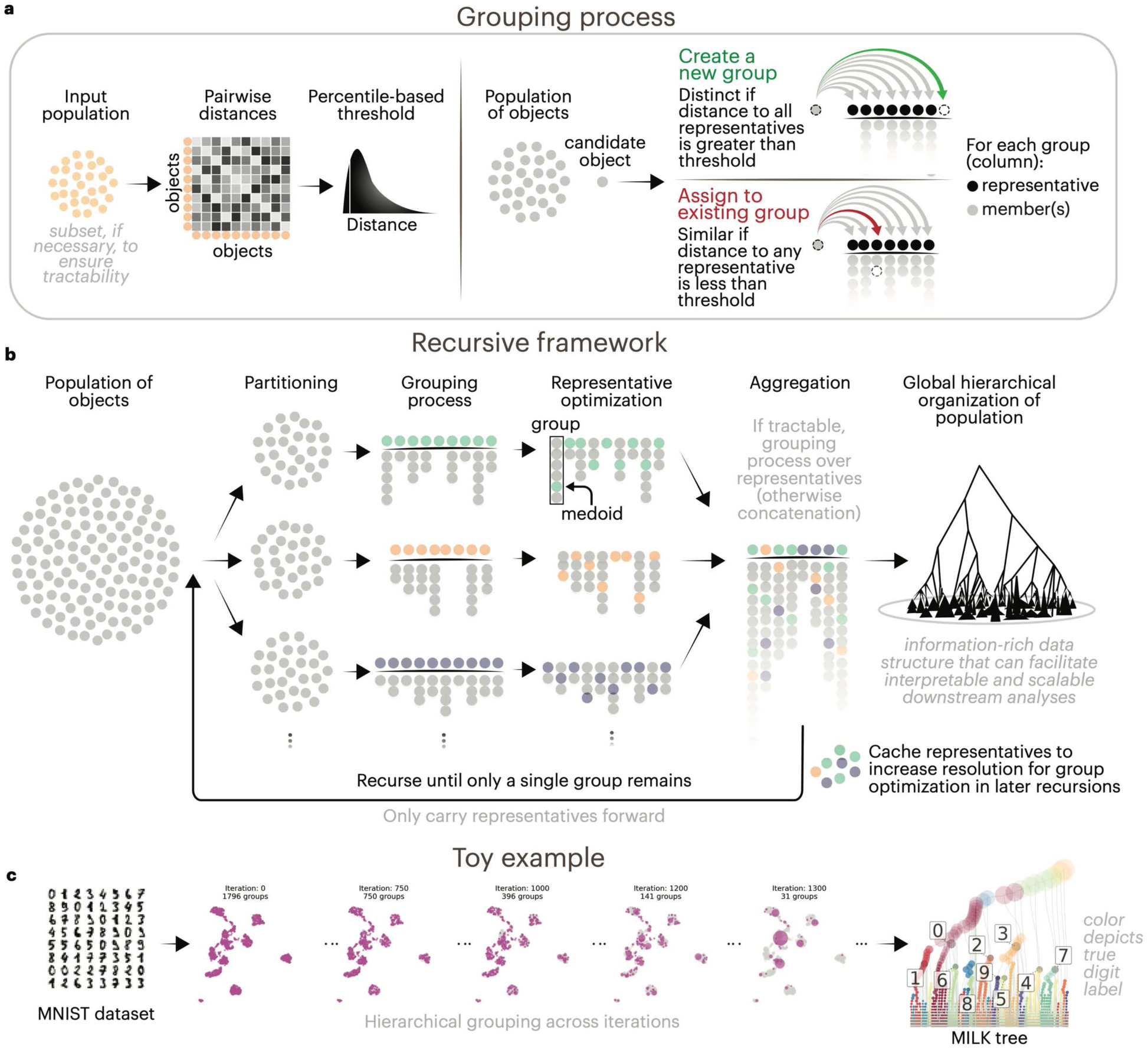
Overview of MILK. **a,** Core distance-based grouping process. An approximate global distribution of pairwise distances is estimated from randomly sampled objects, and an extreme-proximity threshold is defined as a lower-tail percentile of this distribution. Candidate objects are then processed in a single pass. Objects outside the threshold distance from all existing representatives are assigned as representatives of new groups, whereas objects within the threshold distance of an existing representative are assigned to their most similar group. Group representatives are subsequently updated as medoids of their assigned members. **b,** Recursive hierarchy construction. The core grouping process is applied recursively to progressively merge representatives from fine to coarse resolutions. For large datasets, input objects are partitioned for parallel processing, followed by representative aggregation when computationally tractable. Cached representatives from earlier recursive cycles are used to refine medoid selection in later iterations. Recursion continues until a single root group remains, yielding a hierarchical tree representation of the complete population. **c,** MNIST demonstration. MILK applied to the MNIST dataset of pixelated handwritten digits. Representative objects are shown in magenta within the UMAP embedding of the complete dataset in light gray, with point size proportional to group size. The corresponding MILK tree is annotated by ground-truth digit labels.

**Fig. 2.**
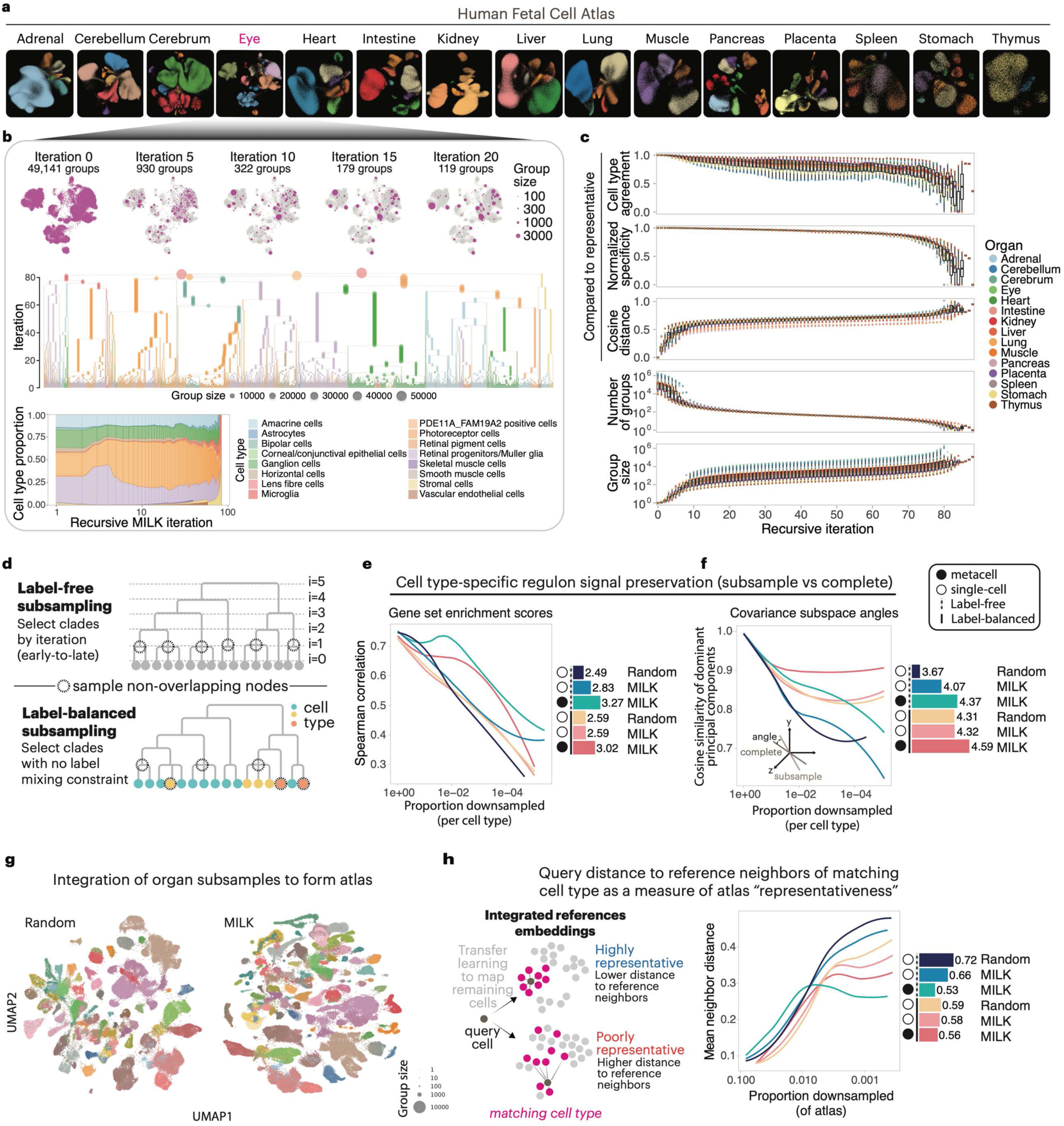
Information-rich subsampling of a 4M human fetal cell atlas. **a,** Human fetal organ datasets. UMAP embeddings of 15 processed human fetal organ scRNA-seq datasets, colored by cell type. **b,** Recursive MILK hierarchy construction. MILK trees were constructed from 100 principal components of each organ dataset using cosine distance and a similarity threshold set to the 0.1 percentile. The eye dataset is shown as a representative example. Top, group representatives in magenta overlaid on the UMAP embedding of the complete organ dataset in light gray across selected recursive iterations; point size denotes group size. Middle, complete MILK tree colored by cell type labels, with each level on the y-axis corresponding to a recursive iteration. Bottom, alluvial plot showing the relative composition of cell types among group representatives across recursive iterations. **c,** Recursive MILK summary statistics. Distributions of representative cell type agreement, normalized specificity, cosine distance to representative cells, number of groups, and group size across recursive MILK iterations for all 15 organs. **d,** Representative subsampling strategies. Label-free and label-balanced subsampling strategies for selecting non-overlapping representative clades from MILK trees. Organ MILK trees were sampled with both strategies across target sample sizes of 100, 300, 1,000, 3,000, 10,000, and 30,000 cells per organ. **e,** Regulon signal preservation. Preservation of cell type-specific regulon enrichment profiles after downsampling. Spearman correlations between regulon activity profiles from subsampled and complete organ datasets were calculated for each cell type and summarized across sampling proportions. Lines indicate LOESS fits across cell types. **f,** Regulon covariance preservation. Preservation of regulon-associated transcriptomic covariance after downsampling. Cell type-specific covariance structure was assessed by the average cosine similarity of dominant principal-component subspace angles between subsampled and complete datasets. Lines indicate LOESS fits across cell types. **g,** Integrated fetal reference atlases. scVI-integrated atlases generated from organ subsamples with a target size of 10,000 cells per organ. UMAP embeddings are shown for random subsampling and MILK-based label-balanced subsampling with metacell aggregation; colors denote cell types, and point size denotes the number of cells represented by each sampled group. **h,** Reference-atlas representativeness by query-cell mapping. Non-subsampled query cells were mapped onto scVI-integrated reference atlases using scArches. Representativeness was evaluated as the average cosine distance from each query cell to its eight nearest reference neighbors of the same cell type, with lower values indicating better representation of the complete atlas. Curves show mean neighbor distance as a function of atlas sampling proportion, and AUC values summarize performance for each subsampling approach.

**Fig. 3.**
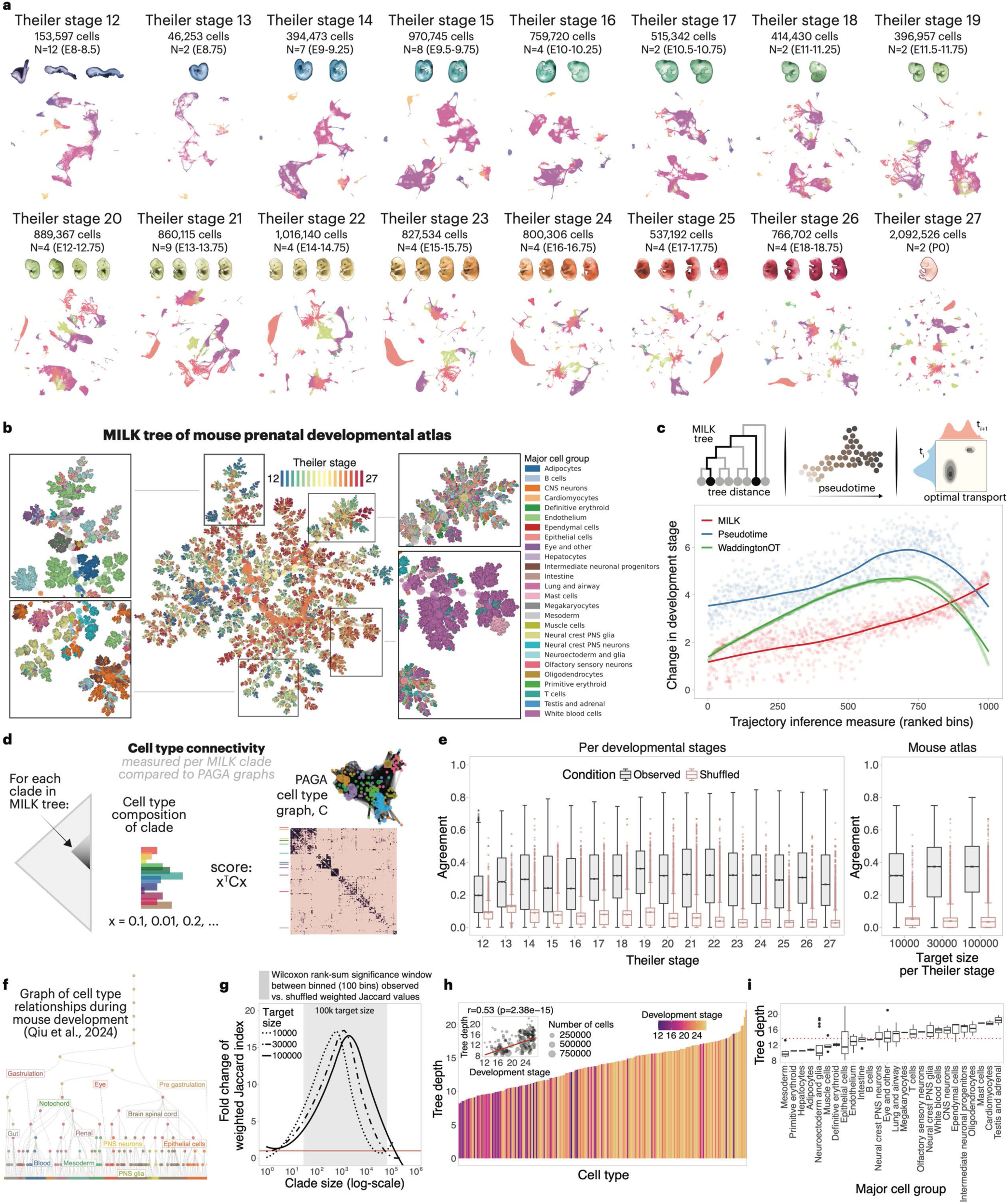
Single-cell developmental trajectories in a mouse prenatal atlas. **a,** Mouse prenatal developmental atlas. UMAP embedding of the mouse prenatal developmental scRNA-seq atlas reported by Qiu et al.^19^, comprising 83 embryos from late gastrulation to birth across 16 Theiler stages. **b,** Global developmental MILK hierarchy. Global MILK tree reconstructed from a cell type- and Theiler stage-balanced representative atlas, generated by label-balanced subsampling of each Theiler stage to a target size of 30,000 cells. Nodes in the middle are colored by Theiler stage, and selected zoom-up examples are colored by cell type. Node size denotes group size. **c,** Developmental progression captured by MILK tree distance. Absolute differences in Theiler stage between cell pairs were compared across three trajectory measures: MILK tree distance, Diffusion Pseudotime, and the complement of expected transport mass from Waddington-OT. For comparison across methods, trajectory measures were ranked and partitioned into 1,000 equal-sized bins before calculating the average Theiler-stage difference per bin. **d,** PAGA-based cell type connectivity comparison. Cell type frequency profiles were calculated for MILK clades and compared with the PAGA-derived cell type connectivity matrix using a quadratic form. **e,** PAGA-concordant cell type organization in MILK clades. Distributions of clade-level agreement scores between MILK cell type frequency profiles and the PAGA cell type connectivity graph are shown for Theiler stage-specific and global developmental MILK trees, alongside shuffled cell type-label controls. Bonferroni-adjusted Wilcoxon rank-sum tests were used to compare observed and shuffled agreement scores. **f,** Curated developmental cell type relationship graph. Tree representation of cell type relationships during mouse development derived from the original atlas annotation framework. Node color denotes major cell group. **g,** Intermediate-resolution agreement with developmental cell type relationships. Cell type relationships encoded by clades in the single-cell MILK tree were compared against the curated developmental cell type lineage graph in Qiu et al.^19^ using an asymmetric weighted Jaccard index. LOESS-fitted curves show fold change in clade agreement scores relative to shuffled cell type-label controls across clade sizes. Bonferroni-adjusted Wilcoxon rank-sum tests were applied across binned clade sizes. **h,** Developmental timing bias in MILK branching depth. Clade size-weighted average branching depths were calculated for each cell type in the global MILK tree and compared with the weighted average Theiler stage at which each cell type was observed. Bar colors denote Pearson correlation with developmental timing. **i,** Major cell group differences in branching depth. Cell type-specific weighted average tree depths stratified by major cell group identity.

**Fig. 4.**
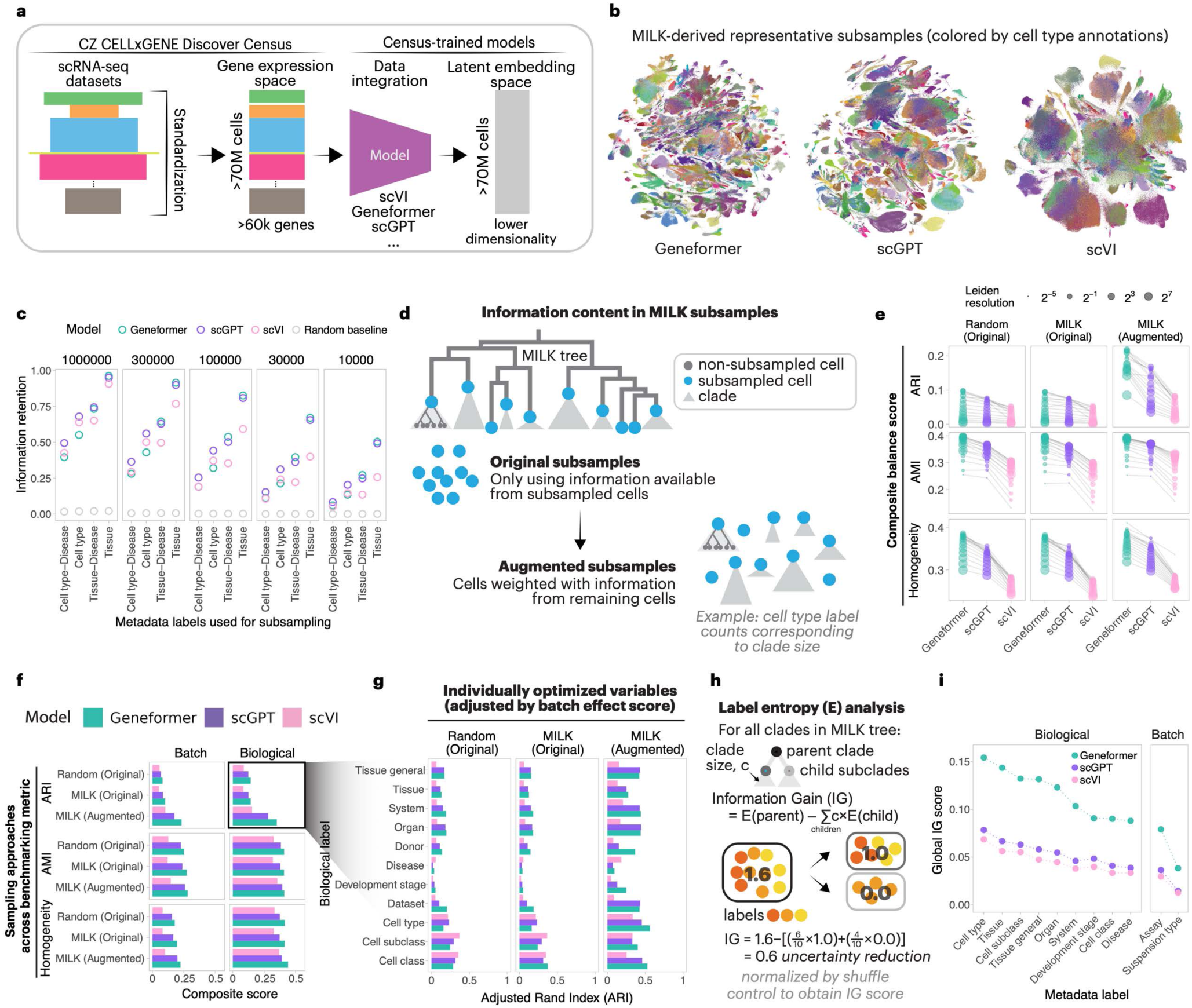
Benchmarking census-trained single-cell embeddings with MILK. **a,** CELLxGENE Census embeddings from census-trained models. Schematic overview of the CZ CELLxGENE Discover Census after processing and corresponding latent embeddings generated by Geneformer, scGPT and scVI. Geneformer and scGPT embeddings were 512-dimensional, whereas scVI embeddings were 100-dimensional. **b,** MILK-derived representative Census subsamples. MILK was applied to the latent embeddings of each model using Euclidean distance and a similarity threshold set to the 0.01 percentile. Representative subsamples were extracted at target sizes of 10,000, 30,000, 100,000, 300,000 and 1 million cells using label-balanced subsampling based on cell type–disease, cell type, tissue–disease or tissue metadata labels. UMAP embeddings of the 1 million-cell subsamples are shown for each model, colored by cell type annotation. **c,** Information retention in representative subsamples. Information retention was calculated as the fraction of cells in the complete Census represented by sampled cells and their associated downstream MILK clades. Values are shown across target sample sizes, models and metadata labels used for subsampling, together with a random-sampling baseline defined by the target sample size divided by the total number of cells. **d,** Original and augmented representative subsamples. Original subsamples use only information from sampled representative cells, whereas augmented subsamples incorporate information from downstream non-sampled cells by weighting each representative cell according to its associated clade size. **e,** Cluster-based embedding benchmark. Geneformer, scGPT and scVI embeddings were benchmarked using Leiden clustering across a grid of resolutions and compared with metadata labels using adjusted Rand index (ARI), adjusted mutual information (AMI) and homogeneity. Composite balance scores were calculated to jointly evaluate biological label separation and batch-associated clustering. Each point corresponds to a 1 million-cell subsample generated using cell type–disease label-balanced subsampling, and connecting lines indicate results obtained at the same Leiden resolution. **f,** Composite biological and batch scores. Composite scores for biological and batch-associated metadata variables after Leiden-resolution optimization, summarized across benchmarking metrics and sampling approaches. **g,** Optimized biological label scores. ARI scores for individual biological metadata variables after optimization of Leiden clustering resolution, shown across models and sampling approaches. **h,** Clade-level information gain. Schematic of entropy-based information gain (IG) calculation for a metadata label in a MILK clade. IG was calculated as the reduction in label entropy between a parent clade and its child subclades and normalized relative to a shuffled-label control. **i,** Tree-topology-based embedding benchmark. Global IG scores were computed by aggregating normalized clade-level IG values across each MILK tree using clade size as weights. Scores were globally scaled across models and metadata variables and are shown separately for biological and batch-associated labels.

**Fig. 5.**
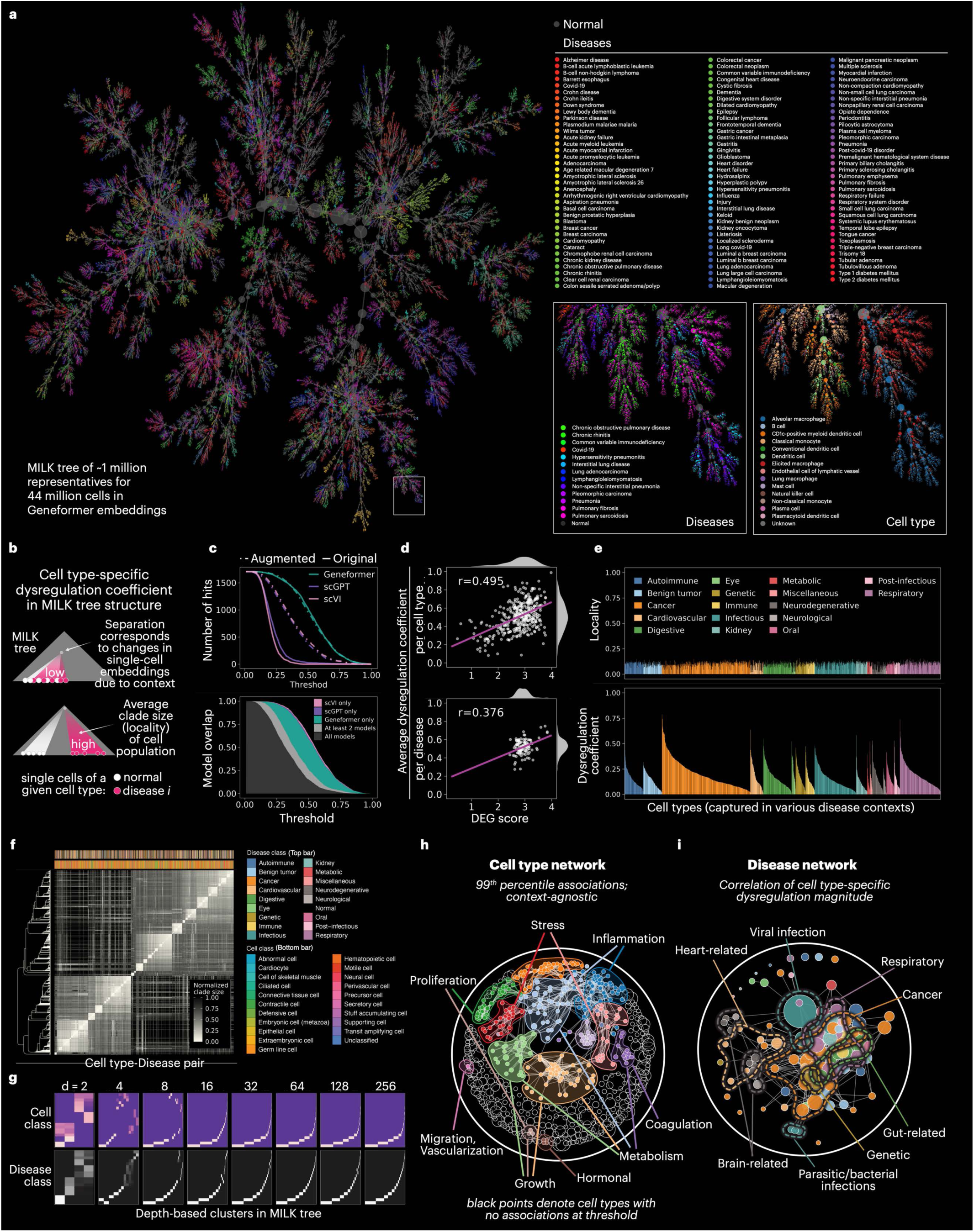
Global contextualization of disease-mediated cell-state dysregulation. MILK provides an interpretable hierarchical framework for quantifying how disease contexts reshape cell states across the CZ CELLxGENE Discover Census. **a,** Disease-context organization in the Geneformer MILK hierarchy. Force-directed layout of the Geneformer-based MILK tree reconstructed from approximately 1 million representative cells derived from the complete Census. Nodes are colored by disease context, with normal-context cells shown in dark gray. **b,** Cell type-specific dysregulation and locality metrics. Schematic of MILK-based dysregulation scoring. For each cell type–disease pair, the dysregulation coefficient was calculated as the weighted average absolute difference in clade-level relative proportions between disease-context cells and their corresponding normal-context cells across the MILK hierarchy. Locality was calculated as the characteristic clade size required to capture the corresponding cell type–disease population, with smaller values indicating stronger localization within the global hierarchy. **c,** Threshold-based dysregulation hit analysis. Cell type–disease pairs with dysregulation coefficients above a specified threshold were defined as dysregulation hits. Hit counts and model overlap were quantified across thresholds for Geneformer-, scGPT- and scVI-derived MILK trees, with and without augmentation by downstream clade information. **d,** Concordance with differential gene expression. Dysregulation coefficients derived from the Geneformer-based MILK tree were compared with the log-transformed number of differentially expressed genes between disease and normal contexts for the corresponding cell type. Values were summarized by cell type and by disease, and Pearson correlation coefficients are shown. **e,** Atlas-wide locality and dysregulation landscape. Locality and dysregulation coefficients for cell type–disease pairs derived from the Geneformer-based MILK tree, stratified by higher-level disease classification. **f,** Cell type–disease association matrix. Pairwise associations between cell type–disease populations were quantified as the weighted average clade size required to jointly capture each pair of groups within the MILK hierarchy. Smaller normalized clade sizes indicate stronger co-localization. UPGMA clustering was used to order cell type– disease pairs. **g,** Higher-order cell and disease class structure. Depth-based clusters from the UPGMA tree in **f** were evaluated for enrichment of cell class and disease class annotations. Matrices were scaled using the Sinkhorn–Knopp algorithm to visualize higher-order organization across clustering depths. **h,** Context-agnostic cell type association network. The pairwise cell type– disease matrix was aggregated into a cell type matrix by taking the median values for a pair of cell types across all contexts. An attractive and repulsive forces network layout was created, where edges were assigned for cell types sharing an association strength at the 99^th^ percentile. Community detection was applied to capture strongly associated groups of cell types, and a GSEA analysis was performed to identify underlying gene programs. Cell types with no associations were colored black. **i,** Disease association network. Disease–disease associations were inferred from Pearson correlations of dysregulation coefficients across shared cell types. Edges connect disease pairs with correlation coefficients of at least 0.5. Nodes are colored by higher-level disease classification, and manually annotated boundaries indicate major disease communities.

**Fig. 6.**
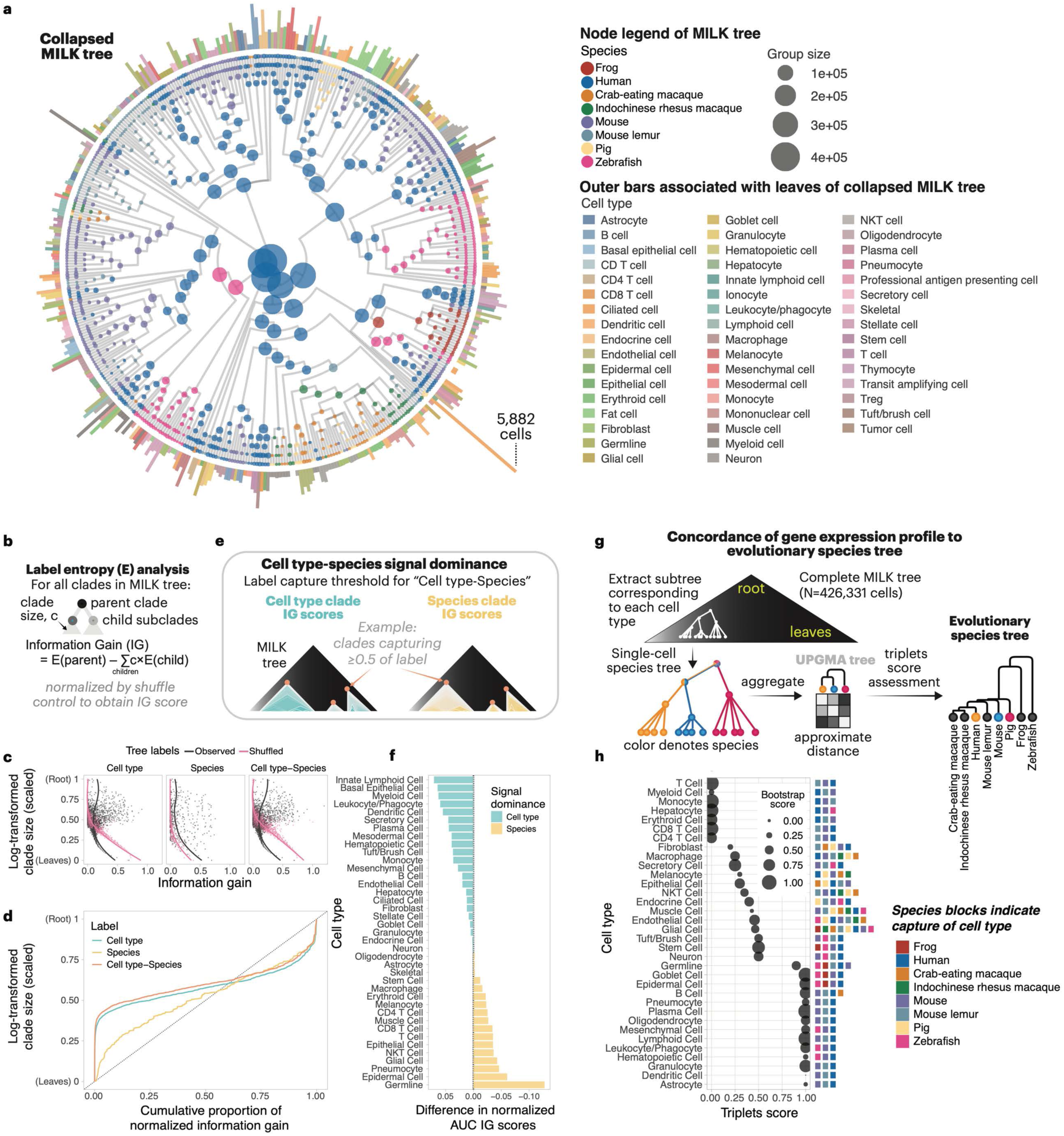
Cell type-dependent conservation and divergence of transcriptional programs across species. MILK was applied to Universal Cell Embedding (UCE) latent representations from a cross-species scRNA-seq atlas comprising approximately 3 million cells across eight species^53^: frog, human, crab-eating macaque, rhesus macaque, mouse, mouse lemur, pig and zebrafish. **a,** Collapsed cross-species MILK hierarchy. Representative MILK tree constructed from UCE embeddings using Euclidean distance and a similarity threshold set to the 0.01 percentile. The hierarchy was collapsed to preserve interpretable cell type and species label diversity. Nodes are colored by species, node size denotes group size, and outer bars indicate cell type composition associated with leaves of the collapsed MILK tree. **b,** Clade-level information gain. Entropy-based information gain (IG) was calculated for each clade with respect to cell type, species, and paired cell type–species labels. For each branching event, IG was defined as the reduction in label entropy from the parent clade to its child subclades, with child entropies weighted by clade size. IG values were normalized relative to shuffled-label controls. **c,** Information gain across clade sizes. Observed and shuffled IG values for cell type, species, and paired cell type–species labels across clades in the MILK hierarchy. Lines indicate fitted trends across clade sizes. **d,** Resolution-dependent distribution of normalized information gain. Normalized IG values were summarized as cumulative distributions across scaled log-transformed clade sizes. Smaller clades correspond to more terminal regions of the hierarchy, whereas larger clades correspond to upstream regions. **e,** Cell type–species-conditioned IG comparison. Schematic of the local signal comparison used to evaluate cell type- and species-associated organization within clades enriched for each cell type–species population. For each paired cell type–species label, clades were selected across proportional capture thresholds, and normalized IG values for cell type and species labels were averaged using paired-label purity as weights. **f,** Cell type-versus species-associated organization. Threshold-dependent IG profiles were summarized by AUC for each cell type– species label and averaged across species within each cell type. The difference between cell type-label IG AUC and species-label IG AUC is shown for each cell type, with positive values indicating stronger cell type-associated organization and negative values indicating stronger species-associated organization within clades capturing that cell type. **g,** Cell type-specific species-tree reconstruction. For each cell type, a species-specific association matrix was extracted from the MILK-derived pairwise association matrix of cell type–species groups. UPGMA clustering was then used to reconstruct a cell type-specific species tree, which was compared with the reference species phylogeny using a normalized triplets score. **h,** Phylogenetic concordance of cell type-specific species trees. Normalized triplets scores for cell type-specific species trees. Higher scores indicate stronger concordance with the reference species phylogeny. Point size denotes bootstrap confidence, calculated from 100 orthogonal bootstrap replicates, and species-colored blocks indicate the species represented for each cell type.

## RESULTS

### MILK captures the global hierarchical structure of large-scale high-dimensional objects

The core process of MILK is split into two simple steps (**Fig. 1a**). Initially, the global distribution of pairwise distances is approximated from a randomly sampled subset of objects, and an extreme-proximity threshold is defined by a percentile on the lower tail of this distribution (e.g., the 0.01 percentile). Groups of objects can then be determined in a single pass through the population by sequentially forming new groups upon encountering distinct objects according to the threshold. Specifically, the first object is automatically chosen as the representative of a new group. Each subsequent “candidate” object is then compared to existing groups for two outcomes: (i) if it is “distinct” (exceeds the threshold distance) from all representatives, it forms a representative of a new group; whereas (ii) if it is “redundant” (within the threshold distance) to any representative, it is assigned to its most similar group. Following this procedure, the representative of each group is updated to be the medoid among its group members. As such, a single cycle results in a redundancy-aware, equitable downsizing of the input population, where only cells exhibiting extremely high similarity are grouped. Recursive iteration of this procedure over representatives progressively merges groups in a bottom-up manner until a single group remains, yielding a high-resolution hierarchical representation of the complete population. Notably, although the percentile is fixed, the similarity threshold is dynamically updated across recursive cycles, reflecting the input object distribution becoming coarser across iterations. Caching group representatives is an additional feature implemented to increase the resolution of representative medoid optimization by retaining representatives from earlier, finer-resolution iterations and incorporating them throughout subsequent recursive cycles.

To enable application at scale, we implemented a recursive distributed computing strategy conceptually related to our previous large-scale lineage reconstruction framework^45^. In this implementation, input objects are partitioned into computationally tractable subsets (**Fig. 1b**), allowing the core grouping process to be performed independently and in parallel across partitions using a shared global threshold. The output consists of object groups with representative information for each partition. If the total number of groups is tractable, the groups are further merged according to their representatives using the same core grouping procedure, after which each merged group can be defined by the union of all constituent members associated with the merged representatives. Otherwise, the groups identified across partitions remain unmerged.

MILK defines similarity relative to an empirical distribution based on a user-specified distance metric, making it a generalizable and flexible method that resolves fine-grained relationships between objects with minimal parameter tuning. By recursively grouping representatives, MILK constructs hierarchical trees without explicitly computing the full pairwise distance matrix, enabling sub-quadratic scaling with respect to the number of objects. Moreover, the partitioning step allows MILK to circumvent the need to load entire datasets into memory (i.e., an “out-of-core” algorithm), ensuring scalable performance for increasingly large datasets. This contextualization of local structure within a global hierarchical tree facilitates interpretable analyses of complex systems that comprise a multitude of relationships across scales.

To demonstrate that MILK is applicable beyond genomic data, we visualized the algorithm over the MNIST dataset of handwritten digits, producing a tree representation that captures the numerical relationships between the pixelated images (**Fig. 1c**)^46^. We further demonstrated how MILK can generate subsamples from simulated evolutionary phylogenies (**Supplementary Fig. 1.1a**). Specifically, we simulated datasets composed of approximately 100,000 nucleotide sequences (1000 base pairs in length) with varying degrees of phylogenetic imbalance, which was verified using the Colless index (**Supplementary Fig. 1.1b and c**). MILK was then applied with no recursion and a similarity threshold set to 5^th^ percentile of pairwise Hamming distances, which resulted in subsamples of approximately 100 sequences (∼0.1% of the original population). These subsamples were evaluated in terms of the total branch coverage in the ground truth phylogeny (**Supplementary Fig. 1.1d and e**) and cluster coverage relative to clusters obtained from the pairwise Hamming distance matrix of the complete dataset (**Supplementary Fig. 1.1f and g**). MILK-derived subsamples exhibited significantly higher quality than random sampling across simulation replicates for both metrics (adjusted *P* values of < 0.05 and <10^−6^ for total branch coverage and cluster coverage, respectively), with effects consistently observed across all levels of topological bias. These findings provided evidence that MILK can generate representative subsamples from nucleotide sequences with diverse population distributions and no prior label information. In the following sections, we applied MILK to multiple transcriptomic atlases, demonstrating that transcriptional trees built from cell embeddings provide a general framework for single-cell analysis at unprecedented scales.

### Creating information-rich subsamples of a human fetal cell atlas of gene expression

To evaluate its ability to generate representative samples from large-scale scRNA-seq data with minimal distortion of the data structure, we applied MILK to a human fetal cell atlas of gene expression^40^ (**Fig. 2a**). We used a total of 77 cell types annotated in the original publication^40^ as biologically meaningful subpopulation markers.

MILK was independently applied to the principal component embeddings of organ datasets using a similarity threshold defined by the 0.1 percentile of pairwise cosine distances (**Supplementary Fig. 2.1a and b**). As illustrated for the eye and other representative organs, MILK identified groups that broadly spanned the distribution of UMAP embeddings of the complete datasets (**Fig. 2b** and **Supplementary Fig. 2.1c**), while maintaining within-group specificity comparable to nearest neighbors for earlier recursive iterations (**Supplementary Fig. 2.2a and b**). Across iterations, MILK progressively captured the data landscape from fine to global resolution (**Supplementary Fig. 2.1d**), as reflected by the qualitative clustering of cell type information in the MILK tree hierarchies (**Fig. 2b** and **Supplementary Figs. 2.1e, 2.3, and 2.4**). We further measured average cell type agreement, cosine distance, and specificity of group members to their respective representatives, together with the number of groups across the recursive MILK iterations for all organs (**Fig. 2c**). These results showed robust preservation of cell type information for dataset compression levels up to around 1,000-fold. Collectively, these analyses demonstrated that early MILK iterations resolve locally distinct subpopulations, whereas later iterations captured broader patterns as subpopulations merged to produce global trends.

As datasets continue to grow, downsampling has become a practical necessity for tractable downstream analyses. Extracting subsamples from a MILK tree can be formulated as selecting a set of non-overlapping (internal) nodes that maximizes leaf coverage while balancing representation across subpopulations. We developed both label-free and label-balanced strategies to subsample from MILK trees, the latter of which leveraged cell type annotations during selection (**Fig. 2d** and **Supplementary Fig. 2.5**). The label-free strategy selects non-overlapping clades from the MILK hierarchy without using annotations, prioritizing structurally distinct regions of the tree until the target subsample size is reached. In contrast, the label-balanced strategy uses pre-existing annotations to merge and select only label-consistent clades, while explicitly selecting an approximately balanced representation across cell types while respecting the MILK topology. Motivated by the observations that MILK-defined groups overlap with nearest neighbors, we additionally implemented pseudobulking of the raw gene counts within groups to form “metacells”, which has been shown to mitigate technical dropout and stabilize biological signal^47^ (**Supplementary Fig. 2.6**).

To assess the extent to which cell type-specific signals were preserved following downsampling, we generated subsamples of varying sizes from the MILK tree under combinations of label usage during subsampling (label-free or label-balanced) and with and without downstream aggregation into metacells. Given the sparsity and noise inherent to expression counts per individual gene, we quantified the preservation of biological signal across a comprehensive list of literature-curated transcription factor-target gene sets, or “regulon targets”^48^. Specifically, for each cell type, we measured the Spearman correlation between complete and downsampled gene set profiles, which were defined as vectors of activity scores (average expression of regulon targets minus background expression) across all regulons in the database. We then summarized the performance of different strategies as the area under the curve (AUC) of correlation versus log-transformed downsampling proportion per cell type (**Fig. 2e**). Label-free MILK subsamples exhibited greater correlation compared to their respective random subsampling baselines (AUC values of 2.83 and 2.49 for MILK and random, respectively). On the other hand, label-balanced subsamples showed similar performance between random and MILK (2.59 for both), which was expected given that much of the work to define robust subpopulations was carried out during the precursor cell type annotation task. Pseudobulking further increased correlation scores relative to single-cell counterparts (from 2.83 to 3.27 for label-free MILK; from 2.59 to 3.02 for label-balanced MILK), indicating robust retention of regulon signals.

We further investigated the preservation of transcriptomic covariance structure following downsampling. For each cell type, we compared the dominant regulon-specific principal component subspaces between complete and subsampled datasets by measuring the average cosine similarity of corresponding subspace angles (**Fig. 2f**). Similar to the Spearman correlation analysis, label-free MILK subsamples showed marked improvement (AUC of 4.07) compared to their random baseline (3.67), whereas label-balanced MILK subsamples exhibited a marginal improvement (4.32 and 4.31 for MILK and random, respectively). Metacell aggregation further improved cosine similarity relative to single-cell counterparts (from 4.07 to 4.37 for label-free MILK; from 4.32 to 4.59 for label-balanced MILK). Taken together, these results indicate improved retention of regulon-specific transcriptomic variation within cell type groups for MILK-derived subsamples compared to random downsampling conventions.

We next evaluated the downsampling strategies in terms of constructing integrated atlases with scVI (single-cell Variational Inference), a generative model that learns latent representations of single-cell transcriptomes while correcting technical variation such as batch effects^34^. Given the independently downsampled organ datasets generated across varying target sizes and subsampling strategies, we trained scVI models to construct integrated (reference) human fetal cell atlases (**Fig. 2g** and **Supplementary Fig. 2.7a**). Preliminary findings showed that MILK-derived atlases led to improved cell type clustering compared to random baselines, particularly for smaller target sizes (**Supplementary Fig. 2.7b**). To further assess the quality of the latent representations, we used scArches^35^ to map all remaining (query) cells from each organ onto the reference atlases. We then measured the average nearest-neighbor (*k* = 8) distance of each query cell to reference cells of the same cell type (**Fig. 2h** and **Supplementary Fig. 2.8**), where a smaller distance corresponds to a higher-quality reference atlas. MILK-derived atlases showed improved performance relative to random sampling counterparts, especially for greater levels of downsampling. These results indicate that MILK improves the retention of population structure during data compression, suggesting that structurally representative subsampling could reduce the data scale required for training single-cell reference and foundation models while preserving downstream performance.

Altogether, we established MILK as a scalable, cost-effective framework (**Supplementary Fig. 2.1f**) for generating representative subsamples that retained biologically relevant signal in a human fetal cell atlas. MILK consistently improved performance both with and without the use of cell type labels during subsampling, highlighting its viability as a label-agnostic method. We also demonstrated that generating metacells according to the MILK tree structure could further improve subsample quality, which provides a scalable, structured, and data-driven approach for retaining information beyond the set of subsampled cells.

### MILK tree structure captures single-cell trajectories in a mouse developmental atlas

To explore whether MILK trees capture dynamic temporal information, we applied it to a transcriptomic atlas of prenatal mouse development containing over 11 million processed cells^19^. Comprised of 83 mouse embryos, the atlas spanned 16 Theiler stages from late gastrulation through birth, as defined by morphological and anatomical features^49^ (**Fig. 3a**). For subsequent analyses, we retained cell type and major cell group annotations provided in the original publication.

For each Theiler stage dataset, MILK was applied to the principal component embeddings (100 components) using cosine distance and a similarity threshold set at the 0.1 percentile. The label-balanced sampling strategy described in **Fig. 2d** was then used with cell type annotations to extract representative subsamples from the resulting Theiler stage MILK trees at varying target sizes. These subsamples were concatenated to construct cell type- and Theiler stage-balanced atlases of mouse development. Notably, we did not perform batch normalization because embryo identity was strongly confounded with Theiler stage signal (**Supplementary Fig. 3.1a and b**), suggesting that correcting embryo-specific batch effects would likely collapse temporal variation. MILK was subsequently applied using the same parameterization to generate global developmental atlas trees, in which robust, cell type-specific Theiler stage-associated gradients were observed across many branching clades (**Fig. 3b** and **Supplementary Fig. 3.1c and d**).

We compared the global MILK trees with established trajectory inference methods to assess their ability to recapitulate developmental progression of gene expression programs. Specifically, for pairs of cells, the absolute difference in Theiler stage was calculated as a function of the following trajectory inference measures: (i) MILK tree distance; (ii) Diffusion Pseudotime (DPT) value^50^ (**Supplementary Fig. 3.2**); and, (iii) the expected transport mass inferred by Waddington-OT (WOT)^51^ (**Fig. 3c**). To enable direct comparison, the complement of WOT transport mass was utilized and each trajectory measure was aggregated into ranked bins. Notably, whereas MILK enabled unrestricted measurement of distances between any pair of cells, DPT- and WOT-based analyses required additional constraints. For pseudotime inference, comparisons were restricted to cell pairs within major cell groups to ensure graph connectivity, with one source cell randomly selected per group. In contrast, WOT explicitly leveraged Theiler stage information by design, with distance measurements restricted to cell pairs across Theiler stages. While all methods captured similar trends at short trajectory distances, DPT and WOT were less effective at capturing long-range relationships, as reflected by a decrease in Theiler stage differences at larger trajectory distances. In contrast, MILK tree distances remained positively correlated with developmental progression across the full trajectory inference spectrum, demonstrating MILK’s ability to encode both local and global temporal dynamics.

Given the complexity and diversity of cell populations captured in large-scale atlases, we next assessed whether the cell type relationships encoded in the MILK tree structure were consistent with cell type trajectory graphs inferred by partition-based graph abstraction (PAGA)^52^. Each clade in the MILK tree was decomposed into a cell type frequency profile, and its agreement with the PAGA adjacency matrix of cell type connectivity was quantified using the quadratic form equation (**Fig. 3d**). Score distributions were evaluated for Theiler stage-specific and global developmental atlas MILK trees, with the results exhibiting significantly higher agreement scores relative to a control MILK tree with shuffled cell type labels (adjusted *P* value ≍ 0) (**Fig. 3e** and **Supplementary Fig. 3.3**). These results suggested that MILK tree structure meaningfully encodes complex cell type-specific relationships at scale.

To holistically assess the hierarchical organization encoded by MILK, we further evaluated whether MILK trees captured cell differentiation trajectories during mouse embryogenesis, as proposed in the original publication^19^ (**Fig. 3f**). Specifically, to compare the single-cell MILK tree with their curated cell type tree (i.e., treating it as a ground truth), each clade was scored using an asymmetric weighted Jaccard index (**Supplementary Fig. 3.4**). To control for baseline similarity, scores were normalized against clades derived from a cell type-shuffled negative control tree, yielding a fold-change value for each clade (**Fig. 3g**). As expected under this evaluation framework, both small and large clades exhibited minimal fold change, reflecting limited information content at the level of individual cells (leaf nodes) and the near-complete mixing of cell types toward the root. In contrast, intermediate clades showed significantly higher fold changes across binned clade sizes ranging from 31.86 to 74,202.15 cells. These observations indicated that MILK detects biologically coherent cell type relationships across mouse development at intermediate resolutions that balance specificity and stability.

These observations collectively motivated the hypothesis that the global MILK developmental atlas may capture the organization of embryogenesis, from the fertilized egg to later developmental stages, through patterns of cell type diversification. To examine this hypothesis, we computed the weighted-average branching depths of cell types in the MILK tree from the single root and compared these values to the weighted-average Theiler stage at which the respective cell types were observed. Despite substantial diversity in cell types, many of which span broad developmental windows, we observed a moderate correlation (r = 0.533), where cell types predominantly present in earlier Theiler stages tended to branch closer to the root in the MILK tree (**Fig. 3h** and **Supplementary Fig. 3.5a and b**). Further stratification by major cell groups revealed higher-order biological structure captured by MILK, with mesoderm and primitive erythroid populations branching closer to the root (smaller depth), whereas cardiomyocytes and testes/adrenal lineages were preferentially localized towards the leaves (greater depth) of the MILK tree (**Fig. 3i** and **Supplementary Fig. 3.5c**).

In summary, applying MILK to a mouse developmental atlas demonstrated that the hierarchical structure inferred solely from gene expression data can encode systems-level lineage relationships of cell differentiation across mouse development. By representing single-cell relationships through high-resolution pairwise hierarchies, MILK scales with increasing biological complexity, such as the number and diversity of cell types, while remaining interpretable across multiple resolutions. Together, these results suggest that MILK is well-suited for the analysis of large-scale single-cell atlases, where conservation of both global organization and fine-grained structure remains essential.

### Benchmarking foundation-model embeddings of the CZ CELLxGENE Discover Census

The emergence of large-scale consortia initiatives to compile and standardize single-cell genomic data creates an opportunity to investigate cellular states across diverse contexts. To analyze such resources at scale, foundation models and deep generative frameworks have emerged as powerful approaches for integrating single-cell transcriptomic data into shared latent spaces. These methods aim to capture biologically meaningful transcriptional structure while minimizing technical variation, such as batch effects stemming from differences in experimental protocols, sequencing technologies, and sample processing. One such resource is the CZ CELLxGENE Discover Census^7^, together with single-cell latent embeddings derived from several census-trained models hosted by CZI, including Geneformer, scGPT, and scVI, facilitating large-scale comparative analyses across diverse biological contexts (**Fig. 4a** and **Supplementary Fig. 4.1a**).

While these models have shown substantial promise in integrating heterogeneous transcriptomic datasets, a remaining challenge is evaluating the extent to which their latent embeddings preserve biologically meaningful structure while correcting for technical variation for tens to hundreds of millions of cells. In particular, aggressive normalization or integration may inadvertently obscure meaningful biological distinctions, whereas insufficient correction can result in embeddings dominated by batch-specific structure. Existing benchmarking approaches are often limited by computational tractability, sensitivity to arbitrary parameter choices, and difficulties in capturing the multi-scale organization inherent to biological systems.

Because MILK enables scalable hierarchical analysis of extremely large cell populations while retaining multi-resolution structure, we reasoned that it could provide a framework for systematically benchmarking latent embeddings derived from census-trained models. Using MILK, we analyzed the global organization of single-cell embeddings for approximately 44 million cells from primary studies in the CELLxGENE Discover Census (Version 2024-07-01) generated by two foundation models, Geneformer (512 dimensions) and scGPT (512 dimensions), as well as scVI (100 dimensions), a probabilistic deep generative model.

For each embedding, we constructed a MILK tree using Euclidean distance as the pairwise metric and a similarity threshold fixed to the 0.01 percentile of the input objects in respective MILK iterations. Across all models and metadata variables examined, label diversity remained broadly preserved until the number of representative cells decreased below approximately one million cells (**Supplementary Fig. 4.1b**). We then extracted representative subsamples of varying target sizes from the resulting MILK trees using the label-balanced subsampling strategy according to different metadata categories: “cell type” (**Fig. 4b**), “cell type–disease”, “tissue”, and “tissue–disease” (**Supplementary Fig. 4.2**).

We next quantified the degree of information retained by representative subsamples relative to the complete census by measuring the concordance between MILK tree topology and the underlying metadata annotations. For each metadata-guided subsampling strategy, information retention was calculated as the fraction of cells in the complete dataset represented by the sampled cells and their associated downstream MILK clades. Across target sizes and models, we observed a progressive reduction in retained information as the metadata labels used in the label-balanced subsampling process became more granular, consistent with finer-resolution biological distinctions exhibiting reduced overlap with the global hierarchy (**Fig. 4c**). Information retention also decreased systematically as target subsample size decreased, with representative subsets of one million cells striking a practical balance between computational tractability and preservation of the original dataset structure.

Because each representative cell in a MILK tree represents a clade of structurally related cells, MILK-derived subsamples can retain additional contextual information beyond the sampled cells themselves. We therefore investigated whether incorporating such additional information could improve downstream analyses relative to using the representative cells alone (i.e., “augmented” subsample vs “original” subsamples). Unlike pseudobulking approaches that explicitly aggregate molecular profiles, this strategy preserves the original latent embeddings of representative cells while incorporating higher-order contextual information in a computationally tractable manner. As a proof-of-principle, we propagated information from non-subsampled cells to each representative cell by weighting its contribution by the number of downstream members in its clade, allowing representative cells to implicitly summarize the broader populations they represent (**Fig. 4d**).

Using the augmented and original MILK subsamples derived from the CELLxGENE Discover Census, we benchmarked latent embeddings generated by Geneformer, scGPT, and scVI models according to their ability to discriminate biologically meaningful metadata label structure while minimizing technical variation. Specifically, we performed Leiden clustering across a broad range of clustering resolutions, and then evaluated the resulting clusters relative to manually-curated biological and batch metadata variables in terms of the Adjusted Rand Index (ARI), Adjusted Mutual Information (AMI), and Homogeneity metrics (**Supplementary Fig. 4.3a and b** and **Supplementary Fig. 4.4**).

To compare model and subsampling performance across clustering resolutions, we first calculated composite biological and batch scores as the geometric mean across their respective metadata variables (**Supplementary Fig. 4.3c and d**). A composite balance score was then defined as the product of the composite biological score and the complement of the composite batch score (**Fig. 4e** and **Supplementary Fig. 4.3e**). Across clustering resolutions, an ANOVA analysis indicated model identity as a statistically significant determinant of benchmarking performance (ARI: *F* = 44.81, *P* = 1.19×10^-15^; AMI: *F* = 52.87, *P* < 2×10^-16^; Homogeneity: *F* = 246.01, *P* < 2×10^-16^). Subsampling strategy also significantly contributed to observed ARI (*F* = 88.45, *P* < 2×10^-16^) and Homogeneity (*F* = 5.24, *P* < 0.01) scores. Notably, a significant interaction between model and subsampling strategy was observed only for ARI (*F* = 12.84, *P* = 6.99×10^-9^), indicating that the impact of subsampling approach on cluster-label agreement varied across models. *Post hoc* paired *t*-tests with Bonferroni correction revealed significant pairwise differences between all models and across all benchmarking metrics (adjusted *P* < 10^-6^ in all cases). Geneformer consistently achieved the highest scores, followed by scGPT and scVI across all three metrics (**Supplementary Fig. 4.4a** and **b**). Similarly, augmented MILK subsamples significantly outperformed both original MILK and random subsamples for the ARI metric (adjusted *P* < 10^-12^) (**Supplementary Fig. 4.4a**), whereas no significant differences were observed between subsampling approaches for AMI or Homogeneity.

Because different metadata label types are optimally resolved at different clustering granularities (**Supplementary Fig. 4.3e**), we subsequently normalized biological variables by the composite batch score at each corresponding clustering resolution. We then independently selected the Leiden clustering resolution yielding the maximum batch-adjusted biological score for each variable. The resulting scores were further aggregated using the geometric mean to obtain a normalized composite biological score, while the corresponding batch scores at the selected resolutions were retained to assess the associated batch structure (**Fig. 4f** and **Supplementary Fig. 4.3f**). Across all metrics, Geneformer-derived embeddings consistently exhibited the strongest preservation of batch-adjusted biological label structure, followed by scGPT and scVI. However, stronger preservation of biological signal was accompanied by increased retention of batch-associated structure, highlighting an inherent tension between biological discriminability and technical harmonization in atlas-scale integration of single-cell transcriptomic data. This trade-off was particularly evident for high-resolution biological variables (**Fig. 4g**), where embeddings that strongly separated fine-grained cellular states often exhibited reduced attenuation of assay- or protocol-associated variation (**Supplementary Fig. 4.3a and b**).

Despite extensive optimization of Leiden clustering resolutions for individual metadata labels and benchmarking metrics, we observed substantial variability in benchmarking scores across metadata variables (**Fig. 4g**), highlighting a broader limitation of conventional label-based benchmarking approaches performed directly on latent embeddings. In particular, different biological variables were optimally resolved at different clustering granularities, complicating direct comparison across resolutions and metadata types. Given the concordance between the MILK tree topology and biological metadata structure, we reasoned that tree topology-based metrics could provide a more coherent framework for evaluating models in the context of complex, multi-resolution biological structure.

To address this, we employed information gain (IG) as an additional benchmarking metric to quantify how effectively biological information was encoded within the MILK tree topology (**Fig. 4h**). Unlike cluster-based benchmarking approaches, IG-based analyses do not require explicit resolution parameterization, therefore enabling direct comparison of biological variables across multiple scales. Upon summarization of normalized IG values into MILK tree-level scores, we observed that the Geneformer-based MILK tree with augmented information retained the strongest biological signal, followed by scGPT and scVI (**Supplementary Fig. 4.5a–c**). Ranking biological label variables by their IG-based scores recapitulated a hierarchical progression of biological organization, ranging from broad disease- and cell-class-level structure to increasingly specific cell type- and tissue-level information (**Fig. 4i**). Consistent with the Leiden clustering analyses, Geneformer-based MILK trees exhibited the highest IG values for batch-associated labels, while scGPT and scVI showed comparatively minimal batch-associated structure. However, focusing on the absolute difference in model performance per metadata variable (ΔIG) revealed that the relative gain in biological signal substantially exceeded that of batch signal for Geneformer compared to scGPT and scVI (**Supplementary Fig. 4.5b**), indicating that its latent embeddings achieve the most favorable balance between preservation of biological structure and suppression of technical variation among the models examined.

In summary, applying MILK to the CELLxGENE Discover Census established a scalable framework for constructing representative and hierarchically contextualized views of atlas-scale single-cell embeddings. Through representative subsampling and hierarchical information retention, MILK enabled tractable benchmarking analyses over approximately 44 million cells spanning diverse cellular contexts. Applying clustering and tree topology-based benchmarking strategies, we systematically evaluated how different models balance preservation of biological organization against suppression of technical variation across multiple biological resolutions. Moreover, augmentation of representative subsamples using contextual information encoded in the MILK tree structure reproducibly improved benchmarking performance, demonstrating that biologically relevant information remains captured within the topological tree structure following substantial data compression (approximately 97.7% compression). These findings suggest that hierarchical contextualization with MILK provides a scalable and information-rich framework for evaluating increasingly large and heterogeneous single-cell genomic resources.

### Global contextualization of disease-mediated cell-state dysregulation reveals emergent cellular and disease associations

Building upon the global hierarchical contextualization of the CELLxGENE Discover Census enabled by MILK, we next investigated how disease context alters cell states within the unified hierarchy (**Fig. 5a**). Using the MILK trees with approximately one million representative cells derived from label-balanced subsampling using cell type–disease label information, we quantified, for each cell type–disease pairing, the extent to which cells in a disease context diverged from their corresponding “normal” state across the global hierarchy. Specifically, we defined a dysregulation coefficient as the weighted average of the absolute differences between the relative proportions of normal- and disease-context cells across clades in the MILK tree (**Fig. 5b** and **Supplementary Fig. 5.1a**). In parallel, we quantified the locality of each cell type–disease group as the characteristic clade size required to robustly capture the subpopulation, with smaller values indicating stronger localization within the global MILK hierarchy (**Supplementary Fig. 5.1b**).

Using the dysregulation coefficients derived from Geneformer-, scGPT-, and scVI-based MILK trees, we identified dysregulated cell types in disease contexts (“dysregulation hits”) and assessed their overlap across models. Dysregulation hits were defined as cell type–disease labels with dysregulation coefficients exceeding a specified threshold, enabling systematic comparison of disease-associated cellular perturbations detected by each model. Across a range of thresholds, Geneformer-based MILK trees consistently identified the largest number of dysregulation hits and recovered the majority of hits detected by scGPT- and scVI-based MILK trees (**Fig. 5c**). Augmentation of the one million representatives with their downstream clade size information (**Fig. 4d**) for the calculation of dysregulation coefficients resulted in a moderate increase in the number of hits for scGPT and scVI-based MILK trees; however, Geneformer-based MILK trees continued to capture the largest set of dysregulated cell type–disease relationships regardless of augmentation state, further supporting the greater sensitivity of Geneformer in capturing the diverse spectrum of disease-associated cell-state shifts within the global MILK hierarchy.

To assess whether MILK trees preserve disease-associated cellular perturbations identified in local tissue contexts, we applied MILK to 54 tissue-specific datasets extracted from the CELLxGENE Discover Census and independently calculated dysregulation coefficients for their respective cell type–disease subpopulations (**Supplementary Fig. 5.2a**). For comparison, we included an alternative clustering-based approach, in which dysregulation coefficients were computed across Leiden clusters generated at multiple clustering resolutions, rather than across MILK clades (**Supplementary Fig. 5.2b**). Due to memory constraints associated with large-scale Leiden clustering, label-balanced subsamples (target size of 1 million cells) of tissue datasets obtained from MILK trees were analyzed to enable evaluation across the full atlas-scale landscape. The resulting dysregulation coefficients from the global MILK hierarchy exhibited moderate, model-dependent correlation with the dysregulation measurements from local tissue MILK hierarchies and were comparable to those observed with the Leiden clustering-based approach (**Supplementary Fig. 5.2c**). Notably, the augmented global MILK hierarchy captured substantially more context-independent (global-specific) dysregulation hits while simultaneously minimizing the number of tissue-specific hits compared to the Leiden-clustering approach (**Supplementary Fig. 5.2d and e**). These results suggest that the global MILK hierarchy retains biologically meaningful disease-associated structure despite substantial data compression, and that contextualizing the complete single-cell landscape can uncover underappreciated gene expression associations that span biological contexts.

We subsequently investigated whether MILK-based dysregulation coefficients reflect underlying transcriptional changes by quantifying the number of differentially expressed genes (DEGs) from the corresponding non-integrated expression profiles for each cell type–disease pairing relative to its normal context (**Fig. 5d**). Aggregating dysregulation coefficients per cell type and disease, respectively, we observed a moderate positive correlation with the log-transformed number of DEGs, with correlation coefficients of 0.495 for cell type and 0.376 for disease.

Locality and dysregulation coefficients for cell type–disease pairings revealed substantial variability in the magnitude of disease-associated cellular state perturbations across higher-order disease classifications (**Fig. 5e** and **Supplementary Fig. 5.3**). Despite this variability, most cell type–disease populations were localized to relatively restricted regions of the global hierarchy, with a median normalized locality of 0.11 (**Fig. 5e** and **Supplementary Fig. 5.1b**). This observation suggests that disease-associated cellular state labels frequently occupy coherent transcriptomic niches within the global MILK hierarchy rather than being broadly dispersed across the cellular landscape.

We therefore investigated whether these dysregulation coefficients could identify cell type–disease pairings associated with meaningful transcriptional divergence. Specifically, we selected representative cell type–disease examples exhibiting high (fibroblast–Plasmodium malariae malaria), intermediate (neuron–Alzheimer’s disease), and low (medium spiny neuron–opiate dependence) dysregulation coefficients and identified MILK clades capturing the corresponding cell populations in both disease and normal contexts (**Supplementary Figs. 5.4 and 5.5**). Cell type–disease pairings with high and intermediate dysregulation coefficients exhibited pronounced disease-associated transcriptional programs, whereas pairings with low dysregulation coefficients showed comparatively limited divergence. These results provided strong preliminary evidence that the MILK hierarchy can prioritize cell type– disease pairings exhibiting substantial disease-associated remodeling of cellular state, thereby providing a scalable framework for identifying biologically meaningful perturbations within atlas-scale single-cell resources.

To comprehensively characterize the global landscape of the normal cellular context space together with the disease-mediated dysregulation of cell types, we first quantified pairwise associations between cell type–disease (including “normal”) subpopulations using the MILK hierarchy. For each pair, we estimated the characteristic scale of the MILK hierarchy at which the two cell type–disease populations co-localized by calculating the weighted average size of clades that jointly captured both groups. Smaller clade sizes, therefore, indicate stronger associations, indicating that the two populations occupy nearby regions of the global cellular hierarchy. Taking the association profiles, we then hierarchically clustered the cell type– disease populations (**Fig. 5f**). Although the association matrix was constructed from fine-grained cell type–disease pairs, this clustering revealed higher-order structure when the paired labels were decomposed into distinct “cell class” and “disease class” components (**Fig. 5g** and **Supplementary Fig. 5.6**). In other words, related cell and disease classes tended to co-organize within the hierarchy, even though the clustering was performed on paired cell type–disease labels rather than on either annotation alone. These results suggest that disease-mediated cell-state shifts are not randomly distributed across the atlas but instead converge into recurring cellular and pathological modules.

We next asked whether cell types commonly dysregulated across distinct disease contexts converge to dysregulation of shared cellular programs. To this end, we aggregated the cell type–disease matrix according to the median normalized clade size for each pair of cell types across all disease (and normal) contexts (**Supplementary Fig. 5.7a**). From this cell type matrix, we applied an association-strength threshold at the 99^th^ percentile to retain only high-association signals between cell types (**Supplementary Fig. 5.7b**). This produced a cell type adjacency matrix comprising 630 cell types and 1,831 high-confidence associations, from which Leiden community detection identified 13 groups containing at least two associated cell types (**Fig. 5h** and **Supplementary Fig. 5.7c**). To interpret the biological structure of these associated cell type groups, we performed gene set enrichment analysis (GSEA) on each Leiden community (**Supplementary Fig. 5.7d–f**). We identified significant enrichment for terms in 11 of the 13 groups, revealing transcriptional programs associated with core physiological response modules, including proliferation, stress, inflammation, metabolism, and related cellular processes. Thus, although the associated groups often comprised phenotypically diverse cell types, their organization was supported by shared gene expression programs. These results suggest that disease-contextualized cell states can converge onto conserved transcriptional modules across otherwise distinct cellular identities, highlighting how MILK can identify emergent molecular associations shaped by both cell identity and disease context.

Similarly, we also explored whether diseases could be related to one another through shared patterns of cell-state dysregulation. We constructed a disease association network across 105 disease contexts, excluding three diseases with fewer than 50 cells, by calculating the Pearson correlation of dysregulation coefficients across shared cell types for each pair of diseases (**Supplementary Fig. 5.8**). Disease pairs with correlation coefficients of at least 0.5 were connected by edges, producing a network that captures coordinated shifts in cell states across distinct pathological contexts (**Fig. 5i**). When nodes were annotated by higher-order disease classifications, related disease classes formed coherent communities. A prominent cluster spanning cancer, benign tumors, genetic, autoimmune, respiratory, and gut-related diseases dominated the landscape, likely reflecting recurrent dysregulation or activation of immune and stromal cell states across these contexts. We also observed partial overlap between heart- and brain-related diseases, suggesting shared perturbation patterns distinct from other groups. In contrast, viral and parasitic/bacterial infectious diseases were largely separated, although viral respiratory diseases, including influenza and COVID-19, remained strongly linked to respiratory diseases. These results suggest that MILK-derived dysregulation profiles can organize diseases according to coordinated cellular perturbations, revealing relationships that may bridge clinically distinct pathological contexts through shared cell-state programs.

Importantly, the standardization of CELLxGENE Discover Census annotations represents a substantial collective effort by the scientific community to harmonize single-cell information across many independent studies. While these annotations provide a shared basis to conduct atlas-scale analyses, the discrepancies in annotation strategies, nomenclature, and cell-state resolution can influence the interpretation of cell type-specific patterns and remain difficult to quantify. We therefore leveraged the data-driven MILK tree structure to evaluate the extent to which these annotations reflect robust cellular states. Initially, we examined annotation consistency solely within normal cellular contexts by quantifying cell type purity in the MILK tree (**Supplementary Fig. 5.9a)**. While a majority of cell types exhibited coherent localization within the hierarchy, others displayed reduced purity scores across clades (**Supplementary Fig. 5.9b**). Extending this analysis to pairwise relationships between cell types revealed a small number of cell type groups exhibiting co-localized purity scores throughout the MILK hierarchy (**Supplementary Fig. 5.9c–f**). While such patterns may reflect a biological continuum of cell states driven by shared transcriptional programs, they may alternatively highlight instances where the annotations do not align with the global organization of cellular states and therefore represent candidates for future refinement using MILK.

We additionally investigated whether cell type–disease subpopulations can occupy cellular states that are not captured by the normal-context cellular landscape. For each group, we quantified whether its localization differed from all normal-context cells by calculating the difference in purity scores across MILK clades (**Supplementary Fig. 5.10a**). Summarizing these differences across the hierarchy, 7% (155/2129) of cell type–disease subpopulations exhibited preferential localization outside of the regions occupied by normal cellular states (**Supplementary Fig. 5.10b**). In contrast, the majority (93%) of groups remained localized within regions represented by normal cell state diversity, suggesting that most disease perturbations reflect modifications of existing cellular transcriptional programs rather than distinct cellular states.

In summary, these analyses demonstrate that the MILK tree structure can contextualize disease-mediated cell-state dysregulation across atlas-scale single-cell landscapes. By quantifying how disease-context cells redistribute relative to their normal counterparts and across diseases, MILK identified that cells dysregulated by diverse diseases tend to shift their transcriptional states toward other states present in normal bodies, and that their transition patterns are shared across diseases. As such, MILK extends beyond detecting isolated disease effects, providing a scalable approach to uncover coordinated cellular programs and generate testable hypotheses about common pathological mechanisms across diverse disease contexts, while supporting both annotation refinement and potential discovery of emergent disease-associated cell states.

### MILK reveals cell type-dependent conservation and divergence of transcriptional programs across species

The molecular states of cells can diverge across species through evolutionary change, yet cellular identity is also shaped by intrinsic developmental and physiological programs. In multicellular organisms, the interplay of gene regulatory networks, lineage constraints, and cell–cell interactions drive cellular trajectories through state spaces often conceptualized as Waddington landscapes. Under this view, ancestral or progenitor-like programs may retain relationships that broadly follow species phylogeny, whereas more differentiated or specialized cell states may be remodeled within each lineage, producing transcriptional relationships that deviate from canonical evolutionary trajectories. Disentangling these two axes of organization, phylogenetic divergence and cell type-specific state diversification, remains a central challenge in evolutionary and comparative single-cell biology, particularly as cross-species atlases continue to expand in scale and complexity.

Because MILK organizes large numbers of single-cell transcriptomic profiles into an interpretable hierarchy, we reasoned that it could provide a framework for interrogating conservation and divergence of cellular programs across species. We therefore applied MILK to latent embeddings from a publicly available subset of a cross-species scRNA-seq atlas generated with the Universal Cell Embeddings (UCE) foundation model^53^. This dataset comprised 49 annotated cell types across eight species (frog, human, crab-eating macaque, Indochinese rhesus macaque, mouse, mouse lemur, pig, and zebrafish), providing a unified embedding space of approximately 3 million cells in which cell type and species relationships could be jointly examined (**Supplementary Fig. 6.1**). MILK was applied using Euclidean distance as the pairwise metric and a similarity threshold fixed to the 0.01 percentile, and label-balanced subsampling was performed to obtain a representative hierarchy of cell type and species diversity containing approximately 400,000 cells. Visualization of the MILK tree revealed broad interdigitation of cell type and species labels throughout the hierarchy with substantial variability in cell type–species group locality (**Supplementary Fig. 6.2**), suggesting that the UCE embedding captures both conserved cellular identity and species-specific transcriptional variation within a shared representation (**Fig. 6a**).

To systematically analyze cell type and species organization within the MILK tree topology, we conducted an entropy-based analysis in which information gain (IG) was independently calculated for cell type, species, and paired cell type–species labels across all clades in the MILK tree (**Fig. 6b**). Following per-clade normalization relative to a shuffle control IG baseline (**Fig. 6c**), the cumulative distribution of IG revealed distinct organizational patterns, with species signal showing relative enrichment toward terminal branches, whereas cell type and cell type–species IG signal was predominantly organized by upstream clades (**Fig. 6d**). This organization supports a model in which conserved cell type programs define the broad architecture of the cross-species transcriptomic landscape, while species-specific regulatory divergence further refines cellular states through evolution.

To further investigate these patterns at the level of individual cell types, we calculated the weighted average species IG and cell type IG across clades in which the corresponding cell type–species label exceeded a specified proportional threshold, using clade purity as weights (**Fig. 6e**). Summarizing these IG profiles across different proportional thresholds for each cell type–species label by AUC and averaging the AUC values across species, we compared the relative contribution of cell type and species organization for each cell type captured by the MILK tree. A higher cell type AUC (positive difference) indicates that clades enriched for a given cell type primarily capture cell type-specific structure, whereas a higher species AUC (negative difference) suggests that its clades preferentially capture species-specific variation. Ranking cell types in terms of their difference between cell type and species AUCs revealed substantial heterogeneity in signal dominance across cell lineages (**Fig. 6f**). Notably, adaptive immune populations, including “CD4 T cell,” “CD8 T cell,” “T cell,” and “NKT cell,” exhibited relatively greater species-associated organization, consistent with the rapid evolutionary diversification of adaptive immune programs. In contrast, innate immune and progenitor populations, including “innate lymphoid cells,” “myeloid cells,” “dendritic cells,” “monocytes,” and “hematopoietic cell,” displayed comparatively stronger cell type-associated variation, suggesting greater conservation of lineage-defining transcriptional programs across species. Additionally, several differentiated cell populations (“epidermal cell,” “pneumocyte,” “melanocyte,” and “germline”) displayed comparatively greater species-associated signal, whereas nervous system cell types showed mixed signal dominance.

We next extracted cell type-specific species trees from the MILK hierarchy and compared each against the reference species phylogeny using a normalized triplets score (**Fig. 6g** and **Supplementary Fig. 6.3**). This analysis revealed substantial cell type-dependent variation in phylogenetic concordance. Among 34 cell types captured by at least three species, 13 exhibited perfect agreement with the reference species tree, 14 showed intermediate agreement, and 7 showed no detectable agreement (**Fig. 6h**). Projecting these cell type-specific phylogenetic concordance scores alongside their corresponding species-associated organizational signals within the global transcriptional hierarchy (**Fig. 6f**) may reveal how cellular programs are shaped by evolution and how cell states further diverge through system-intrinsic regulatory cascades. For example, germline cells, which directly mediate heredity and are central to speciation, showed both high phylogenetic concordance and strong species-associated organization signals. Similarly, among the cell type labels observed in both signal measurements, “stem cell” and “hematopoietic cell” showed moderate-to-high phylogenetic concordance together with species-associated organization signals, suggesting that these progenitor cell states retain transcriptional structures shaped by evolution. In contrast, their descendant adaptive immune populations, which showed high species-associated organization, except for NKT cells, exhibited low phylogenetic concordance despite high bootstrap scores, suggesting that their differentiation may be governed more strongly by species-intrinsic regulatory programs after the emergence of progenitor states. Together, these two analyses highlighted that some cellular programs appear to retain transcriptomic relationships that closely track evolutionary divergence, whereas others show greater departure from the species tree, consistent with lineage-specific remodeling, functional specialization, or convergent cell-state organization.

Collectively, these findings showcase how MILK trees can also provide a scalable and interpretable framework for contextualizing cell type and species information within a unified hierarchical structure. More specifically, integrating single-cell transcriptional landscapes across broader species diversity with MILK may facilitate deeper understanding of evolution–development coupling in the emergence of diverse multicellular systems.

## DISCUSSION

MILK enables the hierarchical representation of high-dimensional data objects at unprecedented scales. Although broadly applicable across data modalities, this framework is particularly well suited to single-cell genomics, where rapidly expanding atlases have outpaced our ability to integrate, compress, and extract interpretable biological information. This suitability reflects a basic property of multicellular life: cellular diversity emerges through iterative, lineage-dependent processes of cell-state expansion and diversification. A complex organism originates from a single zygote and develops through repeated cell divisions, progressive changes in molecular state, and continuous exchange of signals among neighboring cells and their environment. These processes are governed by the information encoded in a finite genome together with physical constraints, environmental inputs, regulatory circuits, and intercellular communication, with each cellular state shaping the conditions under which subsequent states emerge. Because a substantial component of this generative process is hierarchical, single-cell atlases should not be viewed simply as collections of independent molecular profiles, but as high-dimensional snapshots of structured biological processes unfolding across nested scales. From this perspective, tree representations provide a natural and efficient abstraction for compressing cellular state information while retaining a substantial fraction of the multi-resolution organization embedded in the biological system.

This view also changes how individual cell profiles can be interpreted. In many analytical settings, each cell is treated as an isolated high-dimensional object, defined primarily by its measured molecular features. Biologically, however, a cell is also the product of its progenitor history, intrinsic regulatory state, and interactions with neighboring cells. MILK places each cell within a global hierarchy in which its position is defined not only by its own molecular profile, but also by its relationships to other cells across multiple scales. The resulting tree provides a compact topological context for each cell such that nearby leaves represent closely related states, internal clades summarize shared structure, and higher-order branches reveal broader organization across the cellular landscape. Analogous to network biology, where representing genes and proteins within systems of interactions can provide functional context beyond individual molecular measurements^54,55^, hierarchical encodings of cells can make cellular identity more interpretable by assigning each cell a relative position within the organization of a multicellular system. Because a substantial component of cellular diversification arises through lineage-dependent and progressively constrained state transitions, tree topology provides a natural organizing layer for extracting structured biological signal from large collections of single-cell molecular profiles.

A second defining property of multicellular systems is that cellular state spaces are both heterogeneous and deeply redundant. Many states are represented by vast numbers of molecularly similar cells, whereas others are rare, transient or restricted to particular anatomical, developmental or disease contexts. This uneven density is not merely a nuisance of atlas-scale data; it reflects the organizational robustness of biology itself. Abundant populations represent repeated deployment of common physiological programs, whereas rare populations may correspond to specialized progenitors, signalling cells, immune subsets or transitional states that are essential for tissue function but easily lost during analysis. Yet, without a global representation of the cellular landscape, it is difficult to determine at what resolution redundancy should be compressed or rarity should be preserved. Random downsampling can make analysis tractable but risks distorting population structure, while exhaustive use of all cells rapidly becomes prohibitive for downstream analyses and model training. MILK addresses this problem by using tree topology as an empirical guide to cellular redundancy. Dense regions of the landscape can be compressed into representative clades, while sparse or distinct regions remain resolved until higher levels of the hierarchy. Representative subsampling therefore becomes resolution-adaptive rather than arbitrary, allowing cellular states to be selected according to their positions within the global topology.

This principle also suggests a practical route for foundation model training. Rather than relying predominantly on raw cell number, training could begin from representative cells sampled at coarser hierarchical resolutions and be progressively refined until performance saturates. Although construction of a MILK tree itself requires substantial computation at atlas scale, MILK remains accessible because it relies on CPU-based computation rather than accelerator-intensive model training. Moreover, a single hierarchical representation can support multiple resolutions, subsamples, and downstream analyses without recomputing the underlying structure. These properties could reduce the computational barriers to constructing and evaluating single-cell foundation models by prioritizing structurally informative diversity without requiring sheer scale.

Beyond representative compression, the hierarchical organization of MILK provides a practical substrate for extracting robust signal from noisy and heterogeneous single-cell data. Single-cell genomic profiles are discrete snapshots of dynamic biological entities at the moment of measurement, further shaped by technical factors such as dropout, batch effects, and platform-specific biases. Consequently, downstream interpretations can be highly sensitive to analytical choices, including clustering resolution, dimensionality reduction, batch correction, prior annotations and other forms of parameterization. MILK mitigates some of these dependencies by imposing a simple, data-driven hierarchy in which progressively coarser levels encode relationships among cells from local neighborhoods to global population structure. Once constructed, this hierarchy can be interrogated *post hoc* across multiple resolutions, allowing biological structure to be examined at different granularities without repeatedly reconstructing the underlying representation. this sense, MILK is not intended to replace existing single-cell workflows, but rather to provide a complementary organizational layer on which diverse analyses can be efficiently performed.

These principles were reflected across the analyses presented in this study. In the human fetal atlas, MILK converted uneven cellular redundancy into representative subsamples and clade-informed metacells, demonstrating that atlas-scale compression can retain biological variation rather than simply remove cells. In the mouse prenatal developmental atlas, MILK tree distances recovered temporal and cell type relationships directly from gene expression, indicating that transcriptomic hierarchies can preserve information about developmental organization without requiring explicit lineage measurements. In the CELLxGENE Discover Census, the same hierarchical structure provided a reusable substrate for evaluating census-trained embeddings, revealing how different models balance biological resolution against technical harmonization. In disease contexts, MILK contextualized perturbations within a global cellular landscape, allowing disease-associated shifts to be quantified through clade redistribution, locality and topological association. In the cross-species atlas, MILK further enabled cell type and species signals to be examined across hierarchical resolutions, providing a framework to distinguish conserved cellular programs from species-associated divergence. Collectively, these applications show that MILK can go beyond reducing dataset sizes, transforming large molecular atlases into queryable hierarchical structures in which cells, clades, metadata labels and biological contexts can be interrogated across scales.

The analyses presented here explored only a fraction of what may become possible once large single-cell resources are represented as reusable hierarchical objects. One immediate direction is to use MILK trees as multi-resolution pseudobulking frameworks. Because each clade defines a population of cells at a particular level of topological resolution, clade-level aggregation could generate statistically robust pseudoreplicates while preserving rare, transitional and context-dependent states, extending principles from metacell and pseudobulking analyses^47,56,57^. Such representations may improve differential expression analysis, regulatory program inference, multimodal integration and perturbation analysis by allowing biological signal to be summarized across scales rather than only within predefined clusters or manually curated labels.

A second opportunity is the synergistic coupling of MILK with cell lineage tracing and molecular recording technologies. Molecular recorders such as DNA Typewriter make it possible to recover ancestry and transcriptomic profiles from a single destructive single-cell readout^58,59^, and its recent demonstrations illustrate that dense scRNA-seq data can support cellular phylogeny analyses for tasks including clonal dominance, the coupling of developmental timing to cell fate, and ancestral-state imputation^60^. MILK could provide a complementary state-space framework for such analyses. Rather than treating cell type labels as fixed endpoints, phylogenetic branching events could be projected onto a MILK hierarchy constructed from genomic atlases at corresponding developmental time points. This would allow clonal dominance to be evaluated not only by clone size or annotated fate, but also by the extent to which clones occupy, diversify across or converge onto specific regions of molecular state space. Additionally, developmental fate decisions could be generalized from discrete labels to continuous MILK distances or clade transitions, identifying divisions that traverse major state boundaries. Finally, time-resolved tree reconstructions with time-course scRNA-seq datasets and ancestral-node imputation could be regularized by MILK-defined state neighborhoods, providing a principled approach to map lineage histories onto transcriptional trajectories and construct lineage-aware representations of consensus ontogenies. In such a framework, lineage trees describe where cells came from, whereas MILK describes where their molecular states reside. Their coupling could reveal statistical regularities linking ancestry, state diversification and developmental constraint across individuals, genotypes and perturbations.

Perturbational atlases provide another natural extension. Large Perturb-seq and drug-response resources, such as Tahoe-100M, could be projected onto the disease-contextualized MILK tree to identify therapeutic responses that move cells toward, away from or between disease-associated regions of the hierarchy^61–64^. Such analyses could reveal candidate core-response genes and regulatory programs that recur across distinct pathological contexts, while separating broadly shared stress, inflammatory, or proliferative responses from cell type- or disease-specific vulnerabilities. More generally, clade-level perturbation scores could provide interpretable, model-agnostic biomarkers of drug response, CRISPR perturbation, infection^65^, organoid maturation and other experimentally induced state changes, consistent with the broader vision of perturbational cell atlases for causal cell biology^66^.

A related application is the refinement of the metadata labels through which single-cell atlases are interpreted. Current cell type, disease, tissue, and species annotations are essential, but they are often assigned through study-specific clustering, marker selection, nomenclature conventions, and parameterization choices. Consequently, metadata-driven analyses, including disease dysregulation and cross-species comparison, inevitably inherit biases introduced by differences in annotation granularity, sampling density and labeling criteria across datasets. Rather than treating these labels as fixed ground truth, MILK trees could identify where expert annotations align with the global topology of cellular states and where they diverge. Coherent labels may define robust states, repeatedly co-localized labels may suggest candidates for harmonization, and labels that reproducibly split across branches may reveal overlooked subtypes, transitional programs or context-dependent states. This use of MILK would be analogous in principle to prior efforts in network biology, including NeXO^67^ and CliXO^68^, in which large-scale molecular and genetic networks were used to refine hierarchical representations of community-annotated GO terms of genes and gene function inference.

Finally, MILK may serve as an interpretability layer for foundation models. Foundation models learn high-dimensional latent spaces, but their predictions, uncertainties, and failure modes remain difficult to interpret biologically. Assigning model embeddings, predicted perturbation responses, confidence scores or prediction errors to leaves and clades of a MILK tree could reveal which regions of the cellular landscape are robustly represented and which are dominated by sparsity, batch structure or extrapolation. This direction is complementary to emerging single-cell foundation models and atlas-scale biological modeling efforts. Thus, MILK should be viewed not as a final representation of single-cell biology, but as a scalable topological substrate on which measurements, annotations, perturbations, lineage histories and predictive models can be organized, evaluated and interpreted.

At present, the scale of single-cell genomics often forces biological interpretation to proceed through slices of the full landscape: biological pathways, selected tissues, annotated clusters, disease subsets, sampled populations or individual model embeddings. These views remain essential, but they can obscure relationships that emerge only when cells are placed within a broader organizational context. MILK offers a general-purpose data representation by converting atlas-scale datasets into reusable hierarchical objects that can be compressed, queried, and extended across resolutions. In doing so, it shifts the unit of analysis from isolated cellular profiles to cells positioned within a global topology, where local neighborhoods, intermediate clades and broad branches each carry interpretable biological information. As single-cell resources increasingly incorporate perturbation, spatial organization, chromatin state, lineage history and foundation model predictions, such tree-based contextualization may enable analyses that ask not only which cells are present, but how they relate, how they shift across contexts and which relationships persist across individuals, diseases, species and experimental systems. In this sense, MILK points toward a transition in single-cell biology from the accumulation of molecular profiles to the structured understanding of cellular organization.

## ACKNOWLEDGEMENTS

We thank the members of the Yachie lab at the University of British Columbia and the University of Osaka, as well as the broader scientific community, for their constructive discussion and feedback throughout this study. We thank Arman Adel for suggesting the name “MILK” for this framework and Jay Shendure and Chengxiang Qiu for sharing the embryo images used in their original paper^19^. This study was funded by United Therapeutics Corporation, the Canada Research Chair program, the Canadian Institutes of Health Research (CIHR) Project Grants, the Canada Foundation for Innovation (CFI), the Allen Distinguished Investigator Award, the Japan Science and Technology Agency (JST)’s CREST Cell Control (yuCell) Program, the Japan Society for the Promotion of Science (JSPS) KAKENHI Grant-in-Aid for Transformative Research Areas (B), and the Takeda Science Foundation (all to N.Y.). This work was partially supported by World Premier International Research Center Initiative (WPI), MEXT, Japan. Part of the analysis was conducted using the SHIROKANE Supercomputer at the University of Tokyo Human Genome Center.

## AUTHOR CONTRIBUTION

B.K. conceived the original concept of MILK.

B.K. and N.Y. designed the study.

B.K. developed MILK.

B.K. and C.L. performed analyses.

B.K. and H.Y. worked on packaging MILK as a software tool.

B.K. and N.Y. interpreted the results.

B.K. and N.Y. wrote the manuscript.

N.Y. supervised the project.

## AI DISCLOSURE STATEMENT

We disclose that AI-based tools were used to support preliminary data exploration, data analysis, and manuscript writing. All interpretations and intellectual contributions were generated by the authors. The authors take full responsibility for the integrity and content of the work.

## DATA AND CODE AVAILABILITY

The open-source MILK software is available at https://github.com/yachielab/milk. The code used to perform the analyses presented in this study is also available at https://github.com/yachielab/MILK-paper.

A curated collection of analysis files generated or assembled throughout this study can be found at https://doi.org/10.5281/zenodo.22088317.

## METHODS

### Methods 1: MILK algorithm

MILK is a scalable algorithm implemented in Julia that obtains a tree representation over a population of interest by recursively grouping (representative) objects according to a data-driven similarity threshold.

#### Methods 1.1: Percentile-defined local grouping and representative assignment

The core process of MILK uses a distance-based method to cluster the population of input objects into groups of similar members according to a similarity threshold. This threshold is defined by a fixed percentile (user-specified) from the distribution of pairwise object distances. Given this threshold, the algorithm performs a single pass through the population. The first candidate object automatically forms a new group and serves as its representative. Each subsequent candidate is then compared to all existing group representatives based on pairwise distance. If the distance between the candidate and any existing groups is less than or equal to the threshold, it is considered “redundant”, in which case it is assigned to the group it is most similar to. Conversely, if the distance between the candidate and all existing groups exceeds the threshold, it is considered “distinct”, which results in the formation of a new group, where that candidate is designated as the group representative. The ability to use any pairwise comparison metric, coupled with a data-driven similarity threshold, allows MILK to generalize to any high-dimensional dataset with minimal parameterization.

#### Methods 1.2: Recursive representative aggregation, caching, and distributed execution

Using a data-driven threshold on the lower-extreme percentile quickly becomes computationally intractable because the number of comparisons grows quadratically with the number of sequences. To apply MILK to datasets comprising millions of high-dimensional objects, we devised a recursive framework that addresses two major obstacles.

The first modification partitions the input population, constraining the local grouping process (**Methods 1.1**) to computationally tractable subsets. Following data partitioning, a global similarity threshold is determined by independently computing fixed percentile values across many partitions and taking the minimum value observed. The grouping process can then be executed independently and in parallel according to the shared similarity threshold. The resulting groupings identified in each partition are then aggregated to form a global list of groups. If the total number of groups is tractable according to a user-specified cutoff, an additional grouping process is performed only on the group representatives. This additional aggregation step mitigates overlapping groups that were not merged because of the heuristic data partitioning step.

The second modification involves recursively applying this process with a similarity threshold based on the lower-extreme percentile of pairwise distances (e.g., the 0.01 percentile of observed pairwise distances). This aims to eliminate the arbitrary choice of an appropriate similarity threshold and instead emphasizes obtaining high-resolution pairwise relationships between objects in the population. In the grouping process, this results in most objects forming distinct groups, whereas only those exhibiting extremely high similarity will be clustered. Applying this method recursively to cluster similar objects results in a progressive merging of groups until a single group remains. This is also reflected in the recursive application of a fixed percentile, which is dynamically but monotonically updated at each iteration (from local to global).

To ensure that group representatives reflect the distribution of objects from previous iterations, representatives identified in each iteration are optimized to be the medoid, or the object closest to the centroid. Because the MILK process has access only to information for the input objects present at the current iteration, we further introduced a cache to provide additional resolution to the medoid optimization procedure. Once the number of representatives falls below a user-specified upper limit for the cache size, the information for those representatives is cached and included in all subsequent medoid optimization processes for their associated groups. Specifically, the cache information is included in the centroid calculation used to identify the medoid, which is particularly relevant when the group size is 2 objects, where selecting a medoid would otherwise be arbitrary without the additional cached information.

Notably, the MILK algorithm maintains the order of input data throughout. At scale, the partitioning step dictates the potential groups that can be formed. This behavior can be leveraged to encode prior knowledge by ordering cells based on metadata information, if desired. In this study, cells were ordered by dataset, reasoning that the likelihood of finding more similar cells is higher intra-dataset than inter-dataset or in a shuffled ordering, particularly as the diversity of compiled transcriptomic datasets grows. Alternatively, users can shuffle the order of cells prior to running MILK.

The independent processing of data partitions is conducive to out-of-core computing, bypassing the need to load the complete dataset into memory during task execution. This becomes increasingly necessary as dataset sizes continue to grow, and it also presents an opportunity to leverage both distributed computing processes and cluster computing nodes in high-performance computing environments. As a result, the MILK implementation is designed to handle the growing scales of high-dimensional information.

Altogether, MILK can capture high-resolution hierarchical relationships in a data-driven manner, from the bottom (individual objects) to the top (root of the tree). Despite similarities in the hierarchical tree output, conventional agglomerative (hierarchical) clustering algorithms require explicit computation of an all-all distance matrix, from which clusters are iteratively merged according to a linkage criterion. Not only are these conventional approaches unable to scale to millions of cells, but they can also be highly sensitive to linkage criterion, variable density of objects, and the precise contextualization of rare subpopulations.

The strategy of circumventing exhaustive pairwise comparisons by restricting object comparisons to group representatives is conceptually related to scalable sequence clustering approaches^69–71^. However, the clustering outcomes with these methods were found to be highly sensitive to user-defined similarity parameters, leading to variability in performance across datasets and analysis tasks (data not shown). In contrast, MILK eliminates this parameter dependency during execution by maximizing resolution in each iteration and applying the process recursively, enabling post-hoc, data-driven selection of similarity relationships embedded in the resulting hierarchical levels of the tree structure.

#### Methods 1.3: MILK demonstration on handwritten digit embeddings

The Modified National Institute of Standards and Technology (MNIST) dataset comprises pixelated handwritten digits (0 to 9) and is widely used as a benchmark to evaluate the classification accuracy of machine learning models^46^. We downloaded a reduced version of this dataset containing 1,797 observations from the scikit-learn library^72^ using the `sklearn.datasets.load_digits` call, where each image was represented as an 8×8 grid of pixels (i.e., 64 dimensions).

We applied MILK to the MNIST dataset using cosine distance and a percentile threshold of 0.0, which corresponds to the minimum observed pairwise distance at each iteration. Representatives were depicted within the UMAP embeddings of the complete dataset across iterations of the MILK procedure. The overall MILK tree was visualized using the “dendrogram” layout provided by the ggraph package^73^ in R.

#### Methods 1.4: MILK subsampling of simulated phylogenetic nucleotide sequences

To evaluate whether MILK can generate representative subsamples from nucleotide sequences, we applied it to simulated phylogenetic datasets generated using the PRESUME software (https://github.com/yachielab/PRESUME). PRESUME first simulates a rooted tree topology under specified phylogenetic parameters. Next, given a randomly generated root sequence, mutations were introduced and propagated along the tree branches according to a General Time Reversible nucleotide substitution model with rate heterogeneity modeled according to the Gamma probability distribution (GTR-GAMMA). This process yields mutated nucleotide sequences at the leaves of the tree along with the corresponding ground truth tree structure.

Phylogenetic datasets were simulated with a target size of 100,000 sequences, each 1,000 base pairs in length. Neither insertions nor deletions were included in the simulation of sequence evolution. To generate phylogenies exhibiting varying degrees of topological imbalance, we varied the `s` parameter (standard deviation of generation time between sister branches) from 0 (fully balanced trees) to 1.4 (highly imbalanced trees) in increments of 0.2 (n = 3 simulation replicates). The final number of sequences per dataset varied from the target, particularly at higher values of `s` where increased topological bias strongly influenced the simulated branching dynamics. Default values were used for the remaining parameters. The Colless index, which sums the absolute difference in the number of descendant leaves between child subclades across internal nodes, was used to confirm that phylogenies with varying degrees of topological imbalance were generated.

MILK was applied with no recursion and a sequence similarity threshold set to the 5^th^ percentile of pairwise Hamming distances. This resulted in approximately 100 groups, from which stratified sampling generated subsamples (n = 10). These subsamples were then evaluated for their ability to capture diversity across the complete datasets. Random subsamples from the complete dataset (sample size = 100; n = 100) were included as a baseline.

The first analysis utilized the ground truth tree topology. In particular, the ground truth subtree corresponding to MILK and random subsamples were extracted, and branch lengths were optimized with RAxML-NG^74^ software according to the GTR-GAMMA model. The total summation of branch lengths was then used as a measure of subsample representativeness, with the rationale that higher values correspond to greater diversity of subpopulation structure captured.

The second analysis involved computation of the complete pairwise Hamming distance matrix across all 100,000 sequences. Agglomerative clustering (Scanpy implementation) was then applied to the distance matrix with the number of clusters set to be equal to the subsample size. Subsample quality could then be assessed based on the number of distinct clusters the MILK and random subsamples captured.

### Methods 2: Human fetal atlas subsampling and metacell benchmarking

#### Methods 2.1: Human fetal organ atlas data acquisition

The raw gene expression read count matrices of 15 fetal organs (adrenal, cerebellum, cerebrum, eye, heart, intestine, kidney, liver, lung, muscle, pancreas, placenta, spleen, stomach, and thymus) were downloaded from the Descartes platform provided by the Brotman Baty Institute (https://descartes.brotmanbaty.org/bbi/human-gene-expression-during-development/).

#### Methods 2.2: Human fetal organ atlas preprocessing

Each organ dataset was processed independently using a standard single-cell analysis pipeline implemented in Scanpy^75^. As in the original publication^40^, datasets were initially filtered to include only “protein_coding”, “lincRNA”, and “pseudogenes” for each cell. Cells with fewer than 100 expressed genes and genes detected in fewer than 10 cells were excluded. Putative doublets were then identified and removed using Scrublet^76^ with the following parameters (default if unspecified): `batch_key` = “Fetus_ID”, `sim_doublet_ratio` = 2, `expected_doublet_rate` = 0.06, and `n_neighbors` = 10. Read counts for each cell were library-size normalized using `scanpy.pp.normalize_total` with default parameters, such that each cell had the same total count after normalization, and were subsequently log-transformed using `scanpy.pp.log1p`. Highly variable genes (HVGs) were identified per cell type group using default parameters, with the batch key specified as “Assay” (referring to cell versus nuclei transcriptomic profiling). Lastly, only cells that were annotated with a cell type in the original publication were considered, as these were used to define biologically relevant groups in downstream analyses.

Each organ dataset was then subjected to principal components analysis (PCA) with 100 principal components, from which the k-nearest neighbors (KNN) graph was computed using the cosine distance metric and 50 nearest neighbors to preserve global structure. The two-dimensional UMAP embeddings were determined with a `min_dist` = 1 and `spread` = 3.0.

#### Methods 2.3: Organ-specific MILK tree construction

MILK was applied on the PCA embeddings of each single-cell organ dataset using cosine distance as the pairwise comparison metric and a threshold fixed to the 0.1 percentile. Cosine distance was utilized for consistency with the analyses in the original publication. The partition size and upper limit cache size were set to 10,000 cells. A merging threshold for aggregation across partitions through an additional stratification process was set to 50,000 cells.

#### Methods 2.4: Organ-specific MILK tree visualization and summary metrics

Initially, the recursive MILK procedure was qualitatively assessed across the 15 organ datasets in terms of the distribution of representatives in the context of the UMAP embeddings from the complete dataset. The complete MILK tree hierarchies were further visualized per organ dataset using the `dendrogram` layout in the ggraph package in R or the network `sfdp_layout` provided in the graph_tool^77^ Python library, with nodes colored by cell type and edges tracing the grouping events across recursive iterations. Additionally, the cell type compositions among group representatives were visualized with an alluvial plot across recursive iterations.

We next quantified the number of groups and distribution of group sizes across recursive iterations, and further compared group members to their representative in terms of cell type agreement, approximate cosine distance, and normalized specificity. Briefly, cell type agreement was calculated per group as the proportion of group members with the same cell type as their respective representative. The approximate distances were defined as the average cosine distance across group members (leaves of the tree) to their respective representatives (internal nodes), pooled across all groups. Lastly, specificity was computed as the number of groups a cell could have mapped to according to the similarity threshold (cosine distance to other group representatives less than or equal to the global threshold), normalized by the total number of groups at that iteration. Overall, these analyses provided general insights into the distance-based process across diverse organ contexts.

#### Methods 2.5: Representative subsampling from fetal organ MILK trees

A primary use case of MILK trees is the generation of representative subsamples from large-scale single-cell datasets. Representative subsampling can be defined as the process of identifying a subset of cells that effectively captures the transcriptomic variation observed in the overall population. In the context of MILK, this can be formulated as identifying a certain number of non-overlapping clades according to a specified target sample size such that coverage over the leaves of the tree is maximized. For this study, we also aimed to address class imbalance with this process by generating subsamples with roughly equal representation from each group (cell type annotation or data-driven MILK tree cluster), as such biases have been shown to influence downstream analyses towards dominant subpopulations^78^. In the current study, we investigated two strategies to extract representative subsamples from the reconstructed MILK tree, which we refer to as label-free and label-balanced.

##### Methods 2.5.1: Topology-guided label-free subsampling

Label-free subsampling extracts representative subsets based solely on the structural topology of the MILK tree. In particular, according to a minimum clade size cutoff parameter (starting at an upper bound of 25), a list of all viable clades in the MILK tree was obtained and ordered by recursive MILK iterations, from early-to-late (leaves-to-root), as this corresponds to the resolution of similarity at which group formation occurs. The subsample was then constructed by sequentially sampling non-overlapping clades from this list until the target sample size is reached.

If the non-overlapping constraint prevents the target size from being reached, the minimum clade size cutoff parameter is iteratively decremented (to a lower bound of 10), and the selection procedure is repeated. If the subsample size remains lower than the target sample size after this stage, a clade expansion phase occurs, whereby subsampled clades corresponding to the earliest iterations are progressively replaced with their child clades.

This approach capitalizes on the observation that rare subpopulations tend to cluster early in the MILK hierarchy due to their strong, intra-group transcriptomic similarity and/or relatively larger inter-group distances to other subpopulations. As such, prioritizing representation from these robust subpopulations was found to be an effective approach to capture the overall structure of the population and balance cell abundance per group.

##### Methods 2.5.2: Cell type-balanced topology-constrained subsampling

Label-balanced subsampling leverages metadata labels to guide the subsampling process from the MILK tree, which can be useful when prior group markers exist. Cell type annotations were used in this analysis. This approach starts by determining a cutoff number of cells to sample per cell type by dividing the target sample size by the number of unique cell types. If the number of cells for a given cell type is below this cutoff, all cells belonging to that group will be trivially included in the subsample.

Next, a list is initialized for each cell type composed of all constituent individual cells at the leaves of the MILK tree. At each level in the MILK tree, the algorithm examines each cell type list, looking to merge sets of cells if they directly correspond to a branching event in the MILK hierarchy. This merging can only occur if all cells involved in the clade grouping match the cell type of interest (i.e., no cell type label mixing) and results in the replacement of the corresponding cells by their higher-level representative, thereby reducing the size of the cell type list. If at any level in the traversal the number of (representative) cells in the cell type list falls below the cutoff, then all clades in the list are included in the subsample and that cell type subsample is set. Conversely, if the complete hierarchy is traversed without reducing the cell type list below the target size, the remaining clades are ordered by group size (large-to-small), and the target number of representatives is selected.

This approach leverages the pre-existing metadata information that may be biologically meaningful without forcing groupings that are unsupported in the MILK tree. The requirement of cell type groups exhibiting agreement with the MILK tree topology acts as an additional filter that selects for coverage within subgroup topologies. Based on this approach, the information coverage over the complete dataset will increase as the concordance between the MILK tree and cell type groups increases, whereas it will appear equal to random subsampling per cell type if there is no concordance between the MILK tree and metadata labels.

##### Methods 2.5.3: Fetal organ subsampling design and random baselines

MILK-derived subsamples were generated according to label-free and label-balanced strategies for each organ dataset at varying target sample sizes (100, 300, 1000, 3000, 10000, and 30000 cells). Organ-specific subsamples were compared to random baselines (n = 10), which were also derived for both strategies by sampling without replacement across the complete dataset or within cell type groups, respectively. Notably, the label-free and label-balanced MILK tree subsampling approaches utilized in this study were deterministic, so only a single replicate was generated per target sample size.

Random subsampling using label information represents a practical approach to downsampling large-scale data, where subsample quality is contingent on the precursor work of identifying robust groups (e.g., cell types). However, effective annotation at scale remains a non-trivial task. One of the main disadvantages of random downsampling of large-scale datasets is the loss of information that is incurred, as the vast majority of cells must be discarded to achieve tractable dataset sizes for downstream analyses. This loss of information affects both rare subpopulations as well as heterogeneous dominant subpopulations. In contrast, MILK trees provide a framework for representative subsampling where the preserved links of sampled cells to downstream clades comprised of non-sampled cells can implicitly retain information.

#### Methods 2.6: MILK-derived metacell construction by clade-level pseudobulking

For representative subsamples derived from MILK trees (as detailed in **Methods 2.5**), each subsampled cell may correspond either to a single observed cell or to an internal node, or clade, in the MILK tree hierarchy. This representation can therefore provide a principled approach to retain information from non-subsampled cells, as they can be structurally linked to their corresponding representative.

One potential strategy to explicitly leverage the information provided by the additional, non-subsampled cells involves the aggregation, or pseudo-bulking, of transcriptional profiles to construct “metacells”. Such an aggregation of highly similar cells has been found to mitigate challenges surrounding technical dropout and biological stochasticity, potentially leading to more robust downstream analyses^47^.

In the human fetal cell atlas, metacells were generated by aggregating raw read counts for the identified groups (i.e., clades) in the representative organ subsamples. The pseudobulked profiles were obtained by calculating the weighted average of raw gene counts across all cells in the clade, where global weights were determined by initially compiling the list of cosine distance values of every member to its respective representative in PCA space, and then applying min-max scaling per group member as follows:

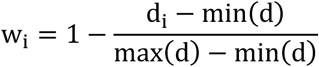

In the equation above, d corresponds to the complete vector of distance values, and d_i_ corresponds to the distance value for a given member to its representative. Following scaling, the weights were defined as the complement of scaled distance values, or w_i_ for a given cell i. For each subsampled cell, the pseudobulking step can be represented by the following formula:

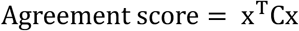

The raw counts for each gene in the transcriptome, gene ∈ G, are averaged across all cells in the MILK clade, cell ∈ Clade. The global weights ensure that cells closer to the medoid (representative) contribute greater influence on the resulting metacell gene expression profiles.

#### Methods 2.7: Cell type-specific regulon preservation after downsampling

To assess whether subsampling can effectively retain the biological signal observed in the complete dataset, we tested the preservation of mean and variance statistics of measured gene expression. However, given the intrinsic noise associated with each individual gene in the transcriptomic profile, we assessed the preservation of gene expression signal in aggregate according to sets of genes. In particular, we downloaded the CollecTRI (Collection of Transcriptional Regulatory Interactions) meta-resource consisting of 1186 transcription factors (TFs) and over 43,175 TF-gene interactions derived from public databases, text mining, and manual curation^48^. For each TF, the set of its interacting target genes formed a regulon, which we utilized as biologically meaningful sets of genes for evaluation.

To mitigate the effects of population bias from dominant cell type groups and constrain comparison to biologically similar groups, measurements between the different subsamples and complete dataset were stratified by cell type. This ensures rare cell types contribute the same amount as dominant subpopulations, while preventing dilution effects due to contradictory gene expression signal.

##### Methods 2.7.1: Regulon enrichment scoring

For each organ dataset, individual cells were scored against the list of regulons to obtain regulon enrichment profiles. The regulon enrichment score represents the average expression of the set of regulon genes subtracted by the average expression of a randomly sampled set of background genes, where background genes are binned in terms of expression value^79^. This enrichment score analysis was calculated using the `score_genes` function in the Scanpy library. In this analysis, each regulon was further separated based on the inferred upregulation and downregulation relationship to its respective TF, as provided in the CollecTRI database. Computing these scores across all regulon sets resulted in a regulon profile for each individual cell.

The Spearman correlation of regulon scores per cell type was computed between the subsampled and complete datasets to quantify the extent to which subsamples retain biological signal. Using these correlation values, we compared MILK-derived subsamples to random subsampling baselines with respect to both label-free and label-balanced approaches. For MILK-derived subsamples, an additional metacell group was included. The rank-based Spearman metric was used to appropriately assess the preservation of regulon signal despite such systematic differences in expression values between the single-cell profiles and the aggregate metacell profiles.

Each cell type-specific correlation value was plotted as a function of the proportion downsampled from the complete dataset, which resulted in a continuous distribution of values of pooled organ datasets. Fitted lines were provided for each subsampling approach, which were further summarized by the area under the curve (AUC) with respect to the log-transformed proportion that each cell type was downsampled. Lines were fit (`geom_smooth` function in ggplot2^80^ R package) according to the non-parametric LOESS (locally estimated scatterplot smoothing) regression method. AUC was computed using the MESS^81^ package in R.

##### Methods 2.7.2: Regulon covariance preservation using principal-component subspace angles

We next assessed whether regulon-specific covariance was preserved following downsampling using an eigenvector-based analysis. Following processing (see **Methods 2.2**), for each cell type and regulon combination, gene expression profiles for the corresponding subset of cells and genes were subjected to PCA with the number of components set to the minimum between the number of cells, genes, or a value of 10 to focus on dominant principal components. Only regulons with at least 2 target genes were considered in this analysis. Due to the large number of cells in certain groups, incremental PCA (scikit-learn implementation) was utilized across all groups for consistency.

Preservation with respect to the directionality of dominant variation in regulon expression between subsampled and complete datasets was calculated by initially measuring the alignment of the two principal component subspaces using the `subspace_angles` function in the SciPy library^82^ in Python. The angle preservation could subsequently be quantified in terms of cosine similarity, and averaged across subspace dimensionality. A cell type-specific average was obtained by aggregating the cosine similarity values across all regulons per cell type group. Plotted as a function of the proportion that each cell type was downsampled relative to the complete dataset, a fitted line using the LOESS metric was provided for the various subsampling approaches, which were summarized by their AUC value.

#### Methods 2.8: scVI integration of representative fetal organ atlases

Representative subsamples from each organ dataset were integrated with scVI (single-cell variational inference) to mitigate batch effects and enable global cell comparisons. The input to the scVI model was the raw count data of the concatenated organ subsamples per target sample size. Prior to integration, data dimensionality was reduced to only include the top 5000 HVGs across all concatenated organ datasets. The model was configured with “Fetus_id” as the batch key, and training involved the following parameterization: `n_latent` = 100, `n_layers` = 2, `n_hidden` = 256, and `max_epochs` = 500. These parameters were selected due to the complexity and diversity in cell states spanning diverse organs and cell types. The integration step provided latent embeddings (100 dimensions) for representative atlases derived from MILK and random subsamples, both with and without the use of cell type information during subsampling.

For MILK-derived subsamples, integrated atlases were also constructed from the metacells that were derived from pseudobulking. However, this required an additional step of rounding the aggregated gene expression values to the nearest integer due to the scVI requirement of input read counts to be integers.

#### Methods 2.9: Leiden clustering assessment of integrated fetal atlases

Representative atlases derived from the downsampled organ datasets were evaluated in terms of the adjusted rand index (ARI) of Leiden clusters and cell type labels. Clusters were determined based on the KNN graph (k = 50), and the Leiden clustering implementation provided by Scanpy was used. ARI scores across subsampling approaches were computed across a range of Leiden clustering resolutions (2^0^ to 2^10^, exponent steps of 1), with the resolution yielding the maximum ARI score used to compare subsampling approaches across target sample sizes per organ (100, 300, 1000, 3000, 10000, and 30000 cells). Briefly, ARI measures the similarity between two lists of clusters after being adjusted for chance. A score of 1 indicates exact agreement between clusters, whereas a score of 0 denotes effectively random labeling.

Label-balanced and label-free subsampling approaches were assessed relative to random baselines. For MILK-derived subsamples, representative atlases composed of metacells were included.

#### Methods 2.10: scArches query mapping to representative fetal reference atlases

To assess how “representative” the downsampled atlas is over the overall population, we performed an analysis that involved mapping the remaining, non-subsampled cells onto the integrated global human fetal cell atlases, which were composed of the downsampled organ datasets at varying target sample sizes. Using the transfer learning approach from scArches^35^, query cells were mapped onto the scVI-integrated representative atlases. Notably, fine-tuning of the reference integration was disabled to ensure that performance in query mapping is a direct reflection of the integrated reference.

In treating a non-subsampled cell as a query and the scVI-integrated atlas as the reference, the average neighbor cosine distance (k = 8) of the query cell to reference cells of matching cell type was computed following mapping with scArches. Averaged by cell type, a lower neighbor distance suggests a more representative reference atlas, whereas a higher average neighbor distance indicates poor representation of the complete dataset in terms of cell type structure. Data corresponding to each subsampling approach were fitted with a LOESS curve and plotted as a function of the proportion that the global atlas was downsampled. AUC values summarized the subsampling approaches with respect to the log-transformed proportion that the complete atlas was downsampled from.

### Methods 3: Mouse prenatal atlas analysis of developmental hierarchy

#### Methods 3.1: Mouse prenatal atlas data acquisition

The raw count information for the mouse prenatal developmental atlas was obtained from the Chan-Zuckerberg CELLxGENE census (https://cellxgene.cziscience.com/collections/45d5d2c3-bc28-4814-aed6-0bb6f0e11c82). Captured with sci-rna-seq3 technology, Qiu et al. (2024) profiled the transcriptomic profile of over 12 million nuclei (11,441,407 after processing) from 83 embryos spanning late gastrulation to birth. In this paper, we utilized the Theiler stage annotations (12-27), which grouped each embryo into coarse development stages based on morphological and anatomical features.

#### Methods 3.2: Mouse prenatal atlas preprocessing by Theiler stage

Theiler stage datasets were processed independently using a standard single-cell analysis pipeline in Scanpy. Datasets were initially filtered to only include cells with at least 100 expressed genes and only genes with at least 3 cells. Features were filtered to include “protein_coding”, “lincRNA”, and “pseudogene”. A total of 11,441,407 cells remained following processing. Cells were then normalized and log-transformed using default parameters, and HVGs were annotated with default parameters. Each developmental stage dataset was then subjected to principal component analysis (PCA) with 100 principal components, followed by computing the KNN graph (k = 50) using the cosine distance metric, and UMAP with a `min_dist` of 1 and a `spread` of 3.

#### Methods 3.3: Theiler-stage MILK subsampling

MILK was applied on the principal components for each Theiler stage dataset with cosine distance as the comparison metric and a threshold fixed to the 0.1 percentile. The upper limit cache size and merging threshold (for aggregation across partitions through an additional stratification process) were set to 50000 cells. The remaining parameters were kept as default.

The number of cells in the Theiler development stage datasets varied between 46,253 and 2,092,526 cells after processing. Representative subsamples were obtained via label-balanced subsampling (see **Methods 2.5.2**) using cell type information (“author_cell_type” variable provided in the original publication^19^) from each Theiler stage MILK tree according to three target sample sizes (10000, 30000, and 100000 cells per Theiler stage dataset).

#### Methods 3.4: Global mouse developmental MILK tree reconstruction

Preliminary analyses qualitatively indicated a minor degree of batch effect across embryos in the Theiler stage datasets (**Supplementary Fig. 3.1**). However, the apparent embryo-specific batch effects were confounded by developmental time. Therefore, to preserve as much biological signal as possible, we did not perform batch correction, and the downsampled Theiler stage datasets were concatenated to form the global atlas. This decision was also based on literature evidence that the experimental results utilizing sci-rna-seq3 technology were accompanied by minimal batch effects^2,19^.

Following concatenation of MILK-derived subsamples across developmental stages to form a balanced prenatal mouse developmental atlas in terms of cell type abundance and Theiler stage representation across development, we applied MILK a second time (same parameterization as **Methods 3.3**) to capture the hierarchical organization of the population.

The MILK tree reconstructed over the representative atlas of mouse prenatal development was visualized using a scalable, force-directed network layout (`sfdp_layout` in the graph_tool Python library). Inferred using a repulsive force parameter set to 100, nodes were colored by Theiler stage or cell type annotations, with node size corresponding to group size and edges denoting grouping dynamics across MILK recursions.

#### Methods 3.5: Comparison with trajectory inference methods

To test whether MILK trees were consistent with established trajectory inference methods, we compared their respective trajectory measures to changes in development stages for sampled pairs of individual cells. Specifically, for the global atlas derived from downsampled Theiler stage datasets (target sample size of 30,000 cells), the developmental progression between a pair of cells was defined as the absolute difference in Theiler stages. The underlying hypothesis was that sampled pairs of cells sharing more similar transcriptomic profiles would, broadly speaking, be found closer in Theiler stages. Trajectory inference measures derived from the MILK tree, Diffusion Pseudotime (DPT), and WaddingtonOT (WOT) were assessed on the representative mouse atlas in terms of their ability to capture general trends in the context of a complex, large-scale developmental atlas of transcriptomic information.

##### Methods 3.5.1: MILK tree distance between cell pairs

For a pair of randomly sampled cells, the tree distance with respect to their corresponding leaves in the MILK tree was computed, where the recursive iterations in the MILK process were converted into tree unit branch length (=1) depths. For degenerate branches, or nodes with a singular child, the depth of the path was assigned as its largest depth at which it is degenerate, which corresponds to the earliest iteration at which the grouping event occurred. The tree distance of cell pairs was scalably computed by identifying the most recent common ancestor (MRCA) for the leaves corresponding to sampled cell pairs and then adding the absolute change in depths from the MRCA to each leaf. MILK tree distances were computed for 100,000 pairs of randomly sampled cells with no constraints on the cell pairs that were considered.

##### Methods 3.5.2: Diffusion Pseudotime within major cell groups

Pseudotime values were computed between pairs of cells within the same major cell groups. Briefly, a source cell was randomly selected in each major cell group, after which a diffusion-based pseudotime method was applied to infer the progression of remaining cells in that major group with respect to the source cell. Specifically, a diffusion map with 100 components was initially constructed, such that pseudotime values correspond to geodesic distances along the graph^50^. The pseudotime values of cell pairs consisted of the source cell to all other cells in the same major group and were compared against the developmental progression.

The cell populations included in this atlas span complex trajectories across developmental time, resulting in disconnected subgraphs that introduce ambiguity when comparing non-local pseudotime values. As such, cell pair comparisons were constrained to be within the same major cell groups, given that they each formed continuous trajectories spanning multiple Theiler stages. In addition to investigating the relationship between Theiler stage changes and pseudotime values, its relationship to MILK tree was also directly evaluated. Specifically, the Pearson correlation coefficient was computed between the pseudotime values and three different measures derived from the MILK tree for cell pairs: (i) recursive iteration of the MRCA; (ii) clade size of the MRCA; and, (iii) the percentile-based threshold of the MRCA.

##### Methods 3.5.3: WaddingtonOT transport mass across Theiler stages

Lastly, we performed an optimal transport analysis over developmental stages with WOT using the following setup: Theiler development stage was provided for the `cell_days` argument, `cell_growth_rates` were inferred from a time-course of induced pluripotent stem cells provided in the “wot tutorial” (https://broadinstitute.github.io/wot/tutorial/), and `growth_iters` was set to 3. Given this configuration, temporal couplings were inferred between cell pairs of adjacent time points with optimal transport. The resulting transport maps represent a probabilistic relationship describing the “descendent”, or transport mass, of a given cell at t_i_ to each cell in t_i+1_.

After inferring the transport maps for each developmental stage transition, all cell pairings across different Theiler stages were exhaustively assessed, in which the long-range temporal coupling was calculated by multiplication of transport value matrices. While this inference task could only be applied to cells from different developmental stages, its explicit use of development stage information makes it a powerful baseline.

##### Methods 3.5.4: Rank-binned comparison of developmental trajectory metrics

To compare the general trends of the three trajectory measures to the developmental progression for pairs of cells, MILK tree distance, pseudotime, and WOT values were independently ranked and then binned (n = 1000 bins), from which the average difference in Theiler stages was computed per bin. For consistent comparison, the transport mass values derived from WOT were ranked in the reverse order, such that similar cells correspond to smaller inverted transport values (positive relationship). The binned average developmental progression values for each trajectory inference method were plotted as a function of the bin ranking, and each method was fitted with a LOESS curve.

#### Methods 3.6: PAGA connectivity comparison with MILK clades

Partition-based graph abstraction (PAGA) is a method that aims to reconcile the continuous patterns in single-cell gene expression with the discretized structures of clusters of cells^52^. By specifying cell type groups to be treated as partitions, it estimates the connectivity across these partitions as interpretable abstractions, such that cell types sharing many transcriptional patterns will have higher connectivity scores than pairs of cell types with dissimilar expression profiles. PAGA was run with default parameters using cell type labels as groups, which outputs an adjacency matrix with values between 0 and 1 (1 indicating a “strong” connection) denoting the connectivity strength between cell types.

To evaluate MILK trees against the PAGA connectivity matrices, each internal node was decomposed into a cell type composition vector, where the count of each cell type among leaves was divided by the clade size. These clade-specific cell type frequencies were then compared to the PAGA matrix using a basic quadratic form equation as follows:

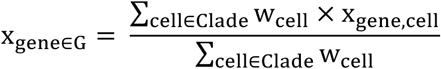

Where x refers to the vector of cell type proportions for a given clade in the MILK tree and C is the connectivity matrix derived from PAGA. Agreement scores were interpreted as the concordance between the relative cell type proportions captured by each clade in the MILK tree and the global connectivity of cell types. By evaluating agreement across all clades in the tree, this analysis provides a comprehensive summary of how well the MILK hierarchy captures the diverse cell type relationships.

The complete MILK trees corresponding to all Theiler stages were evaluated, as well as the global representative atlases of prenatal mouse development that were constructed according to target sample sizes of 10000, 30000, and 100000 cells per Theiler stage. The results were compared alongside a shuffle control, in which cell type labels were shuffled among leaf cells in the MILK trees.

#### Methods 3.7: Curated mouse developmental cell type graph preprocessing

A graph of cell type relationships spanning the prenatal mouse developmental atlas was proposed in the original publication^19^. While most cell types in the expression data could be uniquely mapped to a cell type in the proposed graph, the following cell types were manually mapped to their closest analog based on developmental time point:

**Table 1.** Cell type label mappings for consistency in downstream analyses.

| Cell type labels in the dataset | Cell type labels in the proposed cell type graph |
| --- | --- |
| Mast cells (P2rx7+) | Mast cells |
| Border-associated macrophages (Cd74+) | Border-associated macrophages |
| Lateral plate and intermediate mesoderm | Lateral plate mesoderm |
| Spinal cord dorsal progenitors | Spinal cord dorsal progenitors (after E13.0) |
| Spinal cord ventral progenitors | Spinal cord ventral progenitors (after E13.0) |
| Glutamatergic neurons | Di/mesencephalon glutamatergic neurons or<br>Glutamatergic neurons (after E13.0) |
| GABAergic neurons | Di/mesencephalon GABAergic neurons or<br>'GABAergic neurons (after E13.0) |

Notably, the proposed cell type graph was not strictly a directed acyclic graph (DAG), due to the presence of multiple root nodes, which resulted in some internal nodes having multiple parents. To simplify downstream analyses, a DAG was extracted from each root node, only retaining nodes with paths connecting to the respective root. Each DAG could then be independently traversed.

#### Methods 3.8: Asymmetric weighted Jaccard comparison with the developmental cell type graph

The global atlas MILK tree derived from downsampled Theiler stage datasets (target size of 30,000 cells) was compared to the “ground truth” tree of cell type relationships proposed in the original publication (see **Methods 3.7**). To compare the single-cell MILK tree to the ground truth, an asymmetric weighted Jaccard index was applied. Specifically, each clade in the MILK tree was decomposed into a cell type vector, where clade-specific cell types were divided by their total counts in the complete dataset. This vector of relative cell type proportions was then utilized as weights for the weighted Jaccard calculations against all clades in the ground truth cell type tree. The maximum weighted Jaccard index of each MILK clade against all ground truth clades was taken as that clade’s agreement with the ground truth tree. All cell type weights in the ground truth tree were set to 1 to reflect the relative proportions of cell types captured by the single-cell MILK tree to the aggregate-level cell type relationships.

In this weighted Jaccard analysis, the numerator corresponds to the intersection of cell types captured by both clades (MILK tree and ground truth), with values set as the minimum cell type weights between the two clades, or effectively the relative proportions of the MILK clade. The denominator represents the union of cell types captured in both clades, with the maximum cell type weights between the MILK clade and ground truth clade. The weighted Jaccard indexes for each clade were then normalized by the corresponding average weighted Jaccard indices across 10 shuffle control replicates to obtain a fold change. These fold-change values were plotted as a function of clade size to assess agreement between the MILK tree and ground truth.

#### Methods 3.9: Cell type-specific branching depth in the developmental MILK tree

To test whether cell type subpopulations exhibit localization bias in the MILK tree topology, the weighted average tree depth of each cell type was calculated.

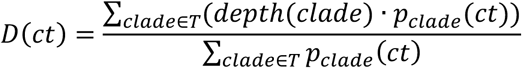

Parsing all internal nodes in the MILK tree, *clade* ∈ *T*, the clade-specific counts for each cell type were divided by the total cell type counts to obtain the relative proportion of each cell type captured in each clade, which is denoted as *p_clade_*(*ct*), where *ct* corresponds to the cell type of interest. For cell types with a relative proportion of at least 0.1, the tree depth (unit branch length) of viable clades and the corresponding relative proportion values were stored in a list. The average tree depth weighted by the relative proportion in a given clade was then computed for each cell type.

The weighted average tree depth of each cell type was compared against the average development stage of each cell type in terms of a weighted Pearson correlation (weights corresponding to the total number of cells of that cell type in each Theiler stage). Similarly, weighted average tree depth values were aggregated and plotted per major cell clusters to assess higher-level trends of biological information embedded in the MILK tree topology.

### Methods 4: CELLxGENE Census embedding benchmarks

#### Methods 4.1: CELLxGENE Census data, metadata and model embeddings

The complete CZ CELLxGENE Discover census (version: 2024-07-01, organism: “homo_sapiens”) of scRNA-seq data was downloaded in batches (batch size of 100,000 cells) using the SOMA (“Stack Of Matrices, Annotated”) schema (https://chanzuckerberg.github.io/cellxgene-census/). Downloaded as `anndata` objects, these datasets provided the raw gene expression counts and standardized metadata information for approximately 71 million cells, which were independently annotated by the submitting authors of each dataset.

Additionally, the latent representations for scVI (100 dimensions), Geneformer (512 dimensions), and scGPT (512 dimensions) of the CELLxGENE census were downloaded. Briefly, scVI is a variational autoencoder that models gene expression to approximate the zero-inflated negative binomial distributions that underlie the observed values. Originally developed for dataset-specific integration, this probabilistic approach has displayed strong performance across data integration benchmarking analyses^31,34^. However, the abundance of single-cell RNA seq data has recently led to the emergence of data-driven, transformer-based model architectures such as Geneformer and scGPT. Geneformer is a context-aware, attention-based model that features a normalized rank-scoring representation of gene expression to prioritize genes that distinguish unique cellular states^36^. In contrast, the scGPT model employs stacked transformer layers with multi-head attention that bins gene expression counts to jointly optimize cell and gene representations^37^. Trained on large corpora of scRNA-seq data, Geneformer and scGPT models have shown promise in learning generalizable representations of transcriptomic profiles that can be fine-tuned to perform various downstream analysis tasks.

Provided the single-cell latent embeddings derived from the CELLxGENE census-trained scVI, Geneformer, and scGPT models, we identified the task of data integration as a point of intersection of their capabilities. We reasoned that MILK can facilitate a comprehensive but tractable analysis to assess how well the models can capture information in massive single-cell resources of scRNA-seq data. While the scVI model learns fewer parameters and its latent embeddings consist of fewer dimensions compared to the transformer model architectures under test, we hypothesized that its stronger inductive bias in modelling observed gene expression values from a statistical probability distribution may compensate for its reduced model complexity.

Only primary data were considered to avoid duplicate cells present across multiple datasets (e.g., meta-analyses). This resulted in the census containing 44,265,932 cells. An overview of the metadata variables in the CELLxGENE census indicated 369 scRNA-seq datasets, 24 assays (i.e., sequencing technologies), 678 cell types, 176 developmental stages, 108 disease contexts, 3 sex categories (male, female, and unknown), 2 suspension types (cell versus nucleus), 264 tissues, and 55 general tissues. For downstream analyses, cell type and disease annotations provided by submitting authors were utilized as biologically meaningful groups for downstream analyses. A minimum cell count cutoff of 50 was applied to the paired cell type–disease groupings. Cells were filtered if no cell type annotations were provided.

#### Methods 4.2: MILK construction and representative subsampling of Census embeddings

MILK was applied to the single-cell latent embeddings derived from scVI, Geneformer, and scGPT using a similarity threshold fixed to the 0.01 percentile. The upper limit cache size was set to 50,000 cells. The partition size, merging threshold, and upper limit cache size were set to 10,000 cells. MILK was applied with Euclidean and cosine distance metrics.

Representative subsamples of the CELLxGENE census were extracted from the resulting MILK trees according to varying sizes (target sample size: 10000, 30000, 100000, 300000, and 1000000 cells) using the following metadata labels (label-balanced subsampling as described in **Methods 2.5**): “cell_type”, “tissue_general”, “cell_type-disease” (paired by concatenating cell type and disease labels per cell), and “tissue_general-disease” (paired). MILK was then reapplied using identical parameterization to obtain a class-balanced hierarchy of the CELLxGENE census.

Random subsampling baselines were also generated according to the varying target sample sizes and metadata information.

#### Methods 4.3: UMAP visualization of representative Census embeddings

Following extraction of representative subsamples, the latent embeddings derived from the scVI, Geneformer, and scGPT models were qualitatively assessed based on their UMAP embeddings. In particular, a KNN graph (k = 50) was constructed over the single-cell latent embeddings and the corresponding distance metric that was used to reconstruct the MILK tree (Euclidean and cosine distance). The UMAP coordinates were then obtained with a `min_dist` value of 1.0 and a `spread` of 4.0. Point size was used to denote the group size of each subsampled cell.

In addition, the UMAP coordinates for the unintegrated transcriptomic profiles were computed by normalizing (`scanpy.pp.normalize_total`) and log1p-transforming (`scanpy.pp.log1p`) the raw expression counts of cells. HVGs (`scanpy.pp.highly_variable_genes`) were identified and then PCA was applied (`scanpy.pp.pca`) with 100 components. On the PCA embeddings, a KNN graph was constructed and used to obtain the UMAP coordinates using the same parameterization as mentioned above.

#### Methods 4.4: Metadata label coverage across Census MILK recursion

For cell type, dataset ID, development stage, disease, general tissue, and tissue metadata variables, label coverage was calculated as the proportion of their respective diversity captured among representative cells for each recursive iteration of MILK on the complete CELLxGENE census. This label coverage was calculated on the latent embeddings derived from each model and was plotted as a function of the number of groups, or representatives, per MILK iteration.

#### Methods 4.5: Augmented subsample information from downstream clade coverage

During the subsampling process from MILK trees, selection of internal nodes (representatives) provides a structured approach to retain information for the corresponding leaf cells that were not explicitly included in the subsample. For each subsample, its retention of cells explicitly selected or indirectly linked to a representative was quantified by taking the total sum of clade sizes in the subsample divided by the total number of cells in the CELLxGENE census (44,265,933 cells). The degree of dataset coverage was visualized for different metadata labels used during subsampling and across target sample sizes. The resulting information retention values pertaining to Euclidean distance MILK trees across models were visualized alongside random subsampling baselines, in which only information from subsampled cells were available.

Information from the associated leaf cells was incorporated as weights for downstream analyses involving the representative subsamples. In particular, the clade size corresponding to each subsampled cell was included as a weight for downstream label-based analyses (e.g., cell type counts). These weighted subsamples, referred to as “augmented”, represent a non-disruptive approach to retain information content as compared to the explicit pseudobulking of expression counts (see **Methods 2.6**), which systematically change the scale of the transcriptomic readout.

#### Methods 4.6: Cluster-based benchmark of Census-trained embedding integration

To comprehensively compare models in terms of their integrated latent embeddings of the global census, a benchmarking analysis was performed using the Leiden clustering algorithm. Metadata variables included in the CELLxGENE census were manually categorized into biological (dataset ID, donor ID, cell class, cell subclass, cell type, development stage, disease, organ, system, tissue, and general tissue) or batch (assay and suspension type) variables. The rationale for categorizing dataset ID and donor ID as biological variables was based on the observation that individual scRNA-seq datasets tended to preserve context-specific transcriptomic profiling more as the number of datasets and donors increased.

The CZI CELLxGENE ontology guide (https://github.com/chanzuckerberg/cellxgene-ontology-guide) was used to derive cell class and cell subclass information from each cell type, as well as organ and system labels for each tissue. Representative subsamples derived across all target sample sizes were clustered according to varying Leiden clustering resolutions (2^0^ to 2^10^ in exponential steps of 1), and the ARI was assessed with respect to each biological and batch metadata variable.

In addition to ARI, we assessed adjusted mutual information (AMI) and homogeneity scores of metadata labels across representative subsamples. Unlike ARI, which focuses on pairwise agreement between cell assignments to clusters, AMI is an entropy-based measure of how well clusters preserve the overall information structure of the metadata variables. Homogeneity is a measure of how specific clusters are to the metadata groups. Both AMI and homogeneity are scored from [0,1], where 0 indicates poor concordance, and 1 denotes high concordance between metadata variables and clusters.

To normalize biological variables to account for batch-related signal, a composite batch score was computed across batch variables according to the geometric mean (or arithmetic mean of log-transformed values) to avoid underflow:

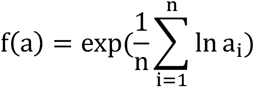

Where a corresponds to the ARI values for each batch variable. The benchmarking score for each biological variable was then normalized by taking the product of the biological score and the complement of the composite batch score per Leiden clustering resolution.

The ARI, AMI, and homogeneity scores were plotted across all Leiden clustering resolutions for MILK and random subsamples. An additional augmented subsample group was included for MILK-derived samples, in which label counts were weighted according to the clade size of each subsampled cell (see **Methods 4.5**). An ANOVA test was applied (`aov` function in the stats R package) for each ARI, AMI, and homogeneity benchmarking metric to assess the overall interaction of model and subsampling approach on the corresponding benchmarking scores. Subsequently, the Geneformer, scGPT, and scVI models were compared via paired t-test with Bonferroni correction (subsampling approach as the pairing variable), which were calculated using the `pairwise_t_test` function in the rstatix^83^ package. Similarly, subsampling approaches were compared with unpaired t-tests with Bonferroni correction. These latter comparisons were unpaired due to the fact that underlying cell subsets were distinct.

For each biological metadata variable, the Leiden clustering resolution that maximized batch-adjusted benchmarking scores was selected. The optimized label scores were then compiled to form a composite score using the geometric mean, which enabled model evaluation according to target sample size, data integration (versus unintegrated control), pairwise comparison metric (Euclidean and cosine), metadata labels used during the subsampling process, between “augmented” versus “original” subsample information content, compared to random subsampling baselines.

Notably, independent optimization of clustering resolution per metadata label maximizes performance per variable, but it comes at the expense of extensive parameter tuning and limitation in cross-interpretability between variables optimized at different clustering resolutions. The inability to jointly capture metadata variables in a shared optimal space is a limitation that can be attributed to the reliance on unstructured data encodings, which becomes increasingly problematic as the complexity and scale of datasets increase.

#### Methods 4.7: Information gain benchmark of Census MILK tree topology

Using the MILK tree topologies corresponding to each representative subsample, an entropy-based analysis was performed across metadata variables. Only subsamples derived according to a target sample size of 1 million cells and Euclidean pairwise metric were considered. Specifically, information gain (IG) was computed for each internal node in the MILK tree by subtracting the Shannon entropy of the parent clade from the conditional entropy of the children’s clades, where the condition is the branching event in the MILK tree. This can be described as follows:

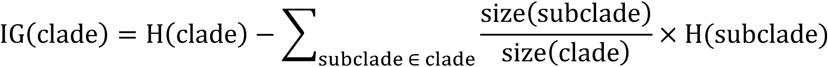

Where H() corresponds to the Shannon entropy of the metadata labels of the leaf cells captured by the clade of interest. The IG value for each clade was then normalized by the IG score of a shuffle control in which MILK tree labels were shuffled among leaf cells.

MILK tree (*T*)’s summary scores were then obtained by taking the weighted average normalized IG score with clade size weights, as this ensures equal weighting of IG scores at each level in the tree. Note that clade size refers to the number of leaf cells associated with the internal node. This can be formally defined by the following equation:

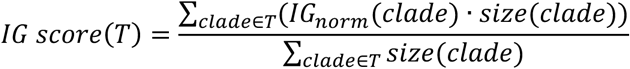

In addition, the AUC scores of normalized IG scores were calculated as a function of log-transformed clade sizes to obtain MILK tree summary scores.

IG-based analysis of MILK tree topologies was used to compare different models across biological variables, information content of subsampled cells (augmented versus original), and label information utilized during the subsampling process. In particular, this analysis was used to assess the overall flow of information explained by the tree topology, providing a more holistic and interpretable measure between variables of interest with no parameter optimization.

### Methods 5: Disease-context analysis in CELLxGENE Census MILK hierarchies

#### Methods 5.1: Disease-context visualization of the Geneformer MILK tree

The MILK tree derived from Geneformer latent embeddings at a target sample size of 1 million cells, Euclidean distance, and augmented subsample information content was visualized as a network using the `sfdp_layout` in the graph_tool Python library. Nodes were colored according to disease context labels, with dark gray representing “normal” context and colors denoting different disease contexts. Node size in the resulting network was proportional to clade size.

#### Methods 5.2: Cell type–disease dysregulation and locality metrics

To explore cell type and disease-specific associations captured by the single-cell MILK tree, a concatenated group label was formed for each cell according to the combination of cell type and disease context metadata information (i.e., “group”, referred to as *ctd*, can be a cell type–disease pairing such as “B cell | COVID-19”).

##### Methods 5.2.1: MILK dysregulation coefficient

A dysregulation coefficient (*D*) was calculated for combinations of cell type and disease labels (*ctd*) in the representative MILK tree (*T*) of the global CELLxGENE census. Specifically, for a given cell type and disease pairing (i.e., group), the count of group labels captured by the leaves associated with each clade was divided by the respective total number of group labels to obtain a relative proportion (*p_ctd_*). Subsequently, dysregulation of a specific cell type was defined as the weighted average of the absolute differences in proportions for normal and disease labels across all clades in the tree, using clade size as weights. This is shown by the following equation:

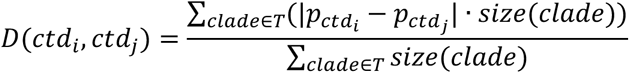

In this equation, a high dysregulation coefficient corresponds to greater separation in the MILK tree topology for the cell type in a specific disease context relative to its normal context. As such, a dysregulation coefficient of 1 indicates that the cell type exhibits clear separation in the MILK tree topology for a specific disease context compared to its “normal” state, whereas a coefficient of 0 indicates no discernible separation. Notably, this perturbation coefficient does not depend on the concentration of each subpopulation in the tree topology.

##### Methods 5.2.2: Cell type–disease locality score

Additionally, the locality of the cell type–disease group was computed as the weighted average clade size, using the relative proportion of the group of interest captured per clade as weights. This can be represented by the following equation:

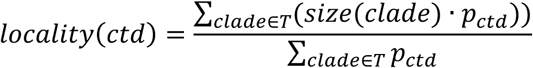

Here, the cell type–disease proportion captured by a given clade can be calculated by dividing the count of cell type–disease labels among leaf cells downstream of the clade by the total number of labels in the dataset. This provides an approximate, or characteristic, clade size value required to robustly capture the cell type–disease grouping within the global MILK tree. A locality of 1 indicates that cells in the group of interest are randomly spread throughout the tree topology and can only be sufficiently captured by the root of the tree. Small locality values indicate that the cells belonging to the group of interest exhibit minimal spread in the tree topology and can be robustly captured by smaller clade sizes.

#### Methods 5.3: Thresholded dysregulation hits and model overlap

Following the quantification of dysregulation coefficients for each cell type–disease pairing in the subsampled MILK trees, binarized dysregulation hits were defined according to different thresholds ranging from 0 to 1 (step size: 0.01). This enabled a systematic comparison of dysregulation hits across models for (augmented) subsamples. The overlap in dysregulation hits across the three models was also characterized across thresholds, stratifying hits based on their intersection across all three models, at least two models, or uniquely by their respective model.

#### Methods 5.4: Differential expression analysis of disease-context cell types

We next investigated whether the disease-mediated dysregulation coefficients per cell type were reflected in the underlying gene expression data. Specifically, each cell type in a specific disease context was evaluated relative to its normal context in terms of its dysregulation coefficient in the MILK tree and the number of differentially expressed genes in the underlying transcriptomic profiles.

For this analysis, we utilized the raw, unintegrated gene expression counts of protein-coding genes. To identify protein-coding genes from the 60350 dimensions included in the standardized CELLxGENE census, we first mapped the Ensembl gene IDs to the corresponding gene “short names” via the `mygene`^84^ Python package (e.g., “ENSG00000000003” could be mapped to tetraspanin 6, or TSPAN6). Inclusion of only features that could be mapped to a known Ensembl gene and characterized as “protein-coding” filtered the dimensionality from 60530 to 19417 genes. The raw counts were then normalized and subjected to log1p transformation.

For each cell type, a DEG analysis was performed for each disease label with respect to its “normal” or “healthy” context using the `scanpy.tl.rank_genes_groups` (method = “t-test”). A minimum cutoff requiring at least 50 cells per cell type–disease (or normal) group was applied. The resulting DEG analysis returned a t-test statistic score, log fold-change, and an adjusted *P* value for every statistically significant gene in the cell type and disease pairing.

The Pearson correlation between MILK tree dysregulation coefficients and the log-transformed number of DEGs was computed. These relationships were assessed in aggregate per cell type and disease by taking the average of respective metrics.

#### Methods 5.5: Global-versus-tissue dysregulation concordance

Given that associations between cells in the unified contextualization could arise from non-biological factors including batch effect or technical artefacts, we sought to validate whether global dysregulation hits successfully captured tissue context-specific hits. Specifically, 54 general tissue subsets were extracted from the complete CELLxGENE census and MILK was applied to the latent embeddings of each model using the same parameterization as **Methods 4.2**. Representative subsamples were then extracted from the MILK trees with label-balanced subsampling using paired cell type–disease label information according to a target sample size of 1 million cells. If a general tissue subset had fewer cells than the target size, the complete dataset was sampled. Dysregulation coefficients were independently calculated for each tissue subset, and the hits identified across thresholds were treated as “ground truth”, or context-specific hits.

In parallel, we evaluated a Leiden clustering-based strategy of identifying dysregulation hits between global and tissue subsets obtained using MILK. Leiden clustering resolution was optimized across a range of values based on a composite ARI score defined as the geometric mean across ARI scores of all biological variables and the complement of ARI scores for batch variables (see **Methods 4.6**). A disease-associated dysregulation coefficient was then calculated for each cell type by dividing the count of normal and disease-context cells in disjoint clusters by the total count of normal and disease cells for the cell type of interest. The Leiden clustering resolution that maximized the composite ARI score was used for each tissue dataset. This coefficient is based on the rationale that if a given cell type’s gene expression profile is altered in the context of a given disease, then its cells will be more likely to localize in different Leiden clusters than in its normal state.

To compare Geneformer, scGPT, and scVI models, as well as MILK (original and augmented) against Leiden-based dysregulation scoring methods, the Spearman correlation of dysregulation coefficients between shared cell type–disease groups was evaluated for each tissue MILK tree and the global MILK tree. The dysregulation hits identified across thresholds for each MILK tree were further quantified in terms of hits only observed in the global context, only in tissue context, or those shared in both global and tissue contexts. The AUC of dysregulation hits as a function of the total number of hits (union of hits between tissue and global datasets) was calculated using the MESS^81^ package in R.

#### Methods 5.6: Clade-level transcriptomic analysis of representative dysregulation cases

Following calculation of dysregulation coefficients across all cell type–disease groups in the Geneformer-based MILK tree (target size of 1 million cells, cell type–disease label-balanced subsampling, with augmented subsampling information content), we selected three groups spanning the range of dysregulation scores: fibroblasts in Plasmodium malariae malaria disease context (high), neurons in Alzheimer’s disease context (intermediate), and medium spiny neurons in opiate dependence context (low). For each group, we identified the smallest MILK clade(s) capturing a majority of the cell type population in normal and disease contexts. The high-dysregulation case resulted in identification of two fibroblast MILK clades (one for normal context and one for disease context), while a single MILK clade captured both normal and disease contexts for the intermediate and low dysregulation cases.

For all cell type–disease groups captured by the MILK clades containing at least 100 cells, we computed an aggregate transcriptomic profile analysis from processed (non-integrated) gene expression counts by averaging expression across all genes per group. We then calculated the pairwise Pearson correlation of group profiles. Hierarchical clustering was then applied to identify clusters of distinct subpopulations captured by the MILK clades, which revealed the contextual shift MILK clades capture as a cell type moves from normal to disease state.

Finally, we used volcano plots to identify differentially expressed genes between normal and disease contexts, defining significance as an absolute log-fold change greater than or equal to 1 as well as an adjusted *P* value less than or equal to 0.01.

#### Methods 5.7: Pairwise association matrix of cell type–disease groups

To summarize group associations captured by the single-cell MILK tree, a pairwise matrix capturing the co-localization of groups in the MILK tree topology was computed. The relative proportion (*p*) and purity (*purity*) values of all group labels were initially computed across all clades. Relative proportion values per group label and clade were defined in **Methods 5.2.1**. Purity for a given clade and group label of interest can be defined as the count of group labels in the downstream leaves of the clade divided by the total number of leaves in the clade (clade size).

A pairwise association matrix of group labels was then defined as the weighted average clade size required to robustly capture both group subpopulations in the MILK tree:

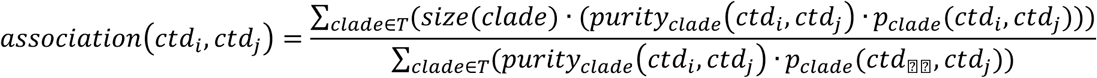

For a given pair of groups, (*ctd_i_*, *ctd_i_*), only clades that captured both group subpopulations according to a relative proportion threshold of at least 0.25 were considered. This threshold was applied to filter noise and focus on clades capturing sufficient proportions of both subpopulations. A weighted average clade size was then computed over these clades, where the weight corresponded to the product of the joint relative proportion (clade-specific relative proportion of each group) multiplied and the joint purity value (clade-specific purity of each group).

The resulting matrix was scaled (min-max scaling) and diagonals were set to 0. Smaller scaled clade sizes (e.g., 0) correspond to stronger associations in the global MILK topology, whereas larger clade sizes (e.g., 1) denote weaker global associations. Unweighted Pair Group Method with Arithmetic mean (UPGMA) hierarchical clustering was applied to the pairwise cell type–disease association matrix, and clusters of higher-level group labels were identified based on the tree depth in the hierarchical tree. Disease ontology labels that mapped the specific diseases to standardized disease classes were obtained from a large-language model (ChatGPT, OpenAI). These tree depth-based clusters aggregated by cell class and disease class were subjected to the Sinkhorn-Knopp matrix scaling prior to visualization.

#### Methods 5.8: Depth-dependent dispersion of cell type–disease association clusters

Given the depth-based clusters derived from the UPGMA tree of the cell type–disease groups, we initially characterized how dispersed cell types were across clustering depths. In particular, the total number of clusters each cell type occurred in across all disease contexts was calculated across depths. The total number of clusters (cluster dispersion) was also divided by the total number of disease contexts it was captured in to obtain a normalized fraction of cell type dispersion. The AUC values were calculated (`numpy.trapz` function in Python) over the normalized cluster dispersion scores as a function of UPGMA tree depth to summarize cell type dispersion in the MILK tree.

Next, for each cell type, we assessed whether its disease-context label diverged from its normal context label across UPGMA tree depth clusters. At each depth, we identified the cluster containing the cell type’s normal-context group and computed the fraction of that cell type’s disease-context groups located in different clusters. This provides a depth-dependent score of the dysregulated fraction for each cell type in the pairwise cell type–disease co-localization matrix derived from the MILK tree. Dysregulation fractions were independently averaged across cell types within each cell subclass (higher-level classification) and tissue, at each depth. The resulting cell type dysregulation fractions per cell subclass and tissue were ranked by the earliest tree depth at which mean dysregulation reached 1 (i.e., full divergence from normal across all disease contexts).

#### Methods 5.9: Disease association network from shared cell type dysregulation

From the disease-mediated cell type dysregulation coefficients derived from the MILK tree topology, we constructed a disease-disease association matrix. Specifically, for a pair of diseases, we computed the Pearson correlation between dysregulation magnitudes across their shared cell types. Only diseases sharing at least two cell types were compared. While such correlations are inherently biased by the availability of data for cell type and disease combinations, we hypothesized that they could still provide meaningful insights into the coordination of disease perturbations of cell states represented in the CELLxGENE census.

Pairs of diseases exhibiting strong correlation of dysregulation coefficients (Pearson correlation greater than or equal to 0.5) were used to define edge associations. The resulting network was visualized using the `arf_layout` (`d` = 100, `max_iter` = 5000) in the `graph_tool` Python library. Node colors corresponded to the disease class labels, and boundaries for these higher-level classifications were manually annotated.

#### Methods 5.10: Context-agnostic cell type association network and GSEA

Given the aggregated pairwise association matrix between paired cell type and disease labels derived from the global MILK tree (outlined in **Methods 5.6**), we also extracted a network of cell type-cell type associations. The association between a pair of cell types was defined as the median normalized clade size value of a pair of cell types regardless of different contexts. To focus on the strongest pairwise associations across the 630 cell types, we applied a threshold to select only the 1^st^ percentile of normalized clade size values (corresponding to a normalized clade size of 0.1040), which resulted in a cell type adjacency matrix containing 1831 edges.

Leiden clustering was applied to this binarized adjacency matrix using the leidenalg implementation in Python, with a resolution parameter of 1.0 and the “RBConfigurationVertexPartition” partition type. This resulted in 13 Leiden clusters with at least two cell types. This network of these cell type associations was visualized using the `arf_layout` (`d` = 100, `max_iter` = 5000) in the graph_tool Python library.

A gene set enrichment analysis (GSEA) was performed for the 13 Leiden clusters. The raw gene expression data was processed as described in **Methods 5.4**, and mitochondrial (“MT-“) and ribosomal (“RPS” and “RPL”) transcripts were filtered. DEGs were identified in the clusters using the `scanpy.tl.rank_gene_groups` function based on the Wilcoxon rank-sum method. The resulting genes were then ranked (ties randomly decided) by their differential expression score and used as input for the GSEA using the MSigDB Hallmark 2020 database^85^ provided in the GSEApy Python library^86^. This analysis was performed with 1000 permutations, and gene sets were retained if they exhibited a false discovery rate (FDR) q-value < 0.01 and a positive normalized enrichment score. The top GSEA terms for each cluster was manually annotated on the resulting cell type association networks.

#### Methods 5.11: Normal-context cell type label coherence and refinement candidates

To focus solely on the robustness of cell type labels to potentially identify candidate labels for refinement, we extracted the MILK subtree corresponding to cells captured in normal contexts (i.e., excluding cells in any disease states). Disease-context cells were excluded due to the potential confounding effects of cell perturbation in calculating MILK tree co-localization at the cell type level. We conducted a purity analysis per cell type, where for each clade in the normal-context MILK tree, the cell type purity scores were individually calculated (see **Methods 5.7**) and summarized as the AUC as a function of the log-transformed clade size. This provided an overall distribution of whether cell types exhibited relatively concentrated or dispersed localization in the normal-context MILK tree, which could be used to identify isolated candidate labels for refinement.

We extended this analysis by computing the joint purity scores of all cell type pairs across all clades in the normal-context MILK tree. Specifically, joint purity was defined as the product of cell type purity scores per clade, and overall co-localization scores between cell types were defined as the weighted average of joint purity scores, using clade size as weights. This identified cell type pairs that co-localized across clades in the MILK tree that could potentially indicate mixed cell subpopulations that could benefit from data-driven label refinement. HDBSCAN clustering was applied to the joint purity matrix of cell types, and clusters comprised of at least 2 cell types were identified. We hypothesize that such clusters of co-localized purity scores could be explained by a continuum of cell states defined by shared transcriptional programs or as potential candidates for cell type annotation refinement using the MILK tree structure.

#### Methods 5.12: Disease-context distinctness relative to normal cellular state space

We next assessed whether cell type–disease groups occupied distinct regions in the MILK tree, or whether the altered cell state exists in the space defined by normal-context cells pooled across all cell types. For each clade in the MILK tree, we computed the difference in purity value (see **Methods 5.7**) of each cell type– disease group minus the purity value of the pooled normal-context population (i.e., normal-context cells from all cell types, treated as a single reference group). We then computed the weighted average of these per-clade differences in purity scores across all clades in the tree, using clade size as weights, to obtain a purity distinctness score for each cell type–disease group. Positive values indicate that the group occupies relatively distinct clades in the MILK tree, whereas negative values indicate that most of the group’s tree localization is encompassed by the pooled normal-context reference cell space. We further stratified the overall distribution of cell type–disease purity distinctness scores at the cell type- and disease-level, identifying metadata features associated with more distinct cellular states.

### Methods 6: Cross-species MILK analysis of cell type and species organization

#### Methods 6.1: Cross-species UCE atlas data acquisition

The Universal Cell Embedding (UCE) foundation model was used to create an Integrated Mega-scale Atlas (IMA) of single-cell embeddings. Comprised of transcriptomic profiles from approximately 36 million cells across 8 species, it enabled a comprehensive analysis of cell state and tissue information in the context of a unified embedding. The scientific names for the included species are as follows: human (*Homo sapiens*), mouse (*Mus musculus*), mouse lemur (*Microcebus murinus*), zebrafish (*Danio rerio*), pig (*Sus scrofa*), rhesus macaque (*Macaca mulatta*), crab-eating macaque (*Macaca fascicularis*), and western-clawed frog (*Xenopus tropicalis*).

A representative subsample of the IMA corpus provided in the preprint was obtained (https://drive.google.com/drive/folders/1f63fh0ykgEhCrkd_EVvIootBw7LYDVI7). The downloaded dataset included the latent embeddings (1280 dimensions) for 2,969,114 cells. In addition, cell type, coarse cell type, tissue, dataset ID, and species metadata label information were provided.

#### Methods 6.2: MILK construction and subsampling of UCE embeddings

MILK was applied to the latent embeddings of the representative IMA dataset with the following parameterization: Euclidean distance as the pairwise metric, a similarity threshold set to the 0.01 percentile, a merge threshold of 50000, a partition size of 10000, and a cache size limit of 10000 cells.

Computationally tractable subsamples (target sample sizes: 100000, 300000, and 1000000 cells) were then extracted from the MILK tree using cell type-species, coarse cell type-species, and tissue-species paired metadata label information. A MILK tree was then reconstructed using the same parameterization over the tractable subsamples of the IMA corpus.

#### Methods 6.3: Collapsed visualization of the cross-species MILK tree

To visualize the distribution of coarse cell type and species labels in the resulting MILK tree, the single-cell hierarchy was collapsed via a clade pruning strategy based on the relative label abundances. For each clade, a sorted list of relative proportions of coarse cell type–species labels was initially calculated, from which a dominance ratio was defined by dividing the highest enrichment value by the second-highest enrichment value. If this ratio was greater than a threshold (parameter set to infinity, or a pure clade), the clade was pruned (collapse all downstream information into the current node) and defined as enriched for the top label. If the ratio was not met, then the process would be recursively applied to its subclades. Additionally, if the highest enrichment value fell below a cutoff threshold (parameter set to 0.10), then that path was pruned from the tree due to no significant label enrichment. This collapsing procedure was performed in a breadth-first traversal of the single-cell MILK trees to obtain a collapsed tree enriched for coarse cell type–species labels. The collapsed tree was then visualized using the circular dendrogram layout provided by igraph^87^ and ggraph R packages.

#### Methods 6.4: Reference species phylogeny from NCBI Taxonomy

The phyloT software^88^ (https://phylot.biobyte.de/) was used to obtain the phylogenetic tree corresponding to the 8 species included in the IMA corpus. The scientific names (*Macaca mulatta*, *Macaca fascicularis*, *Homo sapiens*, *Microcebus murinus*, *Mus musculus*, *Sus scrofa*, *Xenopus tropicalis*, and *Danio rerio*) were provided as NCBI tree elements, and the evolutionary tree was generated based on the NCBI Taxonomy database^89^ (https://www.ncbi.nlm.nih.gov/guide/taxonomy/).

#### Methods 6.5: Cell type–species capture-scale analysis

To investigate the relationship between cell type and species signal in the global hierarchy, the weighted average clade size (*S*) required to robustly capture each label group was computed. Specifically, for a given label (*l*) and threshold (*t*) combination (0 to 1 in steps of 0.01), the weighted average sizes of clades required to capture the label in the MILK topology were computed.

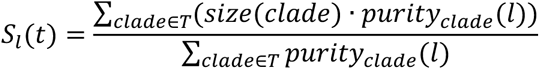

According to the label-capture threshold, all clades capturing at least that relative proportion (**Methods 5.2.1**) of the label of interest were identified, and the purity (see **Methods 5.7**) of the label within the clade was utilized as weights. This process was performed across thresholds for all coarse cell type–species (paired) groups. The characteristic clade sizes to robustly capture the coarse cell type–species populations as a function of label capture threshold were plotted per cell type, with a curve denoting each species. These values were further summarized by computing the AUC values of clade sizes as a function of threshold.

#### Methods 6.6: Information gain analysis of cell type and species signals

IG scores were calculated for each clade in the MILK tree (target sample size of 1 million cells) for the following metadata label combinations: coarse cell type, species, and coarse cell type–species (paired). The resulting values were normalized by subtracting the IG scores per observed clade from the IG values for corresponding controls with shuffled metadata labels (n = 3). Following normalization, a cumulative distribution function (CDF) curve was calculated using the mean normalized IG scores at each observed clade size as a function of the log-transformed clade size for each label. The resulting CDF characterizes whether informative signal for a given metadata label concentrates in small clades or large clades.

A central question given the unified MILK hierarchy comprised of multiple species was whether its hierarchical structure organizes cells primarily by cell type, species, or some mixture of both. To address this, we quantified cell type versus species signal dominance across the MILK tree using the normalized IG values per clade. Given the clade-specific normalized IG scores, we next stratified IG scores according to cell type-specific regions in the MILK tree. Similar to **Methods 6.5**, for each coarse cell type–species group of interest, we performed a threshold sweep (0 to 1, in steps of 0.01) to select clades in which the group’s relative proportion (see **Methods 5.2.1**) met or exceeded the threshold. Among the clades meeting each threshold, we computed a weighted average normalized IG score independently for cell type and species labels, using the group’s clade purity values (see **Methods 5.7**) as weights. This yielded, for each group, a weighted average IG curve for both labels as a function of threshold.

We then computed the AUC of each curve, producing a cell type-specific and species-specific AUC value for each cell type–species group. AUC values were marginalized by averaging across species within each cell type, resulting in cell type-specific AUC scores for both cell type and species labels. Comparing these two scores allows classification of each cell type as predominantly cell type-associated or species-associated in its MILK clade organization. Notably, because this analysis is restricted to clades enriched for the cell type of interest, the comparison indicates only the relative degree of cell type versus species organization within the local regions capturing that cell type, and thus cannot establish global signal dominance across the full MILK tree.

#### Methods 6.7: Phylogenetic concordance of cell type-specific species trees

The MILK tree obtained with a target sample size of 1 million cells was utilized to test the agreement between cell type-specific species relationships and the evolutionary tree. Specifically, the MILK tree was aggregated into a pairwise association matrix of cell type–species groups, as described in **Methods 5.7**.

Following extraction of the pairwise association matrix per cell type, the UPGMA algorithm was applied to obtain the tree structure of the available species relationships. This step was performed using the `upgma` function in the phangorn R package^90^. The UPGMA reconstructed species tree was subsequently compared against the evolutionary tree via a normalized triplet distance score using the `TripletDistance` function in the Quartet R package^91^. This evaluation method exhaustively samples all possible triplets of species subtrees in the UPGMA tree and evolutionary tree and then scores the fraction of subtree topology matches over the total number of subtree topologies. The resulting complement of the triplets’ distance was taken (i.e., 1 – normalized triplets distance value) to obtain a score where 1 corresponds to perfect agreement, and 0 denotes complete disagreement.

A bootstrapping confidence score was also provided for each cell type agreement score. Specifically, 100 orthogonal partitions of each coarse cell type–species subpopulation were extracted from the MILK tree. On each bootstrap replicate, a pairwise association matrix of cell type–species groups was calculated (see **Methods 5.7**), after which, for each cell type subtree, the Spearman correlation was computed over the non-diagonal species relationships between the bootstrap and the complete pairwise association matrix. The average Spearman correlation across 100 bootstrap replicates defined the overall bootstrap score.

---

**Supplementary Fig. 1.1.**
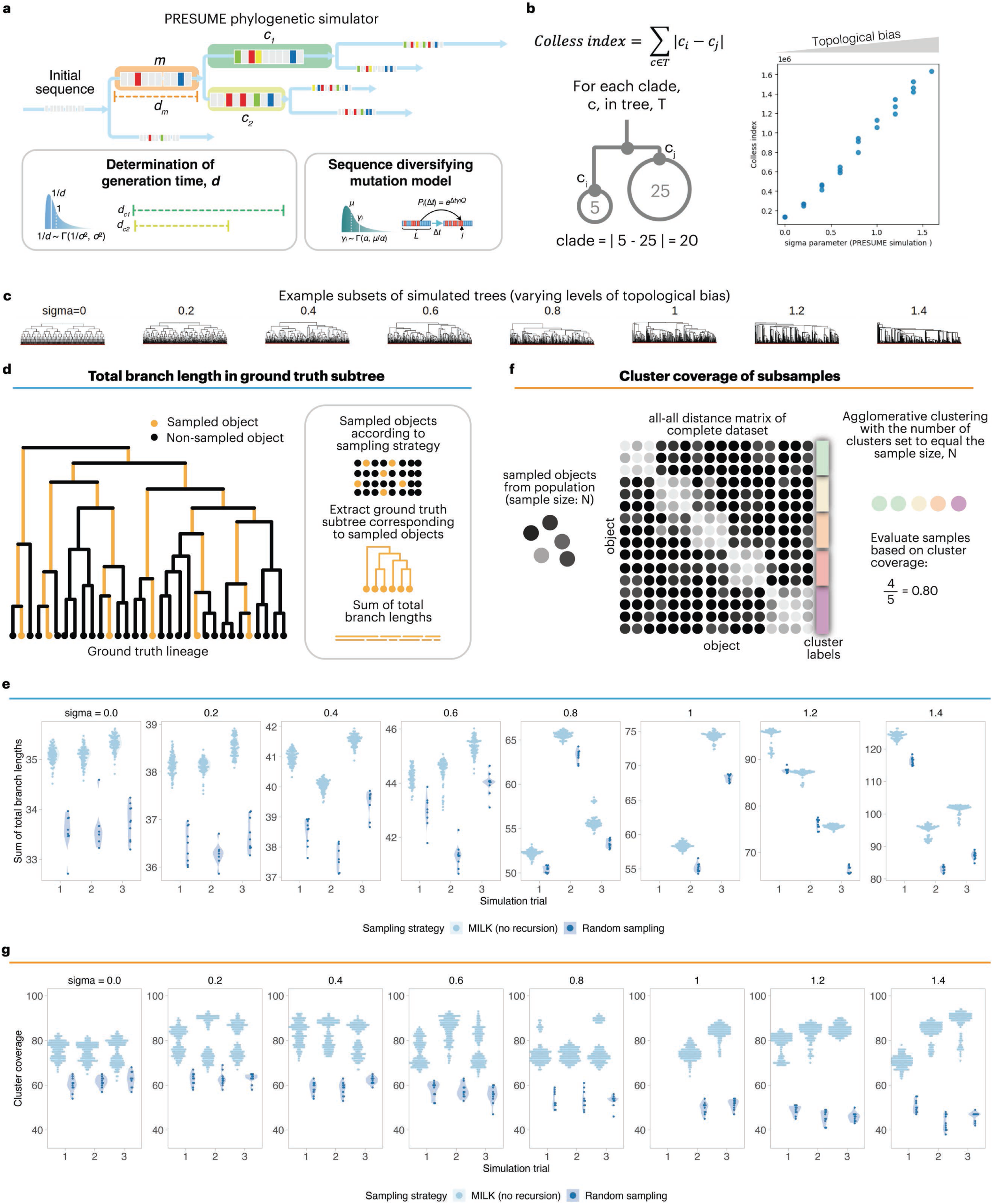
MILK-based representative subsampling of simulated phylogenetic nucleotide sequences. **a,** Phylogenetic sequence simulation. Simulated phylogenetic datasets were generated using PRESUME to jointly produce bifurcating ground-truth trees and nucleotide sequences evolving under a GTR-GAMMA model. Trees were simulated with a target size of 100,000 leaves across increasing values of the sigma parameter (0.0–1.4; n = 3 simulation replicates per value), which controls the degree of topological imbalance in the population. The final number of simulated leaves varied across datasets, particularly at higher sigma values; the lineage for sigma = 1.0, replicate 1, was omitted because of PRESUME simulation failure. **b,** Topological imbalance validation. The Colless index, defined as the sum of absolute differences in child-clade sizes across all internal nodes, was used to quantify topological imbalance in each simulated tree. **c,** Representative simulated subtrees. Example ground-truth subtrees containing 100 leaves illustrate increasing topological imbalance across sigma values. **d,** Tree-based subsampling metric. MILK was applied without recursion to nucleotide sequences using a 5th-percentile Hamming-distance threshold to obtain subsamples of approximately 100 sequences. For each MILK-derived or randomly sampled subset, the corresponding subtree was extracted from the ground-truth phylogeny, branch lengths were optimized with RAxML-NG under a GTR-GAMMA model, and the total branch length was used as a measure of phylogenetic coverage. **e,** Total branch-length coverage. Distributions of total branch lengths for MILK-derived subsamples (n = 10) and random subsamples (n = 100) across sigma values and simulation replicates. **f,** Cluster-based subsampling metric. Pairwise Hamming distances were computed across all simulated nucleotide sequences, and agglomerative clustering was performed with the number of clusters set to the subsample size. Subsample representativeness was quantified as the fraction of clusters represented by each subsample. **g,** Cluster coverage. Cluster coverage for MILK-derived subsamples (n = 10) and random subsamples (n = 100) across sigma values and simulation replicates.

**Supplementary Fig. 2.1.**
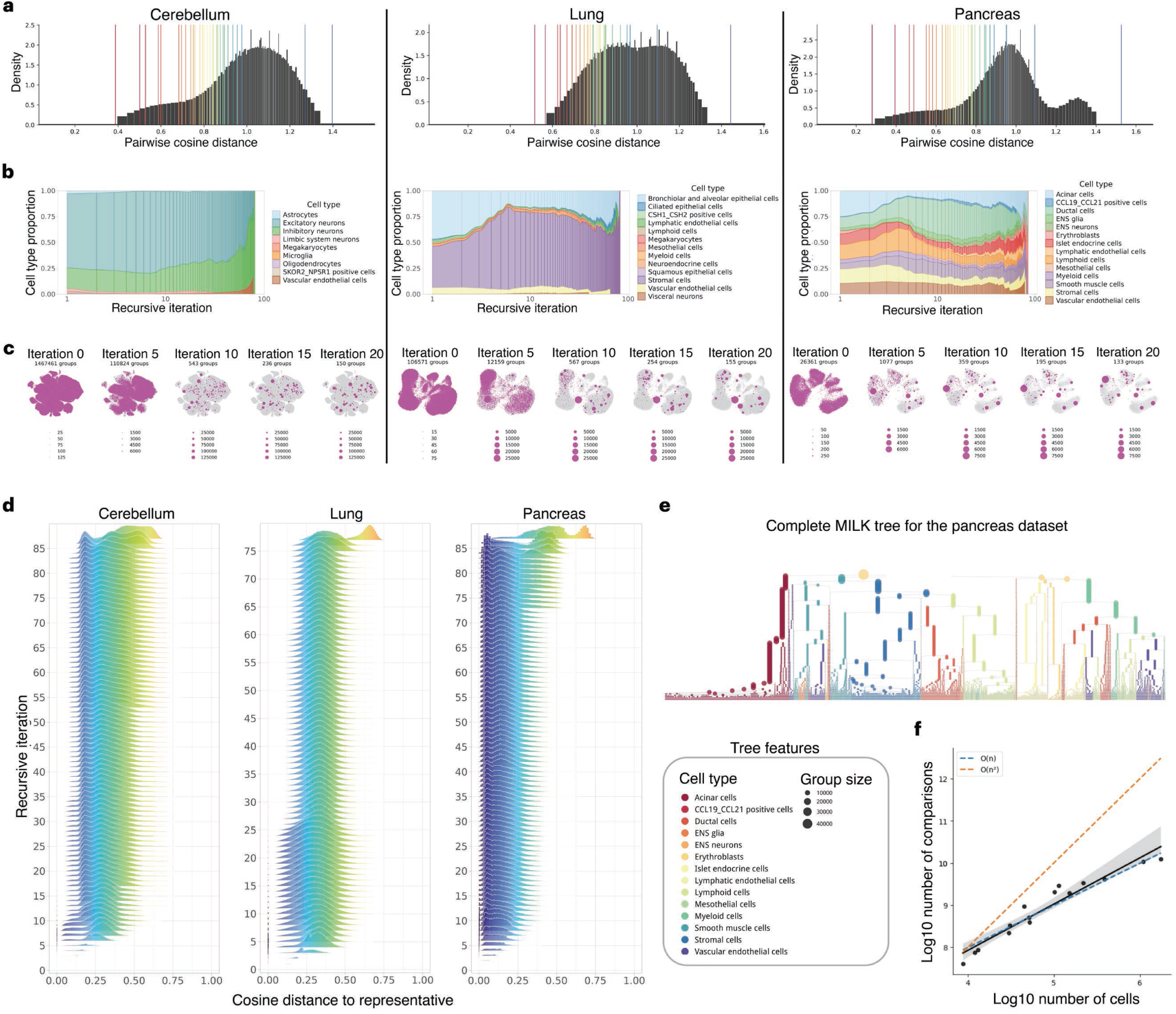
Information progression across recursive MILK iterations. Overview of the recursive MILK grouping process for three representative human fetal organ datasets: cerebellum, lung and pancreas. **a,** Dynamic similarity thresholds. Approximate distributions of pairwise cosine distances in the initial MILK iteration for the cerebellum, lung and pancreas datasets. Vertical lines indicate the similarity thresholds obtained by recursively applying the fixed 0.1 percentile threshold across MILK iterations, illustrating how the effective similarity threshold progressively changes as the input population becomes coarser. **b,** Cell type composition across recursion. Relative cell type composition of group representatives across recursive MILK iterations for the cerebellum, lung and pancreas datasets. **c,** Representative distributions across resolutions. Group representatives, defined as medoid cells, are shown in magenta within UMAP embeddings of the corresponding complete organ datasets in light gray across selected recursive iterations. Point size denotes group size. **d,** Member-to-representative distance distributions. Distributions of cosine distances between group members and their corresponding representatives across recursive MILK iterations. Distance values were pooled across groups at each iteration. **e,** Pancreas MILK tree. Complete MILK tree for the pancreas dataset. The y-axis corresponds to recursive MILK iterations, node color denotes cell type, and point size denotes group size. **f,** Empirical scaling of pairwise comparisons. Total number of object comparisons required to reconstruct each organ MILK tree as a function of the number of input cells. Each point represents one organ dataset. Axes are log10-transformed, the fitted line indicates linear regression, and dotted lines indicate theoretical linear and quadratic scaling.

**Supplementary Fig. 2.2.**
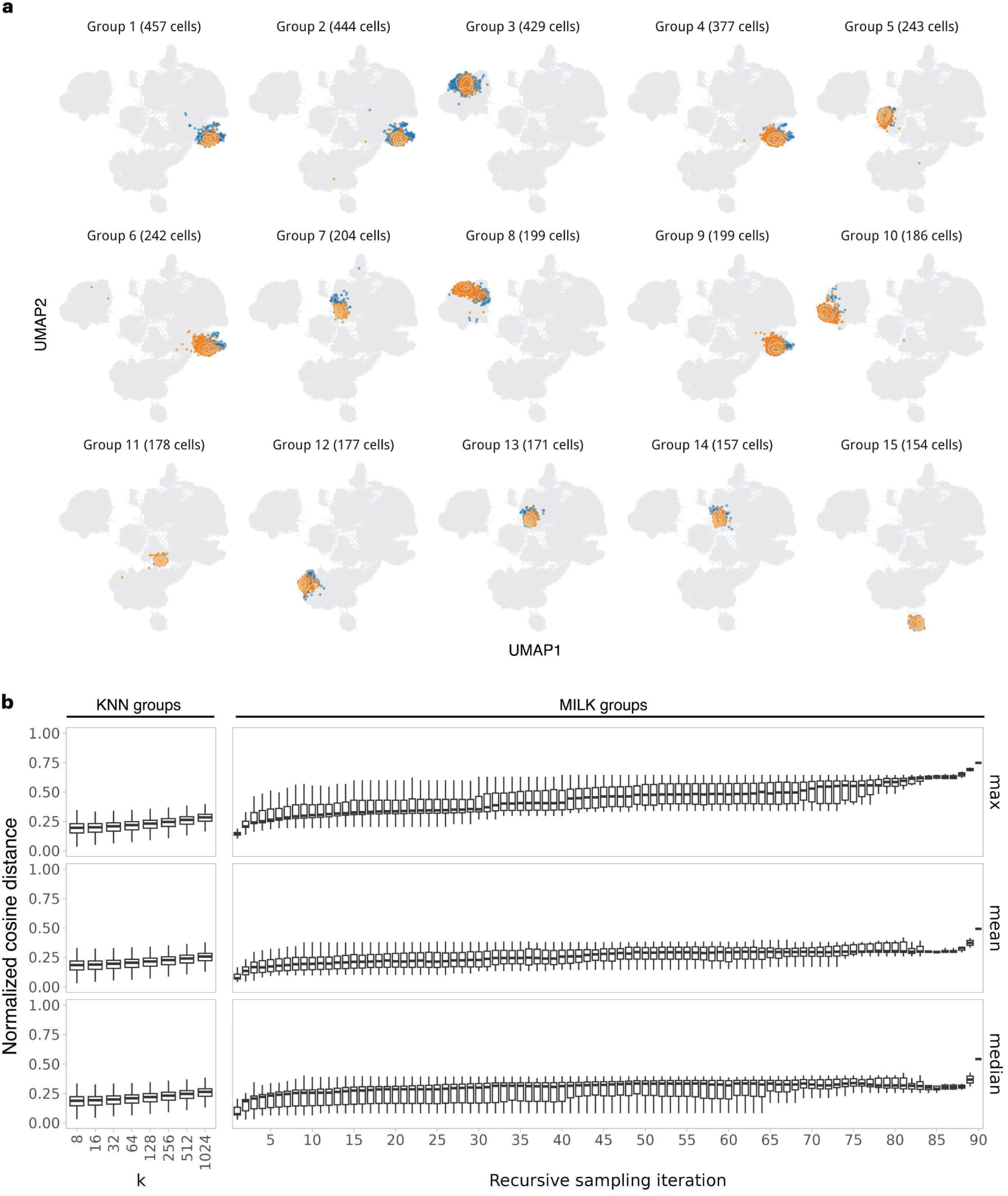
Nearest-neighbor structure of MILK groups in the eye dataset. **a,** Nearest-neighbor overlap of MILK groups. The 15 largest MILK groups identified in the eye organ dataset using a 1^st^ percentile similarity threshold without recursion are shown in orange within the UMAP embedding of the complete dataset in light gray. For each MILK group, the corresponding representative cell was used to identify its 50 nearest neighbors, shown in blue. Contour lines indicate the density of MILK group members and nearest-neighbor cells in their respective colors. The UMAP embedding was constructed from a k-nearest-neighbor graph using cosine distance with k = 50. **b,** Member-to-representative distances across MILK recursions. Normalized cosine distances between group members and their corresponding representatives were summarized for all groups identified across the recursive MILK process. Boxplots show the maximum, mean, and median member-to-representative distances for MILK groups at each recursive iteration. Corresponding distance summaries for conventional k-nearest-neighbor groups are shown across k values of 8, 16, 32, 64, 128, 256, 512 and 1,024.

**Supplementary Fig. 2.3.**
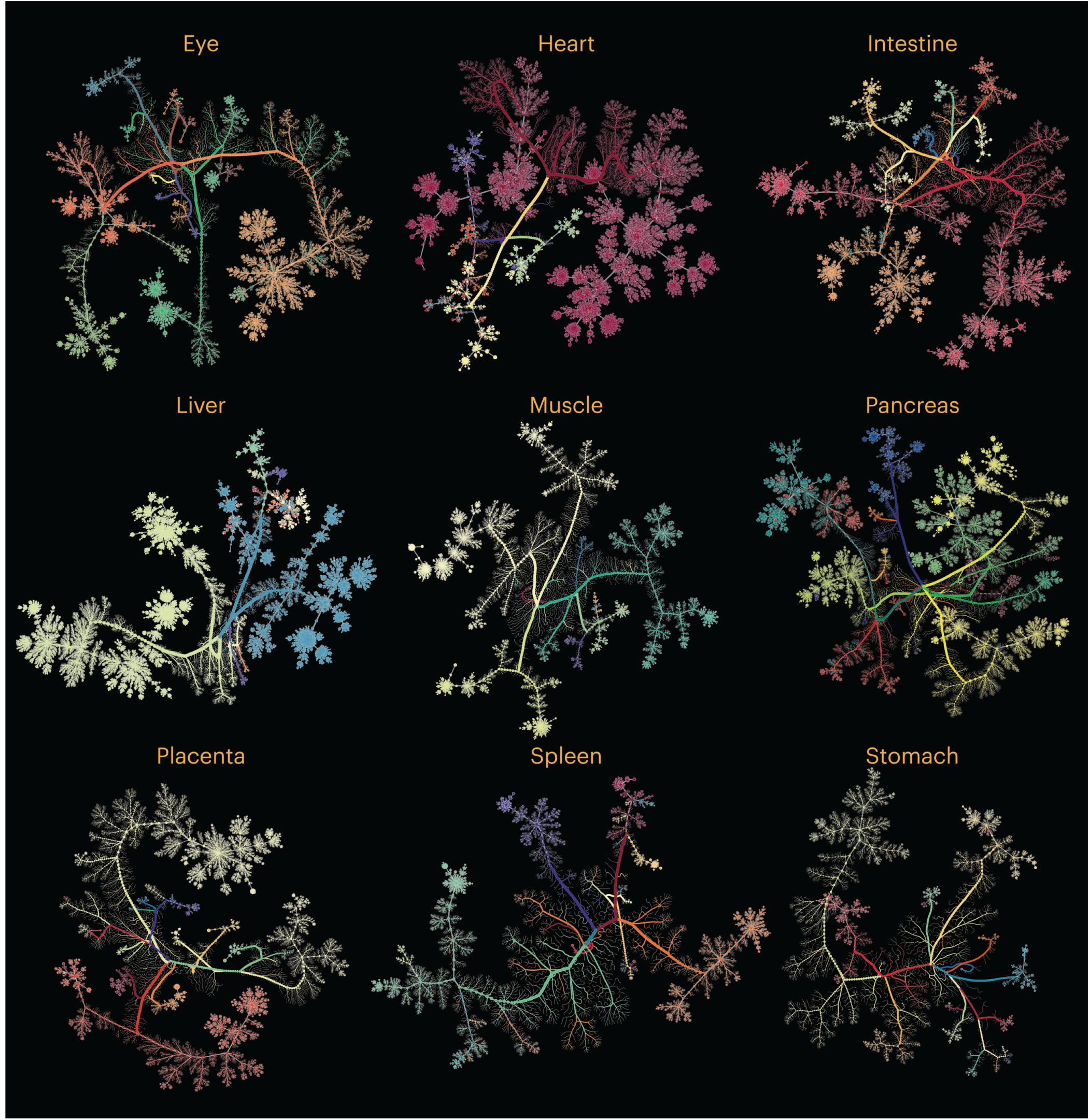
Force-directed layouts of complete organ MILK trees. Scalable force-directed network layouts of complete MILK trees constructed for nine human fetal organ datasets: eye, heart, intestine, liver, muscle, pancreas, placenta, spleen and stomach. Each node represents a MILK group, edges indicate recursive grouping relationships, and node color denotes cell type annotation.

**Supplementary Fig. 2.4.**
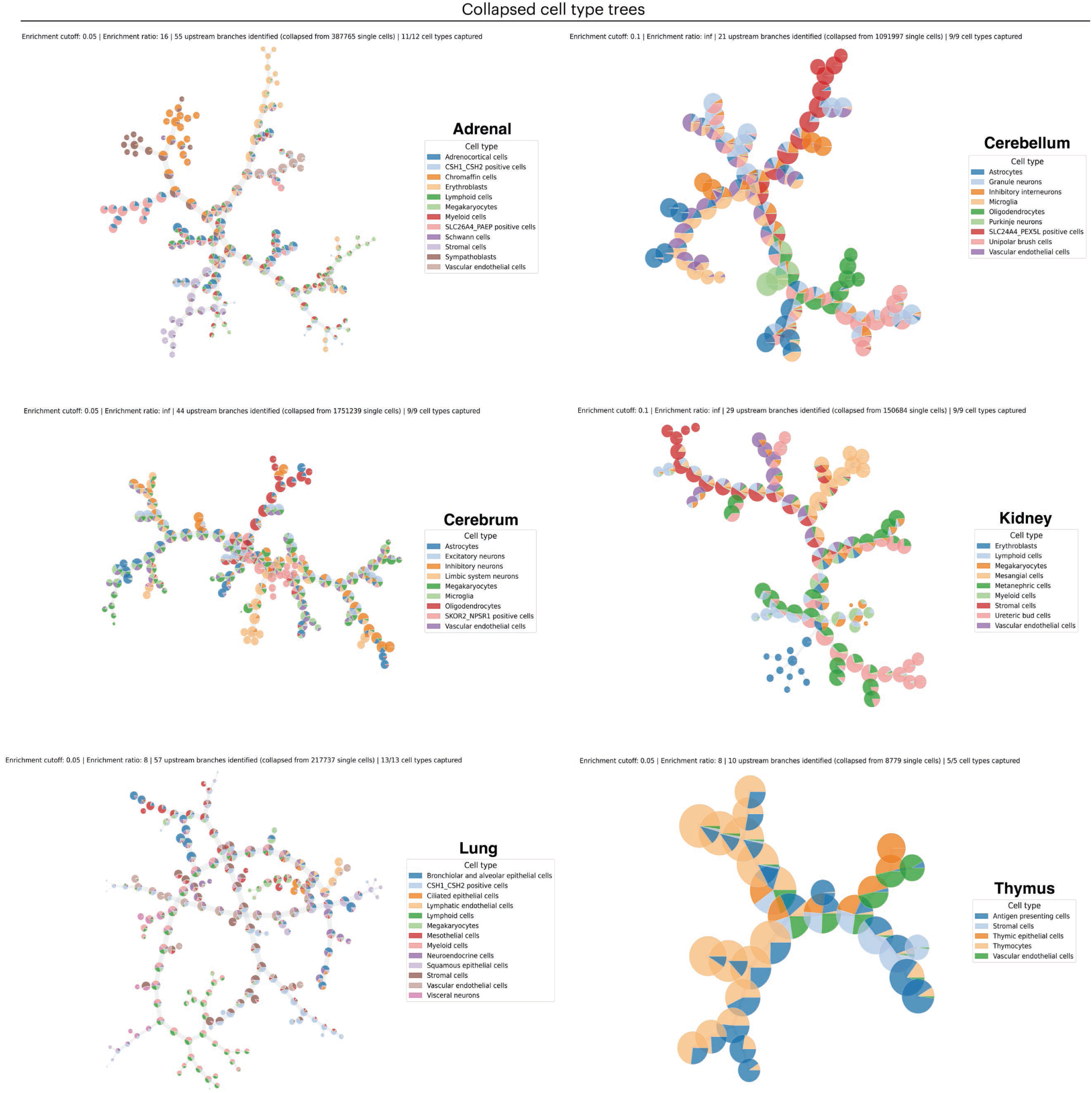
Cell type-collapsed MILK trees. Complete MILK trees collapsed into cell type-enriched network representations for six human fetal organ datasets: adrenal, cerebellum, cerebrum, kidney, lung, and thymus. For each clade, the relative proportion of each cell type was calculated as a cell type enrichment score, and the ratio between the largest and second-largest enrichment scores was used as an enrichment ratio. Clades with enrichment ratios greater than or equal to a specified threshold were collapsed into terminal nodes representing the dominant cell type; clades below this threshold were retained and recursively evaluated through their child clades. Branches were pruned when the highest cell type enrichment score fell below a specified cutoff, indicating insufficient enrichment for any cell type label. Collapsed trees were generated across enrichment-ratio thresholds of 2, 4, 8, 16, and infinity and enrichment-score cutoffs of 0.05, 0.10, and 0.25, after which one representative collapsed tree was manually selected for each organ. Pie charts denote the relative cell type composition captured by each collapsed node.

**Supplementary Fig. 2.5.**
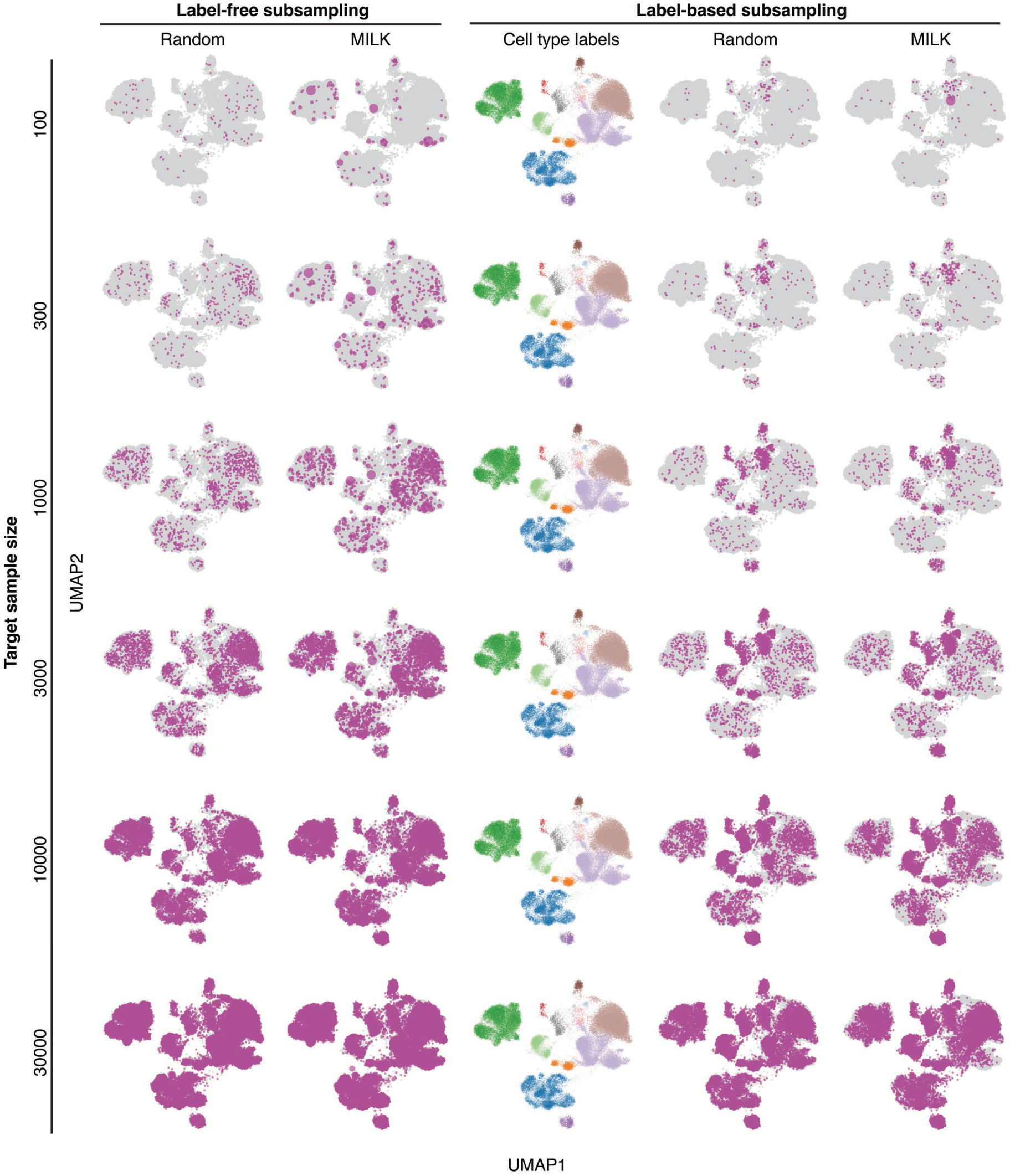
Representative subsampling strategies in the eye dataset. Qualitative comparison of representative subsampling strategies applied to the eye organ MILK tree. Subsampled cells are shown in magenta within the UMAP embedding of the complete eye dataset in light gray. Rows correspond to target sample sizes of 100, 300, 1,000, 3,000, 10,000 and 30,000 cells. Columns show label-free random subsampling, label-free MILK subsampling, cell type annotations, label-balanced random subsampling and label-balanced MILK subsampling. In label-free random subsampling, cells were sampled uniformly at random without replacement from the complete dataset. In label-free MILK subsampling, non-overlapping clades were selected from the MILK hierarchy based only on tree topology, prioritizing structurally distinct regions until the target sample size was reached. In label-balanced random subsampling, cells were sampled within each cell type according to a per-label cutoff defined by the target sample size divided by the number of cell type labels, with all cells included for cell types below the cutoff. In label-balanced MILK subsampling, the same per-label cutoff was applied, but cells were merged and selected through the MILK hierarchy only when clades were consistent with the corresponding cell type label, thereby preserving label balance while requiring support from the MILK topology. If the total number of cells in an organ dataset was smaller than the target sample size, the complete organ dataset was retained.

**Supplementary Fig. 2.6.**
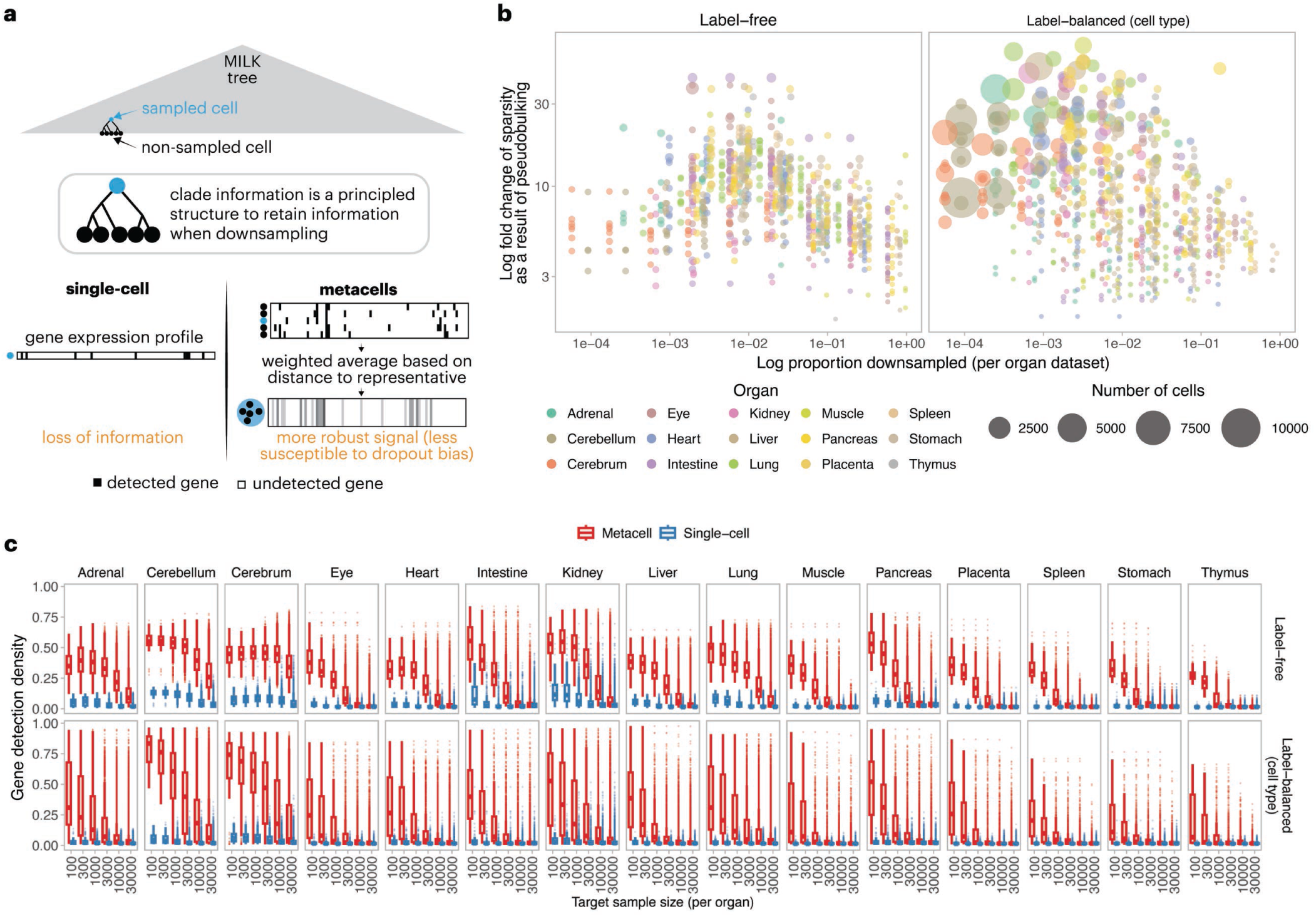
MILK-derived metacells and gene-detection density. **a,** MILK-based metacell construction. Representative cells selected from a MILK tree can correspond to internal clades containing non-sampled cells. For each sampled representative, a metacell expression profile was generated by aggregating the transcriptomic profiles of cells within its associated clade using distance-based weights, such that cells closer to the representative contributed more strongly to the resulting profile. **b,** Increased gene-detection density in MILK-derived metacells. For label-balanced and label-free MILK subsampling across all 15 organ datasets, gene-detection density was quantified as the fraction of non-zero entries in each expression profile. The log-transformed fold change in gene-detection density for metacells relative to their corresponding single-cell representatives is plotted as a function of the log-transformed proportion downsampled per organ dataset. Point size denotes the number of cells represented by each metacell, and color denotes organ identity. **c,** Organ-specific gene-detection density. Gene-detection density for metacell and single-cell profiles across label-balanced and label-free MILK subsampling strategies. Boxplots summarize distributions across target sample sizes for each organ dataset.

**Supplementary Fig. 2.7.**
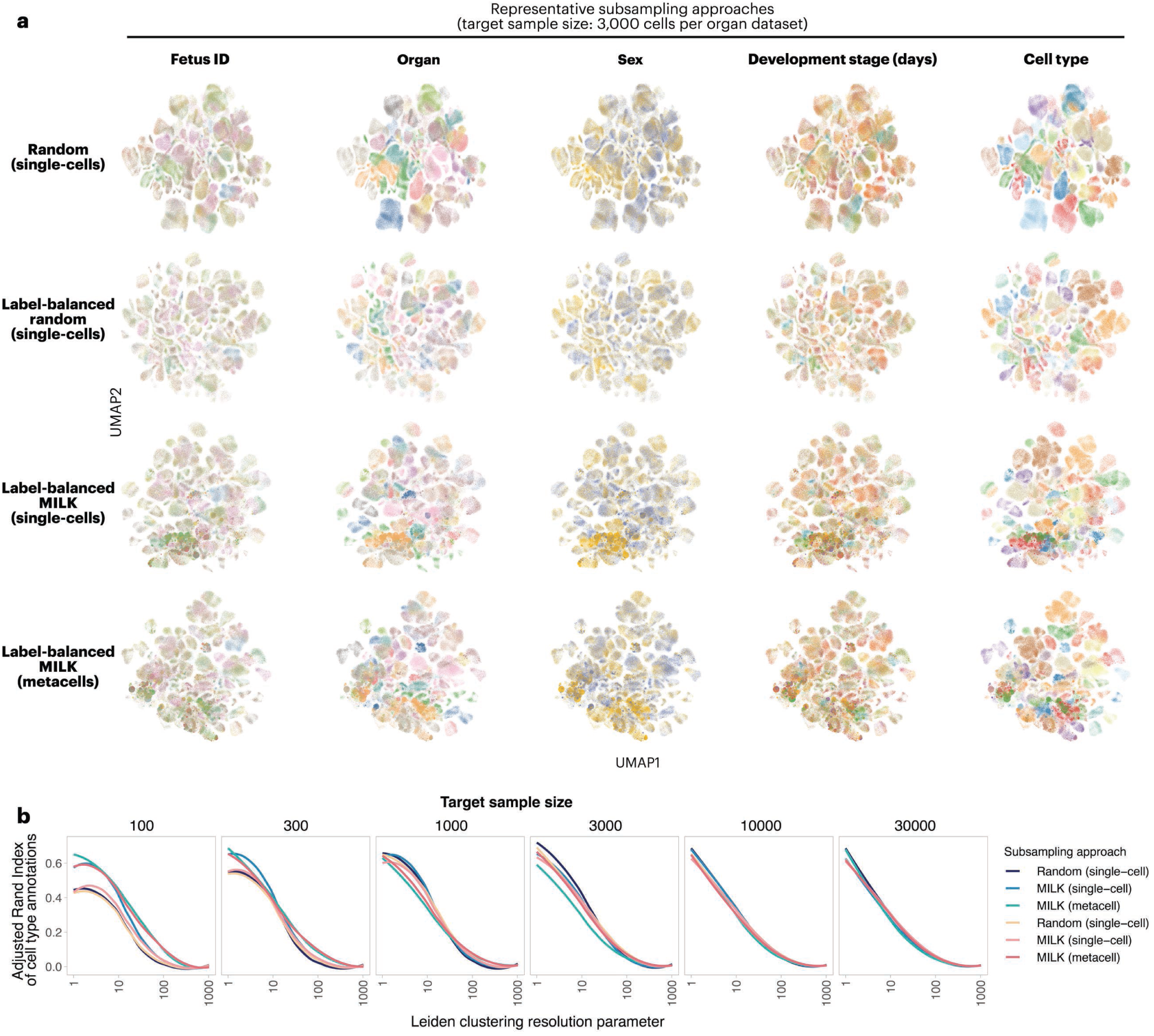
scVI integration of subsampled human fetal organ atlases. Organ-level subsamples were concatenated by target sample size and integrated with scVI using fetus ID as the batch variable, yielding 100-dimensional latent representations. **a,** Integrated atlas embeddings across subsampling strategies. UMAP embeddings of scVI-integrated human fetal atlases generated from organ subsamples with a target size of 3,000 cells per organ. Rows correspond to representative subsampling strategies, and columns show the same integrated embeddings colored by fetus ID, organ, sex, developmental stage and cell type. **b,** Resolution-dependent cell type clustering. Agreement between cell type labels and Leiden clusters was evaluated for integrated atlases generated at target sample sizes of 100, 300, 1,000, 3,000, 10,000 and 30,000 cells per organ. Adjusted Rand index (ARI) scores were calculated across Leiden clustering resolutions and plotted as a function of resolution on a log scale. Lines indicate LOESS fits for each subsampling approach.

**Supplementary Fig. 2.8.**
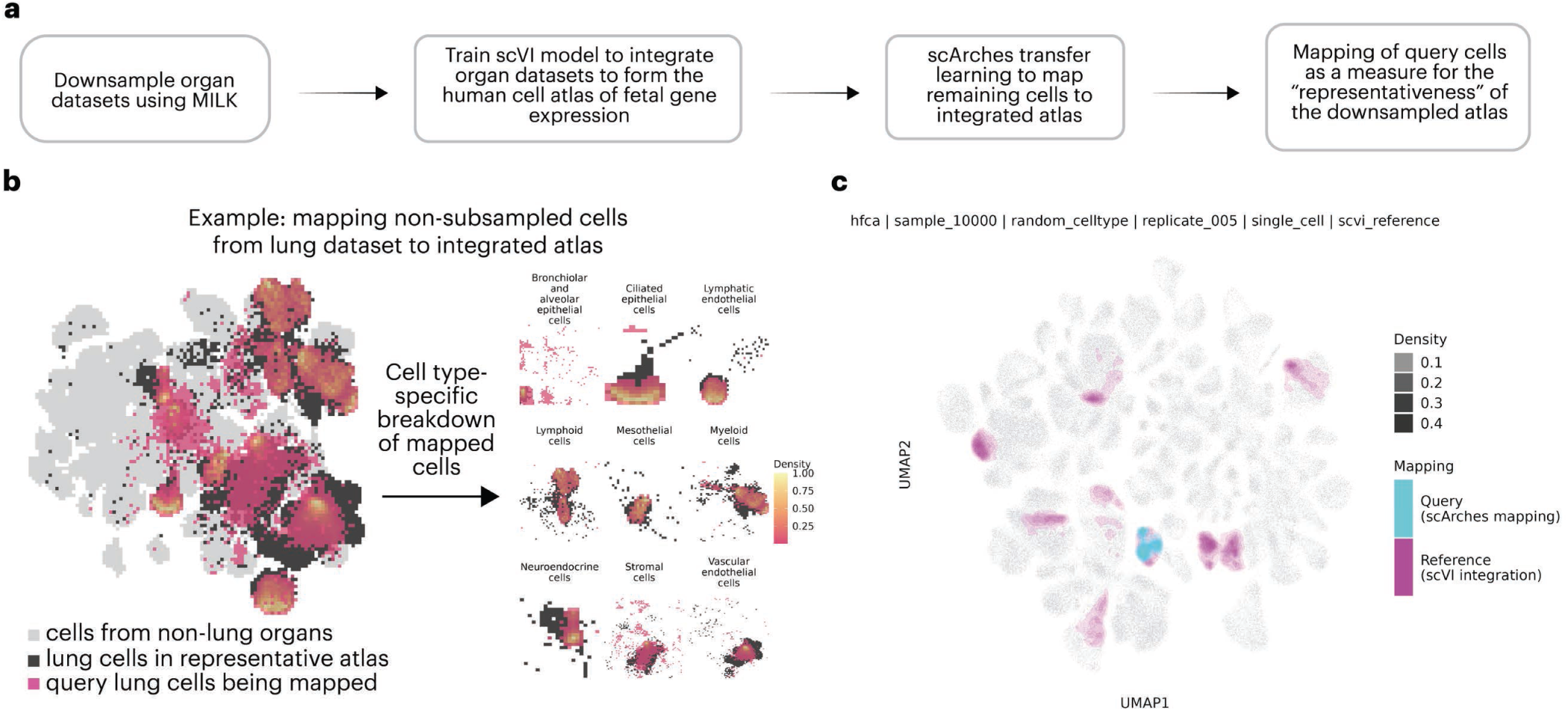
Reference-atlas representativeness by query-cell mapping. **a,** Query-mapping workflow. Organ datasets were first downsampled using representative subsampling from MILK trees, after which the organ subsamples were integrated with scVI to construct a representative human fetal cell atlas. Remaining non-subsampled cells were then mapped onto the integrated reference atlas using scArches, and representativeness was quantified as the average distance from each query cell to its nearest reference neighbors of the same cell type. **b,** Lung query-cell mapping example. Example mapping of non-subsampled lung cells onto a representative integrated atlas. Cells from non-lung organs are shown in light gray, lung cells included in the reference atlas are shown in dark gray and mapped query lung cells are shown by density. Cell type-specific breakdowns show the spatial distribution of mapped lung query cells, from which the average distance to the eight nearest reference neighbors of the same cell type was calculated. **c,** Integrated reference and query-cell locations. Example UMAP embedding showing reference cells from the scVI-integrated atlas together with query cells mapped by scArches. Only cell types represented among mapped query cells were included in the representativeness analysis.

**Supplementary Fig. 3.1.**
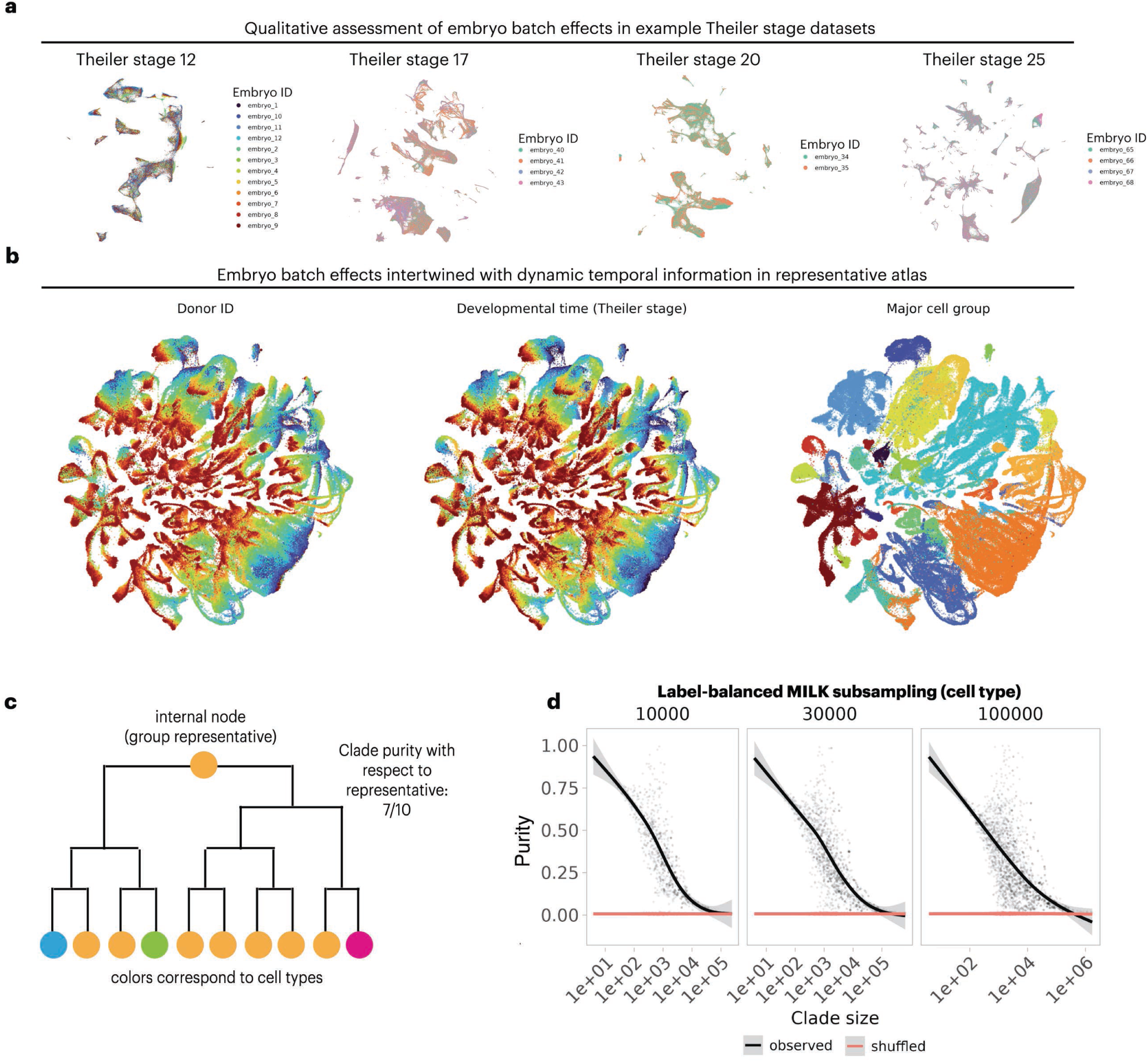
Embryo-level variation and cell type purity in mouse developmental MILK trees. **a,** Embryo identity within individual Theiler stages. UMAP embeddings of example Theiler stage-specific datasets, colored by embryo ID, for Theiler stages 12, 17, 20 and 25. **b,** Embryo identity and developmental progression in the representative atlas. UMAP embeddings of the concatenated representative mouse developmental atlas generated by cell type label-balanced subsampling of each Theiler stage to a target size of 30,000 cells. The same embedding is colored by embryo ID, developmental time represented by Theiler stage, and major cell group. **c,** Representative-based cell type purity. Schematic definition of group purity, calculated as the fraction of cells within a MILK clade that share the same cell type label as the corresponding representative cell. **d,** Cell type purity across MILK clade sizes. Cell type purity was calculated for clades in MILK trees reconstructed from cell type label-balanced representative atlases generated at target sizes of 10,000, 30,000 and 100,000 cells per Theiler stage. Purity values are plotted as a function of log-transformed clade size and compared with MILK trees in which cell type labels were shuffled.

**Supplementary Fig. 3.2.**
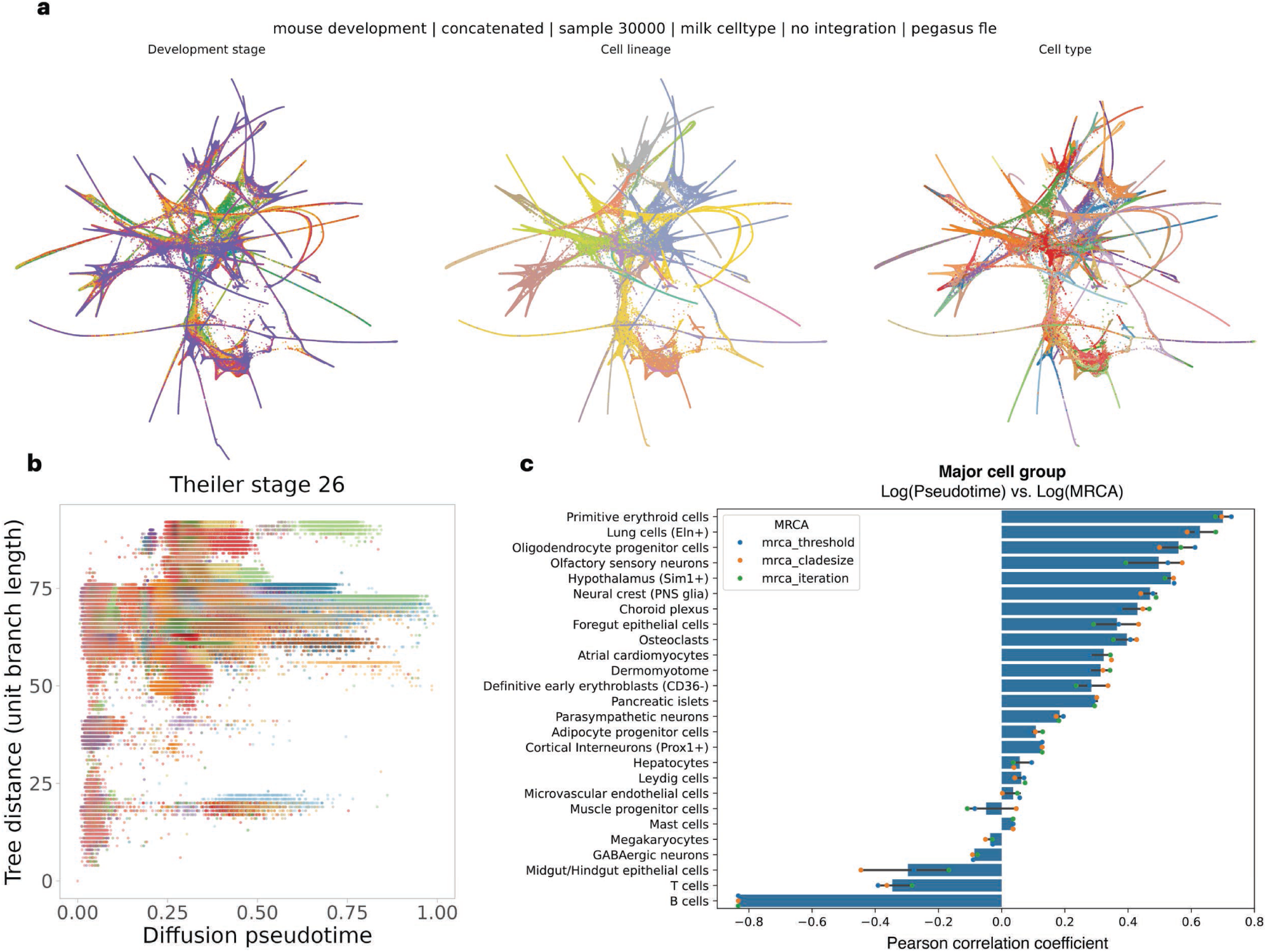
Diffusion pseudotime analysis of the mouse developmental atlas. **a,** Force-directed atlas embeddings. Force-directed layouts of the cell type label-balanced mouse developmental atlas generated from Theiler stage subsamples with a target size of 30,000 cells per stage. The same embedding is colored by developmental stage, major cell group, and cell type. **b,** Example relationship between diffusion pseudotime and MILK tree distance. Diffusion pseudotime values were computed relative to a randomly selected source cell, and MILK tree distance was calculated as the sum of unit branch lengths from each cell pair to their most recent common ancestor (MRCA). An example comparison is shown for Theiler stage 26. **c,** Major cell group-specific pseudotime concordance. Diffusion pseudotime was computed within major cell groups to avoid ambiguities arising from disconnected graph structure. For each major cell group, pseudotime values were compared with MRCA-derived MILK tree measures, including MRCA clade size, recursive MILK iteration and similarity threshold. Pearson correlation coefficients summarize the concordance between diffusion pseudotime and each MILK tree-derived measure.

**Supplementary Fig. 3.3.**
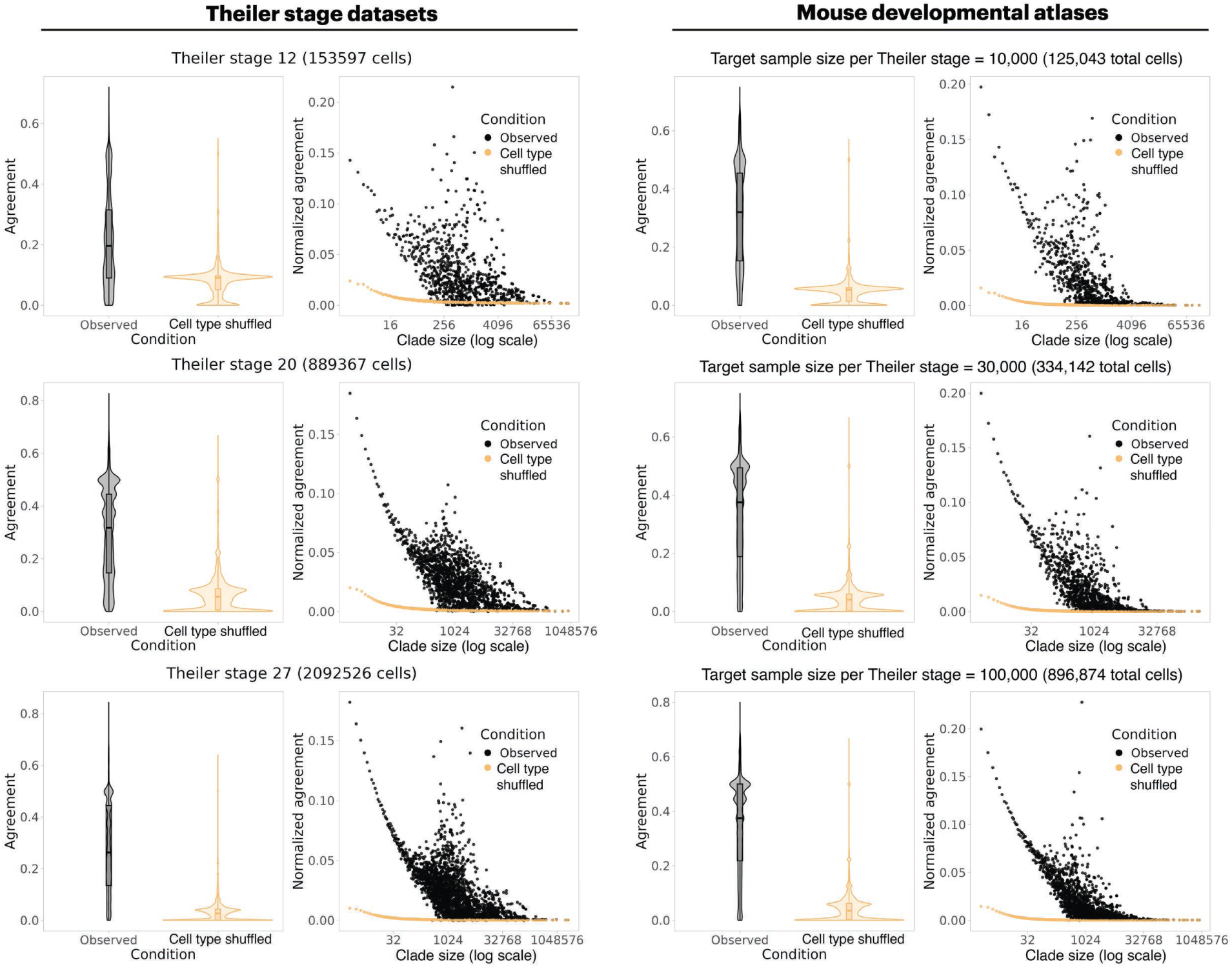
PAGA concordance of Theiler stage-specific and global developmental MILK trees. PAGA-based analysis of cell type relationships captured by MILK clades in mouse developmental datasets. MILK trees were evaluated for three representative Theiler stage-specific datasets, corresponding to Theiler stages 12, 20 and 27, and for global developmental atlases generated from cell type label-balanced subsamples with target sizes of 10,000, 30,000 and 100,000 cells per Theiler stage. For each dataset, a PAGA cell type connectivity graph was inferred using default parameters. Each MILK clade was decomposed into a cell type composition vector, with entries corresponding to the relative proportions of cell types captured by that clade. The resulting clade-level cell type vector was compared with the PAGA-derived cell type connectivity matrix using a quadratic form to obtain a PAGA agreement score. Distributions of agreement scores are shown for observed MILK trees and corresponding cell type-shuffled controls. Scatterplots show clade-level normalized agreement scores as a function of clade size on a log scale, with points representing individual clades and fitted trends summarizing clade-size-dependent agreement.

**Supplementary Fig. 3.4.**
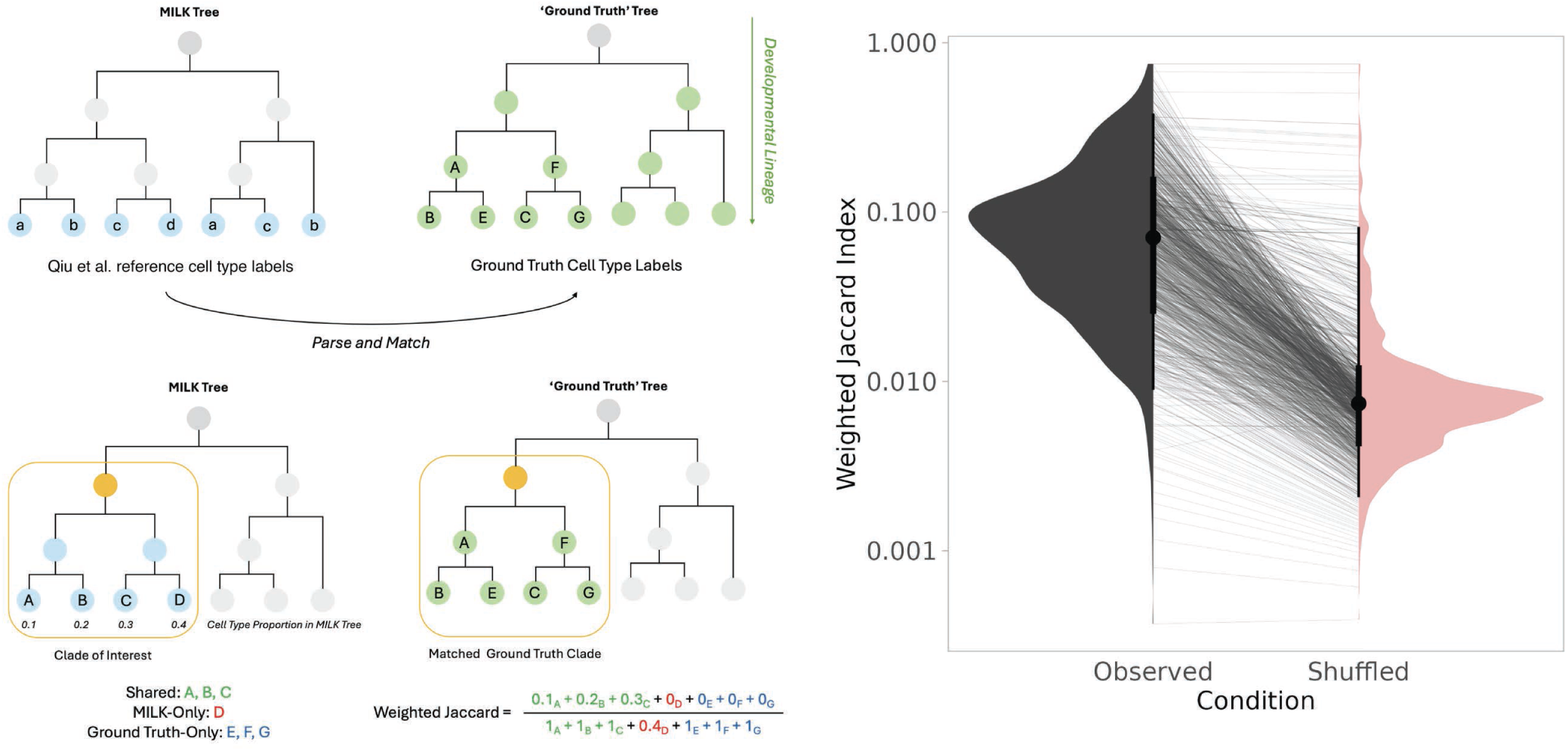
Agreement between MILK clades and curated mouse developmental cell type relationships. **a,** Asymmetric weighted Jaccard matching strategy. Cell type relationships encoded in the MILK tree were compared with the curated developmental cell type relationship graph proposed by Qiu et al. For each MILK clade, leaf cells were decomposed into a vector of relative cell type proportions, and this vector was compared against each clade in the reference cell type graph using an asymmetric weighted Jaccard index. In this comparison, MILK clade weights corresponded to the relative proportions of cell types captured by the clade, whereas cell types in the reference graph were assigned binary weights. Each MILK clade was assigned the maximum weighted Jaccard score across all candidate reference clades. Because the reference cell type graph was not strictly a directed acyclic graph, all clades across the multi-root reference structure were considered during matching. **b,** Observed versus shuffled cell type-label agreement. Distributions of maximum weighted Jaccard scores for MILK clades are shown for the observed MILK tree and a negative-control MILK tree with shuffled cell type labels.

**Supplementary Fig. 3.5.**
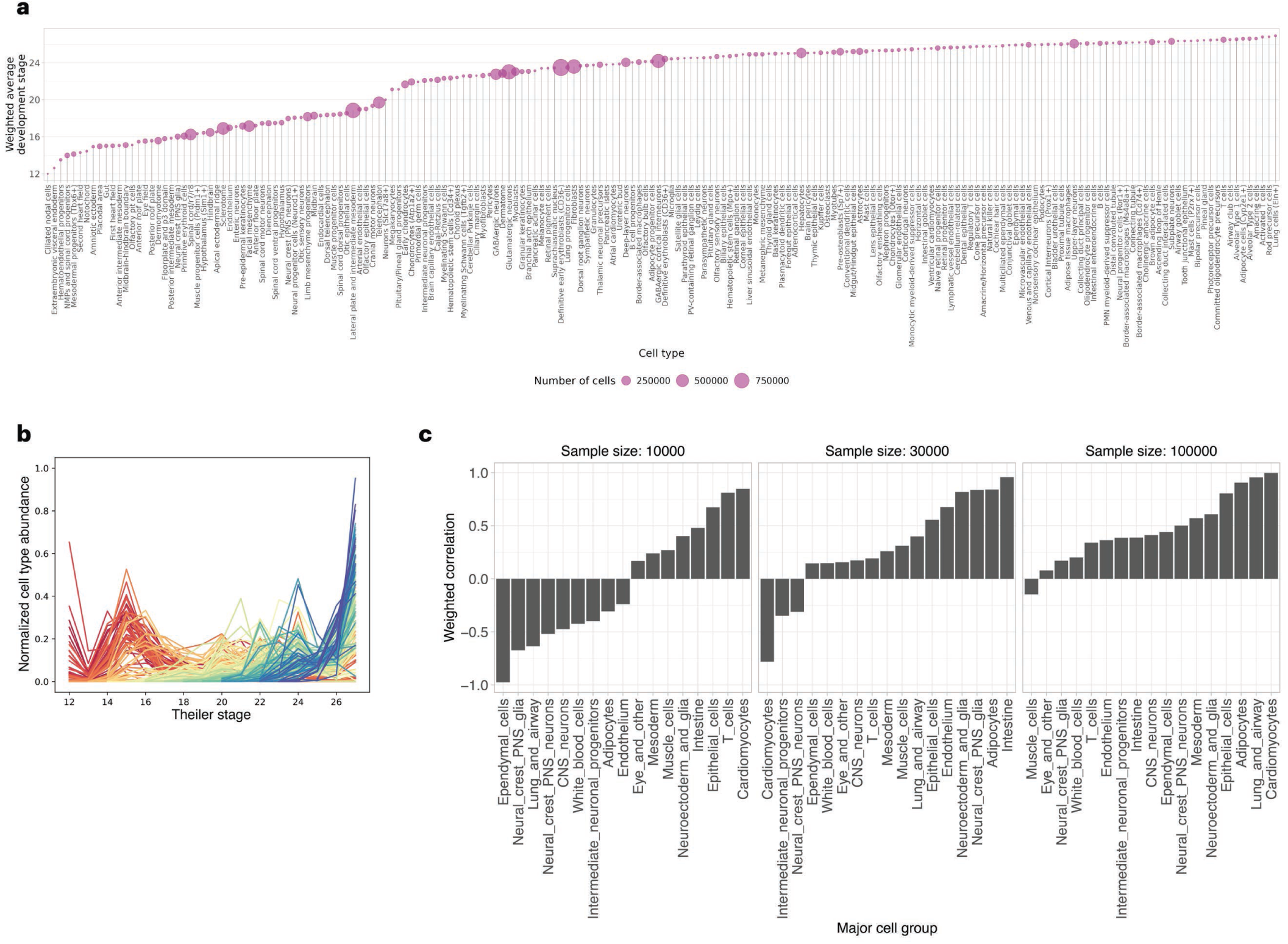
Developmental timing and branching-depth bias of cell types in mouse developmental MILK trees. **a,** Cell type distribution across developmental time. Weighted average Theiler stage for each cell type in the mouse developmental atlas, calculated using the abundance of that cell type across Theiler stages as weights. Point size denotes the total number of cells assigned to each cell type. **b,** Cell type abundance dynamics across Theiler stages. Normalized abundance profiles of cell types across Theiler stages, with counts normalized by the total number of cells captured at each stage. Each line represents one cell type, and color denotes its weighted average Theiler stage. **c,** Major cell group-specific association between developmental timing and MILK branching depth. Weighted Pearson correlations between the weighted average Theiler stage of cell types and their weighted average branching depth in the MILK tree, stratified by major cell group. Developmental timing was weighted by cell type abundance across Theiler stages, whereas MILK branching depth was weighted by cell type-specific clade purity. Correlations are shown for global developmental MILK trees generated from target sample sizes of 10,000, 30,000 and 100,000 cells per Theiler stage.

**Supplementary Fig. 4.1.**
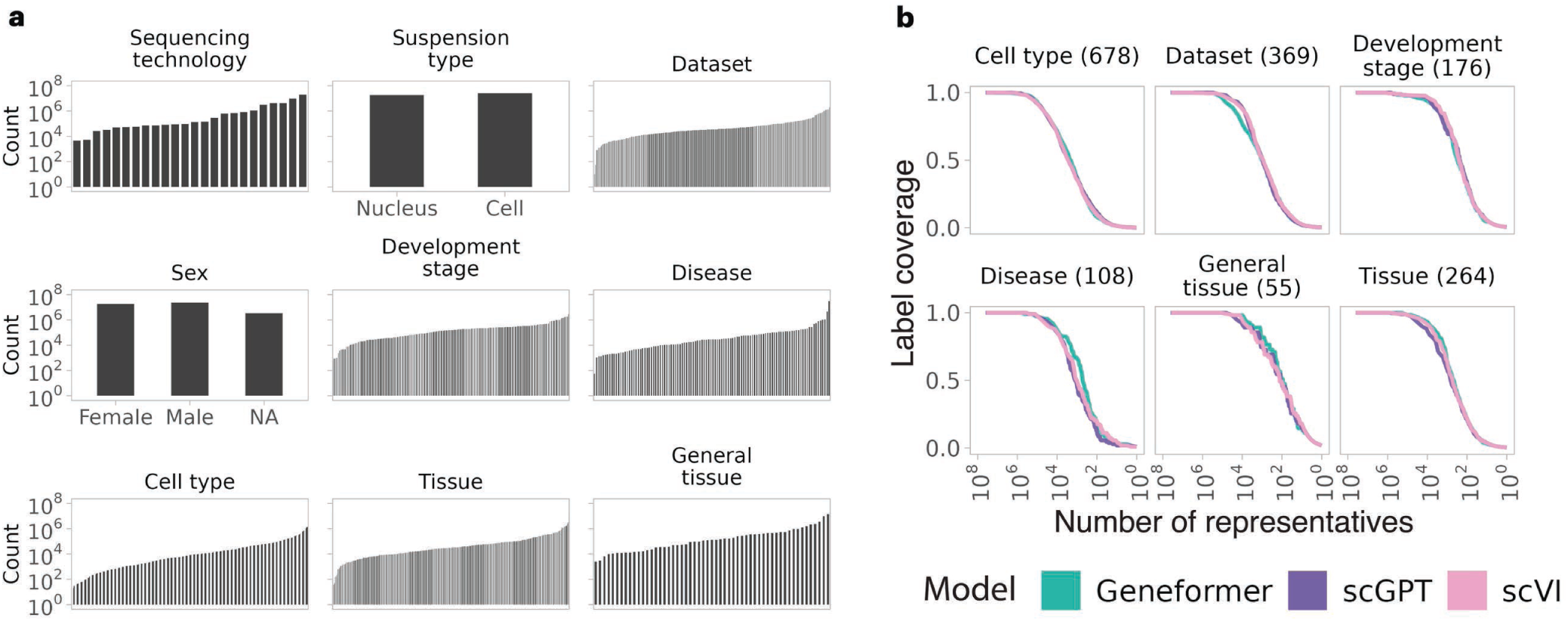
Metadata composition and label coverage in the CZ CELLxGENE Discover Census. The CZ CELLxGENE Discover Census (version: 2024-07-01) was processed by retaining primary human scRNA-seq data and filtering cells according to the metadata annotations used for downstream analyses. **a,** Metadata category abundance. Cell-count distributions across categories for nine metadata variables in the processed Census: sequencing technology, suspension type, dataset, sex, developmental stage, disease, cell type, tissue and general tissue. Bars represent metadata categories, and y-axes show cell counts on a log scale. **b,** Metadata label coverage across MILK recursion. Label coverage among group representatives across recursive MILK iterations for Geneformer-, scGPT- and scVI-derived MILK trees. Coverage was calculated as the fraction of metadata categories retained among representatives as the number of groups decreased through recursive MILK compression. The total number of categories for each metadata variable is shown in parentheses.

**Supplementary Fig. 4.2.**
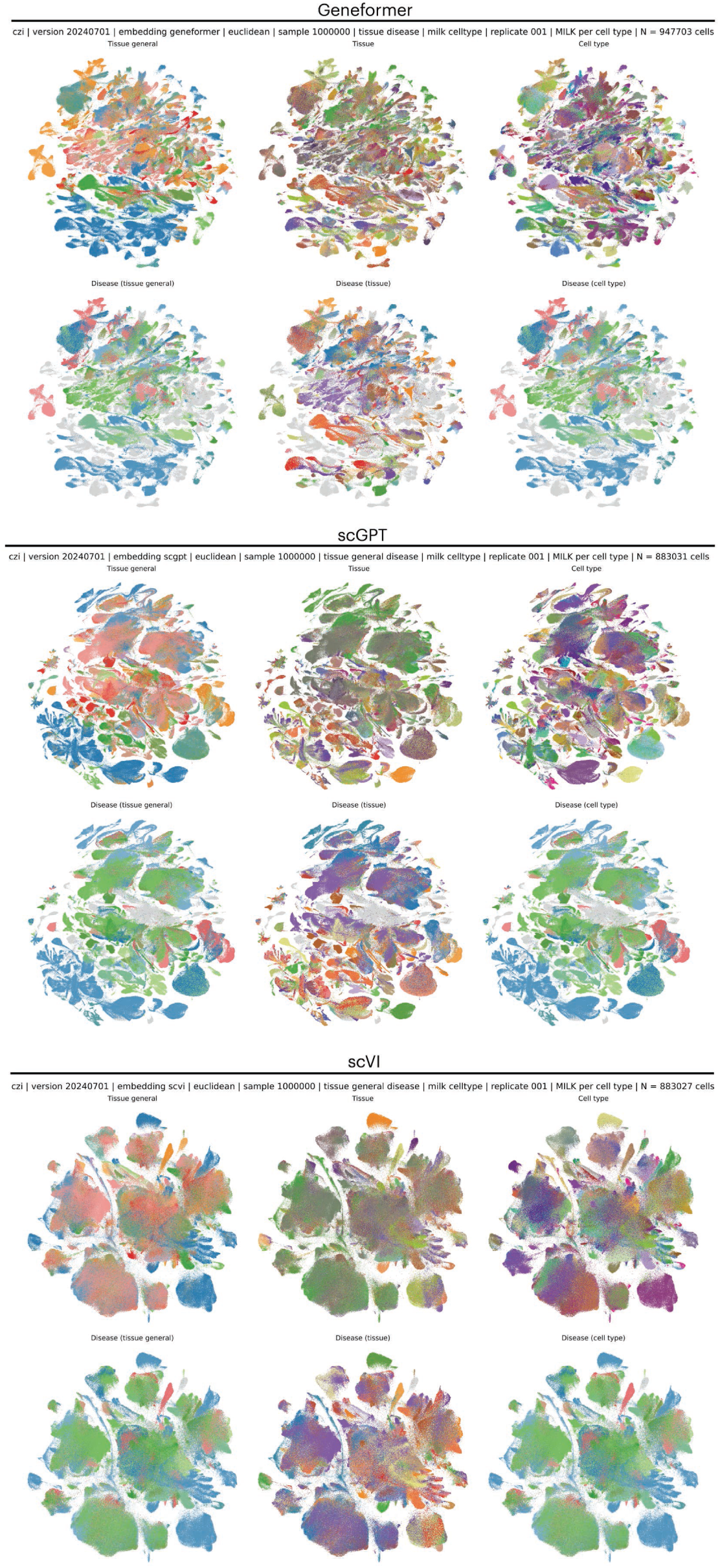
Representative CELLxGENE Census subsamples from census-trained embeddings. MILK was applied to latent embeddings of the CZ CELLxGENE Discover Census derived from Geneformer, scGPT, and scVI, and representative subsamples were extracted by label-balanced subsampling using different metadata labels. UMAP embeddings are shown for 1 million-cell representative subsamples from each model. Top row for each model: subsamples generated using general tissue, tissue, or cell type labels, with cells colored by the corresponding metadata annotation. Bottom row for each model: subsamples generated using paired disease-context labels with general tissue, tissue, or cell type labels, with cells in normal contexts shown in light gray and disease-context cells highlighted by disease annotation.

**Supplementary Fig. 4.3.**
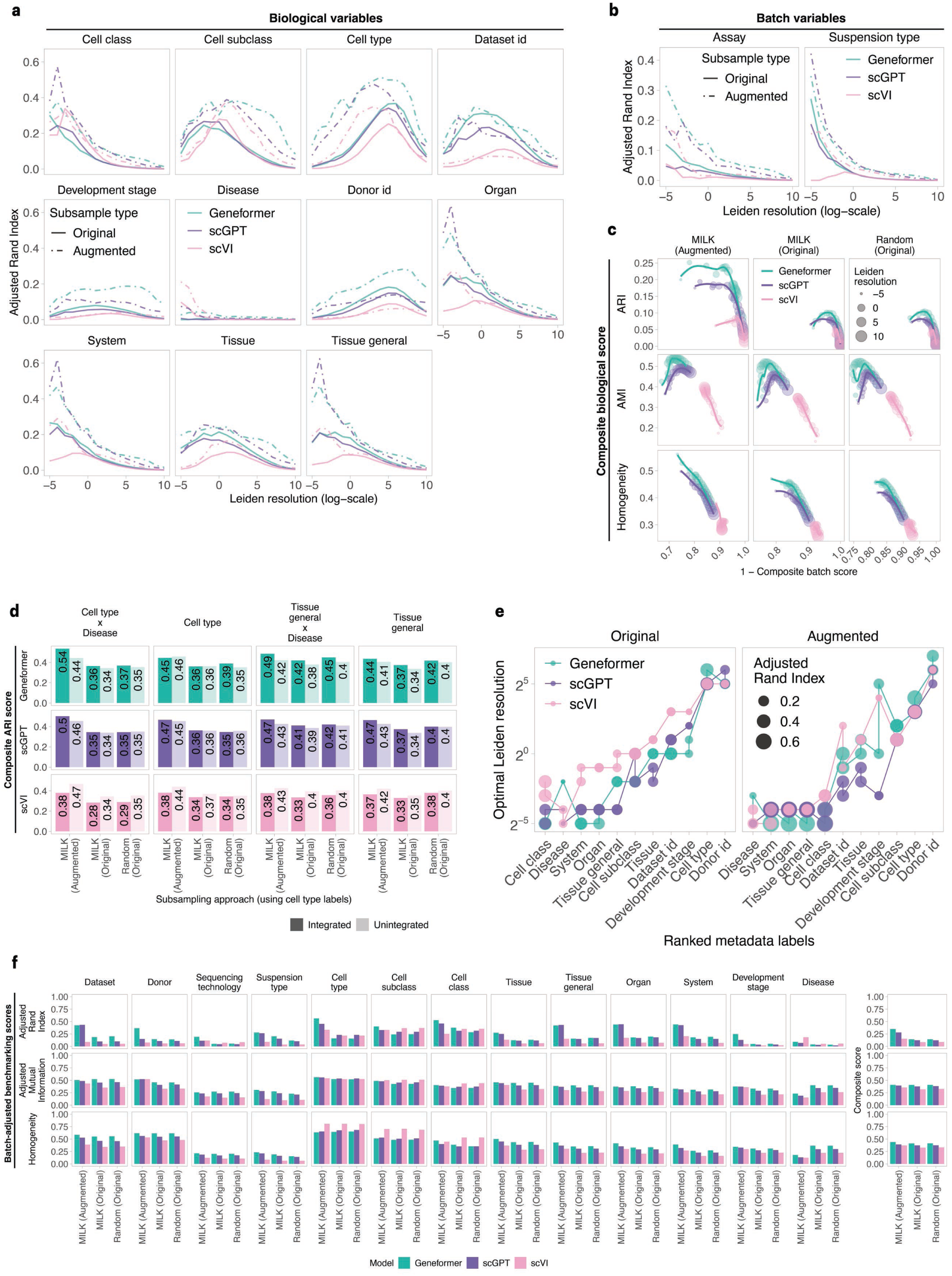
Optimization of cluster-based benchmarking across CELLxGENE Census embeddings. Leiden clustering was performed across a grid of clustering resolutions for 1 million-cell representative subsamples derived from Geneformer, scGPT and scVI embeddings of the CZ CELLxGENE Discover Census. Subsamples were generated by label-balanced subsampling using cell type, cell type–disease, tissue or tissue–disease metadata labels. MILK-derived subsamples were evaluated either as original subsamples, using only sampled representative cells, or as augmented subsamples, in which metadata counts were weighted by the clade size represented by each sampled cell. **a,** Biological label resolution sweeps. Adjusted Rand index (ARI) between Leiden clusters and biological metadata variables across Leiden clustering resolutions. Biological variables included cell class, cell subclass, cell type, dataset ID, developmental stage, disease, donor ID, organ, system, tissue and general tissue. Dataset ID and donor ID were treated as biological variables because they can capture context-specific biological structure across diverse Census datasets. Lines denote models and subsample information states. **b,** Batch label resolution sweeps. ARI between Leiden clusters and batch-associated metadata variables, including assay and suspension type, across Leiden clustering resolutions. **c,** Biological–batch score trade-off across clustering resolutions. Composite biological and batch scores were calculated at each Leiden resolution as geometric means across their respective metadata variables. The complement of the composite batch score was plotted against the composite biological score to visualize the trade-off between biological label separation and batch-associated clustering. Panels are faceted by benchmarking metric, including ARI, adjusted mutual information and homogeneity, and by subsampling approach, including MILK augmented, MILK original and random original subsamples. Lines indicate LOESS fits for each model. **d,** Composite ARI scores across subsampling labels. Global composite ARI scores were calculated as the product of the composite biological score and the complement of the composite batch score. Scores are shown for integrated model-derived embeddings and unintegrated PCA embeddings across models and metadata labels used for label-balanced subsampling. **e,** Metadata-specific optimal clustering resolutions. Leiden clustering resolutions that maximized ARI for each biological metadata variable. Metadata variables were ranked by the median optimal resolution across models and subsampling configurations. **f,** Batch-adjusted biological benchmarking scores. Optimized benchmarking scores for individual metadata variables after adjustment by the composite batch score at the corresponding Leiden clustering resolution. Scores are shown across ARI, adjusted mutual information and homogeneity metrics for Geneformer, scGPT and scVI, comparing MILK augmented, MILK original and random original subsampling approaches.

**Supplementary Fig. 4.4.**
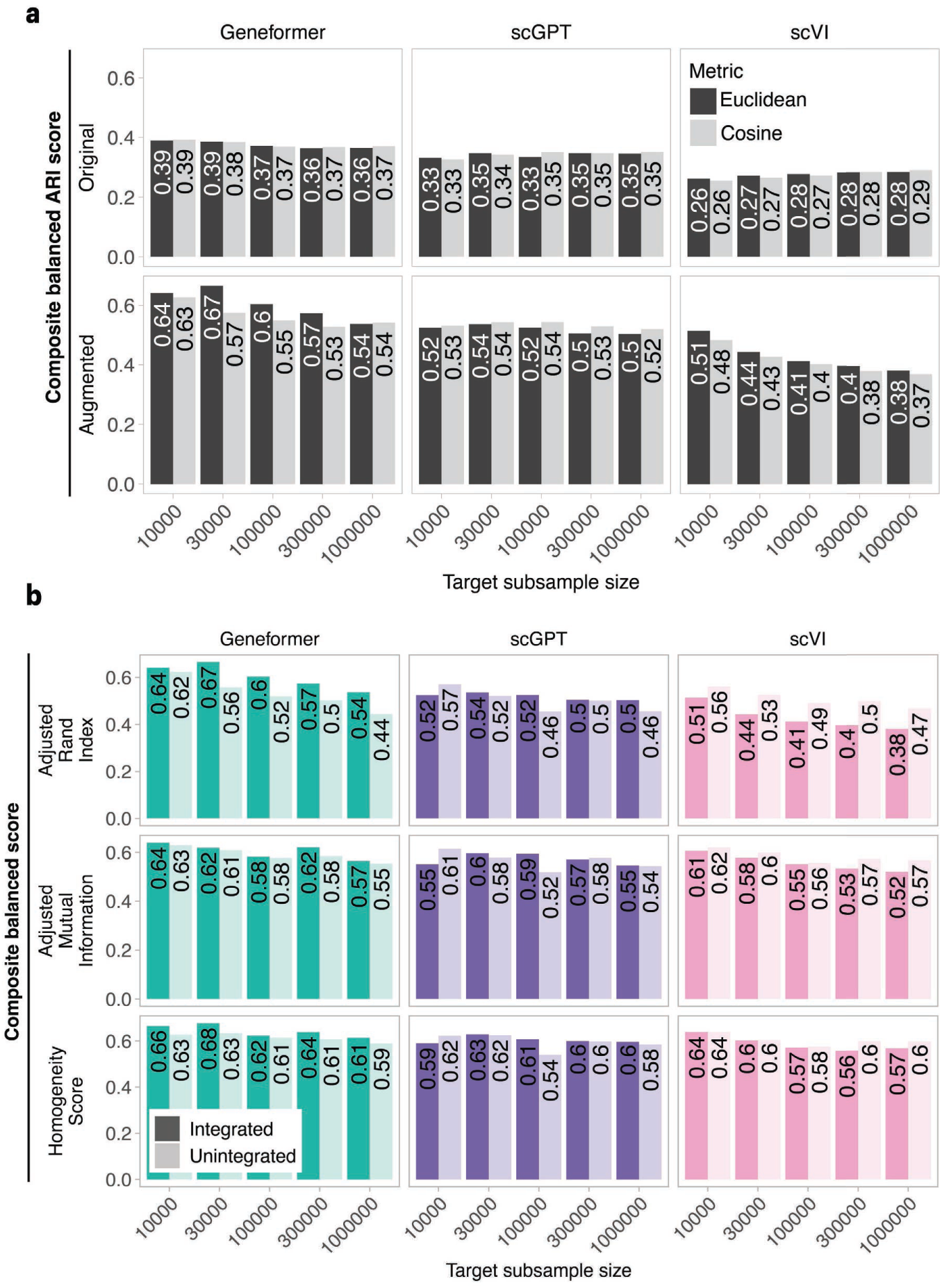
Composite balance scores across embedding and subsampling configurations. Composite balance scores were used to evaluate how well each embedding and subsampling configuration preserved biological metadata structure while minimizing batch-associated structure. For each configuration, the composite balance score was calculated as the product of the composite biological score and the complement of the composite batch score after Leiden clustering resolution optimization. Scores are shown across target subsample sizes of 10,000, 30,000, 100,000, 300,000 and 1 million cells. **a,** Distance metric and subsample augmentation effects. Composite ARI balance scores for MILK trees constructed using Euclidean or cosine distance on Geneformer-, scGPT- and scVI-derived embeddings. Scores are shown for original and augmented MILK subsamples across target subsample sizes. **b,** Multi-metric composite balance scores. Composite balance scores computed using adjusted Rand index (ARI), adjusted mutual information (AMI) and homogeneity metrics across Geneformer, scGPT and scVI embeddings. Model-derived latent embeddings (integrated) and unintegrated data are included, where the latter consists of PCA embeddings from log-normalized gene expression counts and unintegrated embedding benchmarks are shown across target subsample sizes.

**Supplementary Fig. 4.5.**
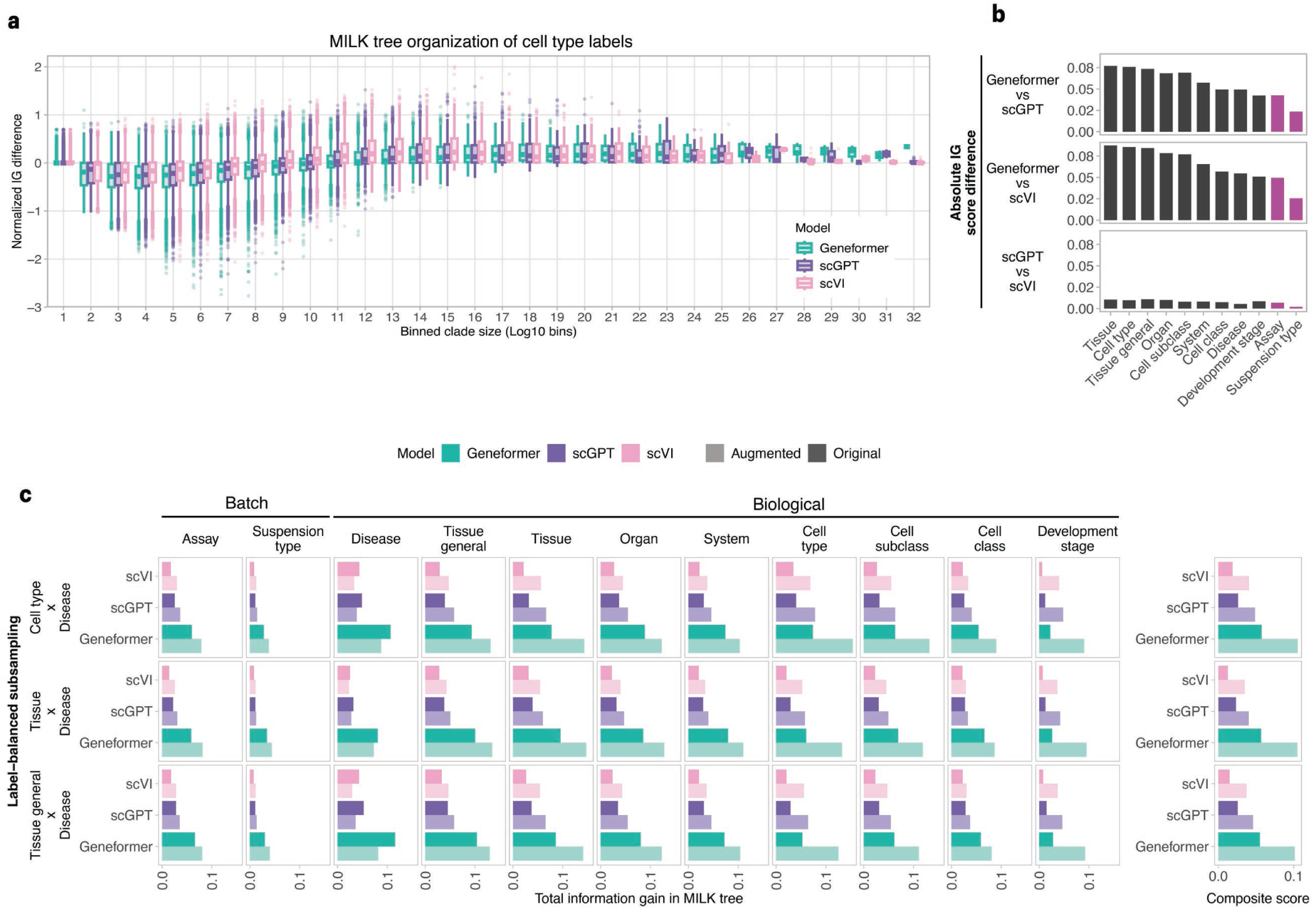
Information gain benchmarking of census-trained embeddings. Normalized information gain (IG) was calculated for clades in MILK trees derived from Geneformer, scGPT and scVI embeddings of the CZ CELLxGENE Discover Census. For each clade, raw IG was calculated as the reduction in metadata-label entropy between the parent clade and its child subclades. Raw IG values were normalized by subtracting the corresponding IG values from control MILK trees with shuffled metadata labels. **a,** Clade-size dependence of normalized IG. Normalized IG scores for cell type labels were calculated across clades in Geneformer-, scGPT- and scVI-derived MILK trees and stratified by log10-transformed clade size. Clade sizes were divided into 32 bins, and boxplots show the distribution of normalized IG scores for each model within each bin. **b,** Pairwise differences in global IG scores. Global IG scores were computed by aggregating normalized clade-level IG values using clade size as weights. Pairwise absolute differences in global IG scores were calculated between models for each metadata variable, with biological variables shown in black and batch-associated variables shown in magenta. Bars are ordered by the mean pairwise difference across metadata variables. **c,** Global IG scores across metadata labels. Weighted average normalized IG scores were computed across MILK trees derived from different embedding models, subsample information states and metadata labels used for label-balanced subsampling. Scores are shown for biological and batch-associated metadata variables across Geneformer, scGPT and scVI, comparing original and augmented MILK subsamples. Composite scores summarize biological variables by geometric mean.

**Supplementary Fig. 5.1.**
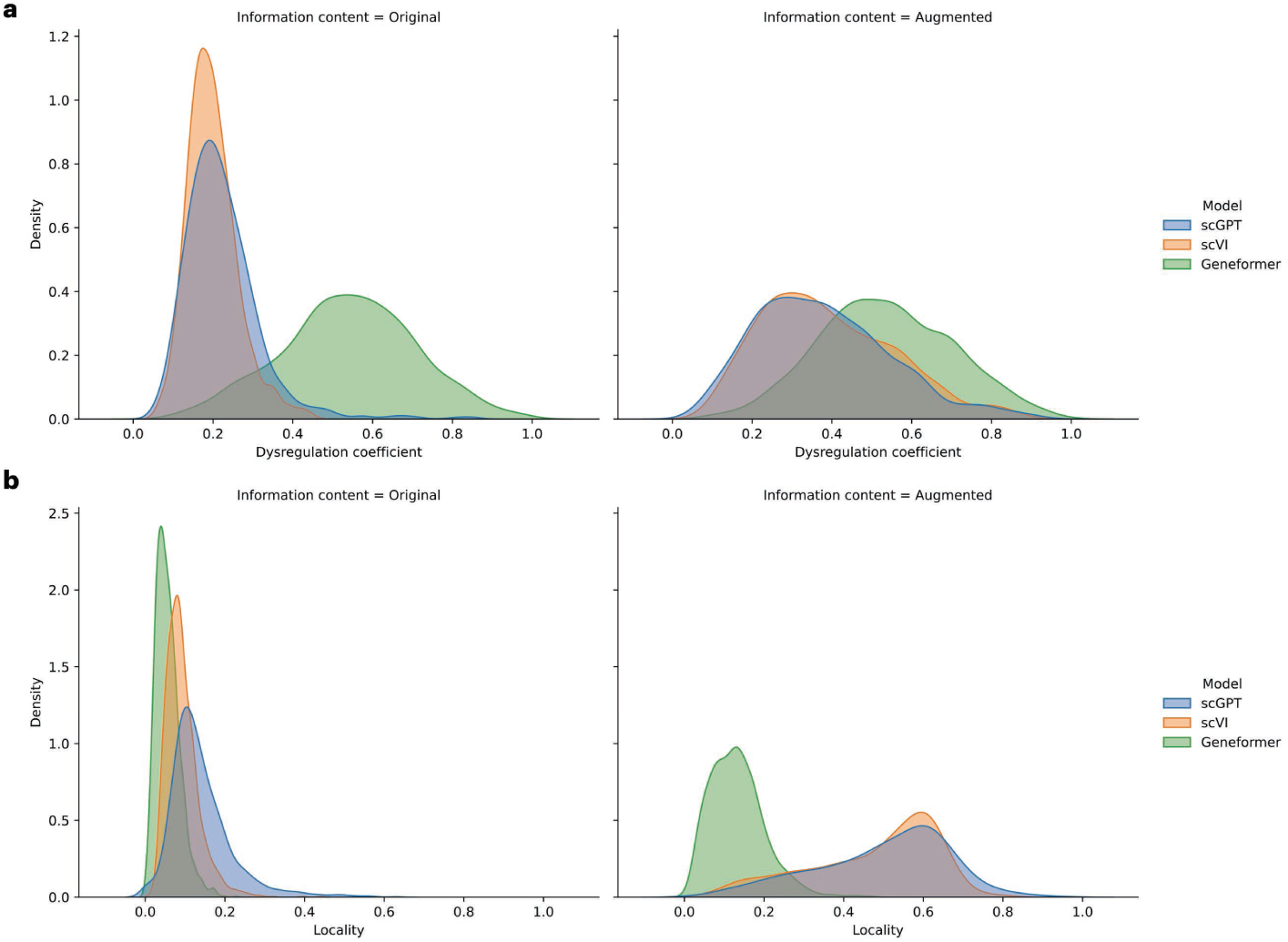
Distributions of dysregulation and locality across cell type–disease pairs. **a,** Dysregulation coefficient distributions. Density distributions of dysregulation coefficients for cell type–disease pairs across Geneformer-, scGPT- and scVI-derived MILK trees, shown for original and augmented subsample information states. For each cell type–disease pair, the dysregulation coefficient was calculated relative to the corresponding normal-context cell type as the weighted average absolute difference in clade-level relative proportions between disease-context and normal-context cells across the MILK tree. Clade size was used as the weight. **b,** Locality distributions. Density distributions of locality scores for cell type–disease pairs across Geneformer-, scGPT- and scVI-derived MILK trees, shown for original and augmented subsample information states. Locality was calculated as the weighted average clade size required to capture each cell type–disease population, using the relative proportion of the group captured by each clade as the weight. Smaller locality values indicate that a cell type–disease population is concentrated within more restricted regions of the MILK hierarchy, whereas larger values indicate broader dispersion across the tree.

**Supplementary Fig. 5.2.**
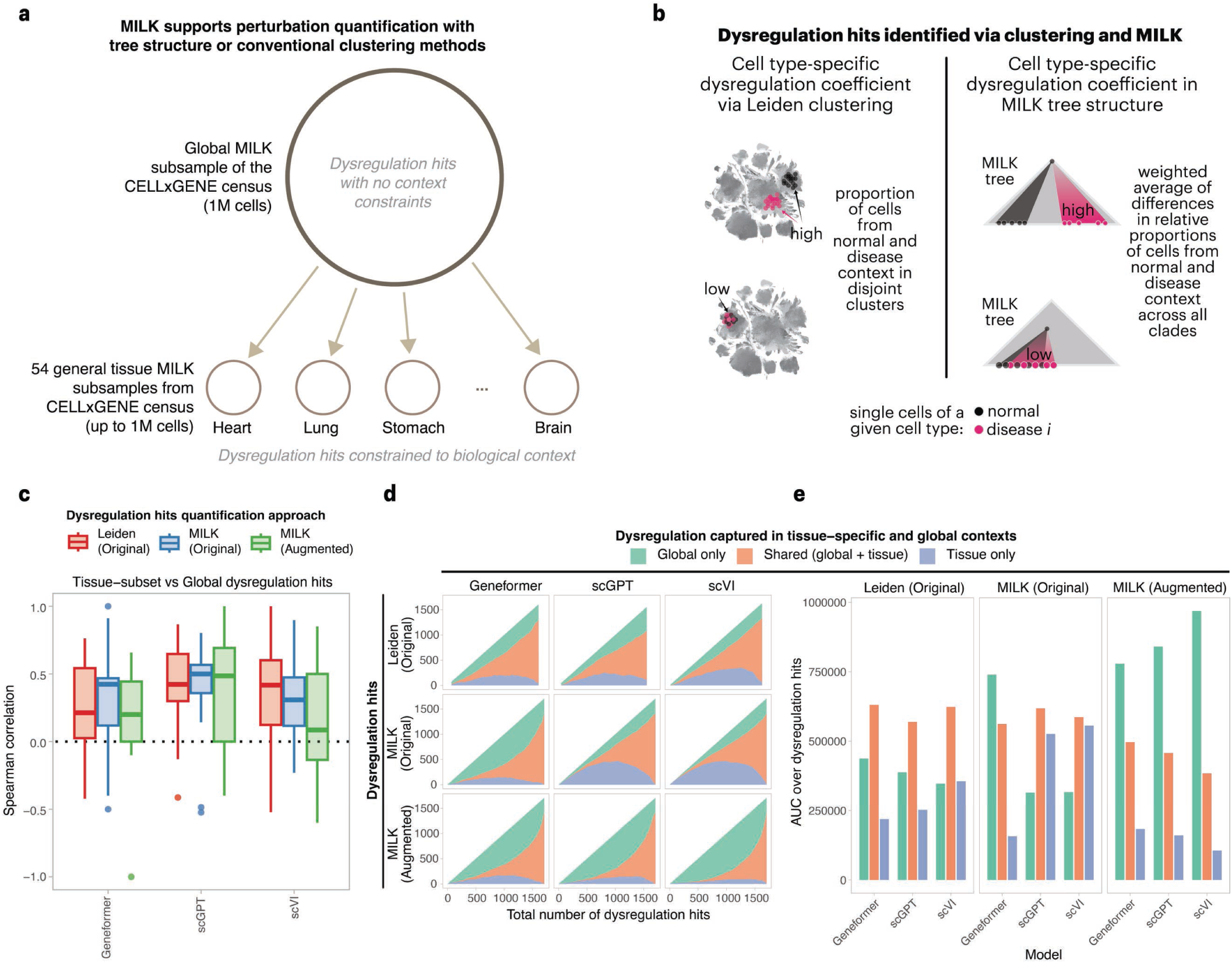
Global and tissue-specific quantification of disease-mediated cell-state dysregulation. **a,** Global and tissue-specific dysregulation comparisons. Schematic overview of dysregulation scoring in the global CZ CELLxGENE Discover Census and in tissue-specific subsets. Dysregulation coefficients were calculated for cell type–disease pairs in a global representative subsample of the Census and independently in 54 general tissue-specific subsamples. Tissue-specific analyses provided local context-restricted references for evaluating whether dysregulation patterns identified in the global hierarchy were also observed within individual tissue contexts. **b,** Leiden- and MILK-based dysregulation scoring strategies. In the clustering-based approach, dysregulation coefficients were calculated from the separation of normal-context and disease-context cells across Leiden clusters generated at multiple resolutions. In the MILK-based approach, dysregulation coefficients were calculated as the weighted average absolute difference in clade-level relative proportions between normal-context and disease-context cells across the MILK hierarchy, using clade size as the weight. **c,** Global–tissue concordance of dysregulation coefficients. For each tissue, dysregulation coefficients from the global analysis were compared with tissue-specific dysregulation coefficients across shared cell type–disease pairs. Spearman correlations were calculated for each of the 54 tissue contexts and summarized across Geneformer-, scGPT- and scVI-derived embeddings for Leiden original, MILK original and MILK augmented scoring strategies. **d,** Dysregulation hit categories across threshold sweeps. Dysregulation hits were defined by applying thresholds to continuous dysregulation coefficients. Across threshold sweeps, hits were categorized as global only, shared between global and tissue-specific analyses, or tissue only. Stacked curves show the numbers of hits in each category as a function of the total number of dysregulation hits for each model and scoring strategy. **e,** AUC summary of dysregulation hit categories. Area under the curve summaries of the hit-category distributions shown in d, comparing global-only, shared and tissue-only dysregulation hits across Leiden original, MILK original and MILK augmented scoring strategies.

**Supplementary Fig. 5.3.**
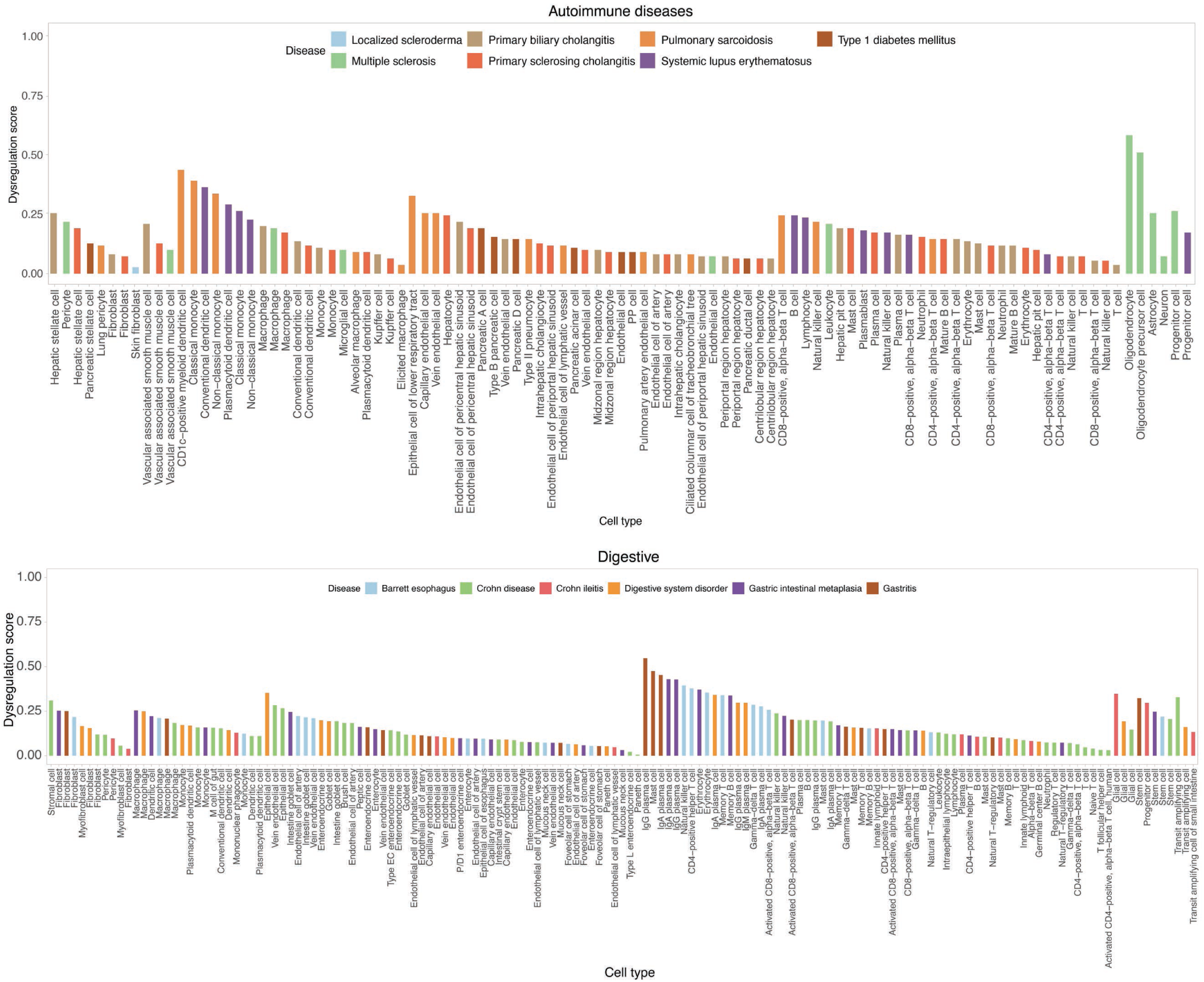
Cell type–disease dysregulation coefficients for autoimmune and digestive disease classes. Dysregulation coefficients were calculated for cell type–disease pairs using the 1 million-cell Geneformer-based MILK tree. Values are shown for representative disease contexts within the autoimmune disease class (top) and digestive disease class (bottom). Bars represent cell type–disease pairs, with bar height indicating the dysregulation coefficient and color denoting disease context.

**Supplementary Fig. 5.4.**
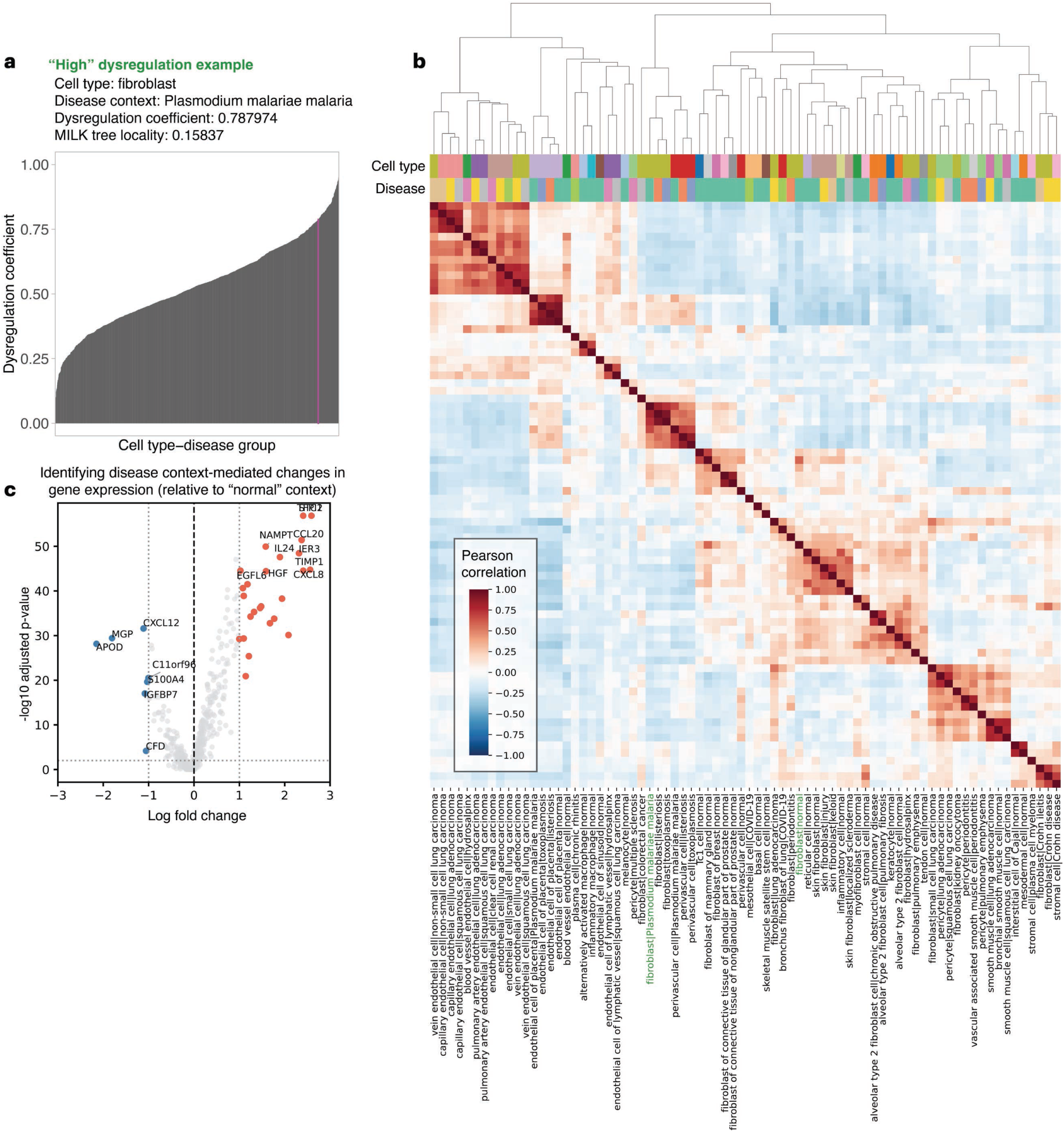
High-dysregulation fibroblast–malaria example in the MILK hierarchy. **a,** Fibroblast–malaria dysregulation coefficient. Dysregulation coefficients for all cell type–disease pairs in the Geneformer-based MILK tree, with the fibroblast–*Plasmodium malariae* malaria pair highlighted as a high-dysregulation example. The highlighted pair had a dysregulation coefficient of 0.787974 and a MILK tree locality score of 0.15837. **b,** Clade-level transcriptomic separation of normal and malaria-context fibroblasts. The smallest MILK clades capturing a majority of fibroblasts in normal context and in *Plasmodium malariae* malaria context were identified. The normal-context fibroblast clade contained 21,516 cells, whereas the malaria-context fibroblast clade contained 822 cells. For all cell type–disease groups represented by at least 100 cells within these clades, aggregate gene-expression profiles were calculated by averaging expression across cells in each group. Pairwise Pearson correlations between aggregate profiles were then hierarchically clustered to identify transcriptionally related cell type–disease groups. **c,** Differential gene expression in malaria-context fibroblasts. Volcano plot comparing fibroblast gene expression in *Plasmodium malariae* malaria relative to normal context. Upregulated genes are shown in red and downregulated genes in blue, defined by absolute log fold change ≥ 1 and adjusted *P* value < 0.01. The top upregulated and downregulated genes are labeled.

**Supplementary Fig. 5.5.**
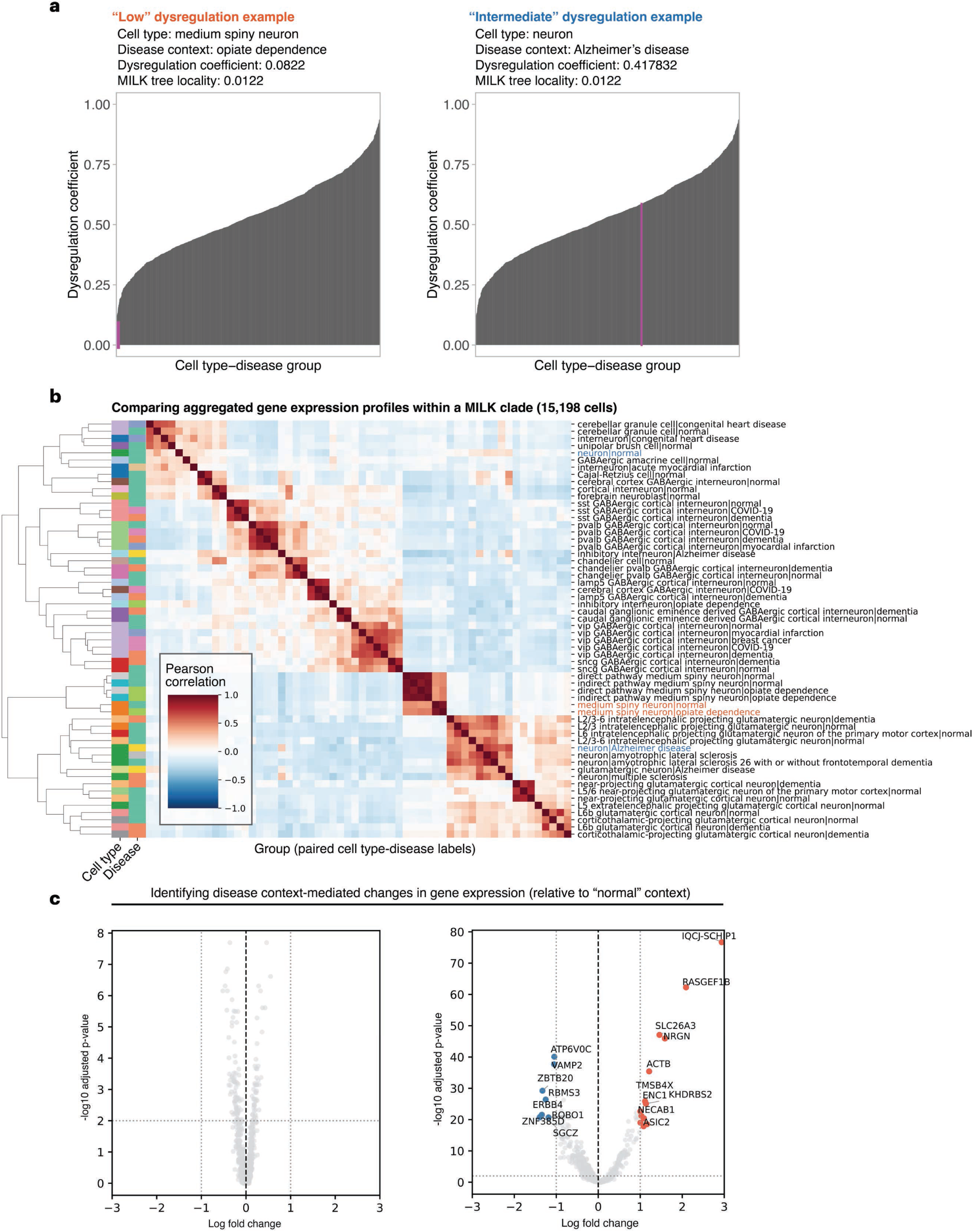
Low- and intermediate-dysregulation neuronal disease examples in the MILK hierarchy. **a,** Neuronal cell type–disease dysregulation coefficients. Dysregulation coefficients for all cell type–disease pairs in the Geneformer-based MILK tree, with medium spiny neuron–opiate dependence highlighted as a low-dysregulation example and neuron–Alzheimer’s disease highlighted as an intermediate-dysregulation example. The medium spiny neuron–opiate dependence pair had a dysregulation coefficient of 0.0822 and a MILK tree locality score of 0.0122, whereas the neuron– Alzheimer’s disease pair had a dysregulation coefficient of 0.417832 and a MILK tree locality score of 0.0122. **b,** Clade-level transcriptomic organization of neuronal disease contexts. The smallest MILK clade capturing a majority of both neuronal cell type–disease populations and their corresponding normal-context populations was identified. For all cell type–disease groups represented by at least 100 cells within this clade, aggregate gene-expression profiles were calculated by averaging expression across cells in each group. Pairwise Pearson correlations between aggregate profiles were then hierarchically clustered to identify transcriptionally related cell type–disease groups. **c,** Differential gene expression in low- and intermediate-dysregulation examples. Volcano plots comparing disease-context gene expression with corresponding normal-context expression for medium spiny neuron–opiate dependence and neuron–Alzheimer’s disease. Upregulated genes are shown in red and downregulated genes in blue, defined by absolute log fold change ≥ 1 and adjusted *P* value < 0.01. The top upregulated and downregulated genes are labeled.

**Supplementary Fig. 5.6.**
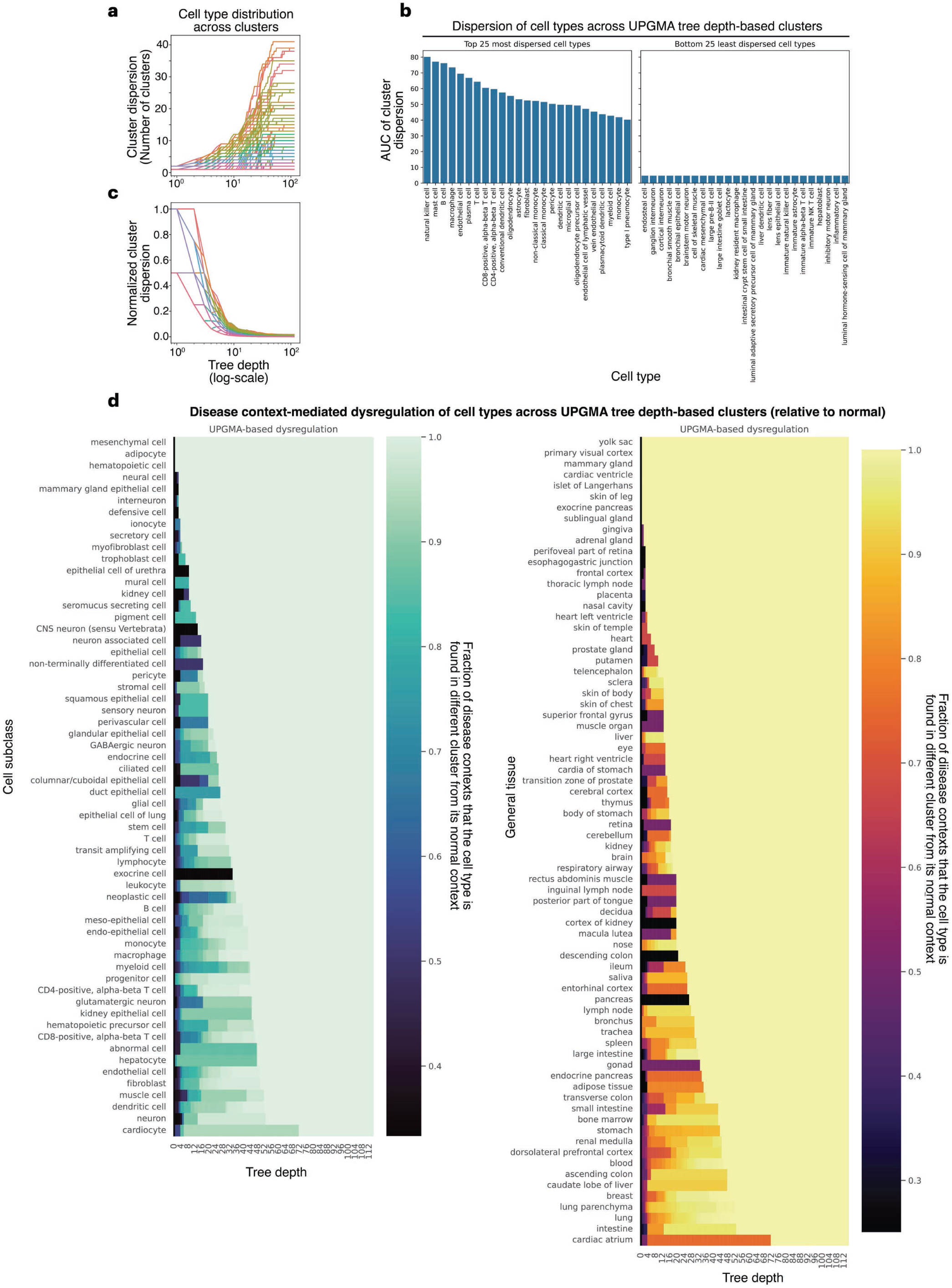
Depth-dependent dispersion of cell type–disease groups across UPGMA clusters. Pairwise associations between cell type–disease groups were calculated from the MILK hierarchy and clustered using UPGMA. Clusters were then defined at different depths of the UPGMA tree to evaluate how cell types and disease contexts distribute across hierarchical resolutions. **a,** Cell type distribution across UPGMA cluster depths. For each cell type, the number of UPGMA depth-based clusters containing that cell type was calculated across disease contexts and plotted as a function of tree depth. **b,** Cell type dispersion rankings. Cell type dispersion across UPGMA cluster depths was summarized by the area under the curve of the normalized dispersion profile. The 25 most dispersed and 25 least dispersed cell types are shown. **c,** Disease-context-normalized cell type dispersion. Cluster counts from **a** were normalized by the number of disease contexts in which each cell type was represented, providing a context-adjusted measure of cell type dispersion across UPGMA cluster depths. **d,** Disease-context-mediated separation from normal cell states. For each cell type and UPGMA tree depth, disease-context groups were compared with the corresponding normal-context group. Dysregulation was quantified as the fraction of disease contexts in which the cell type appeared in a different UPGMA cluster from its normal-context counterpart. Values were aggregated by cell subclass (left) or general tissue (right) by averaging across cell types.

**Supplementary Fig. 5.7.**
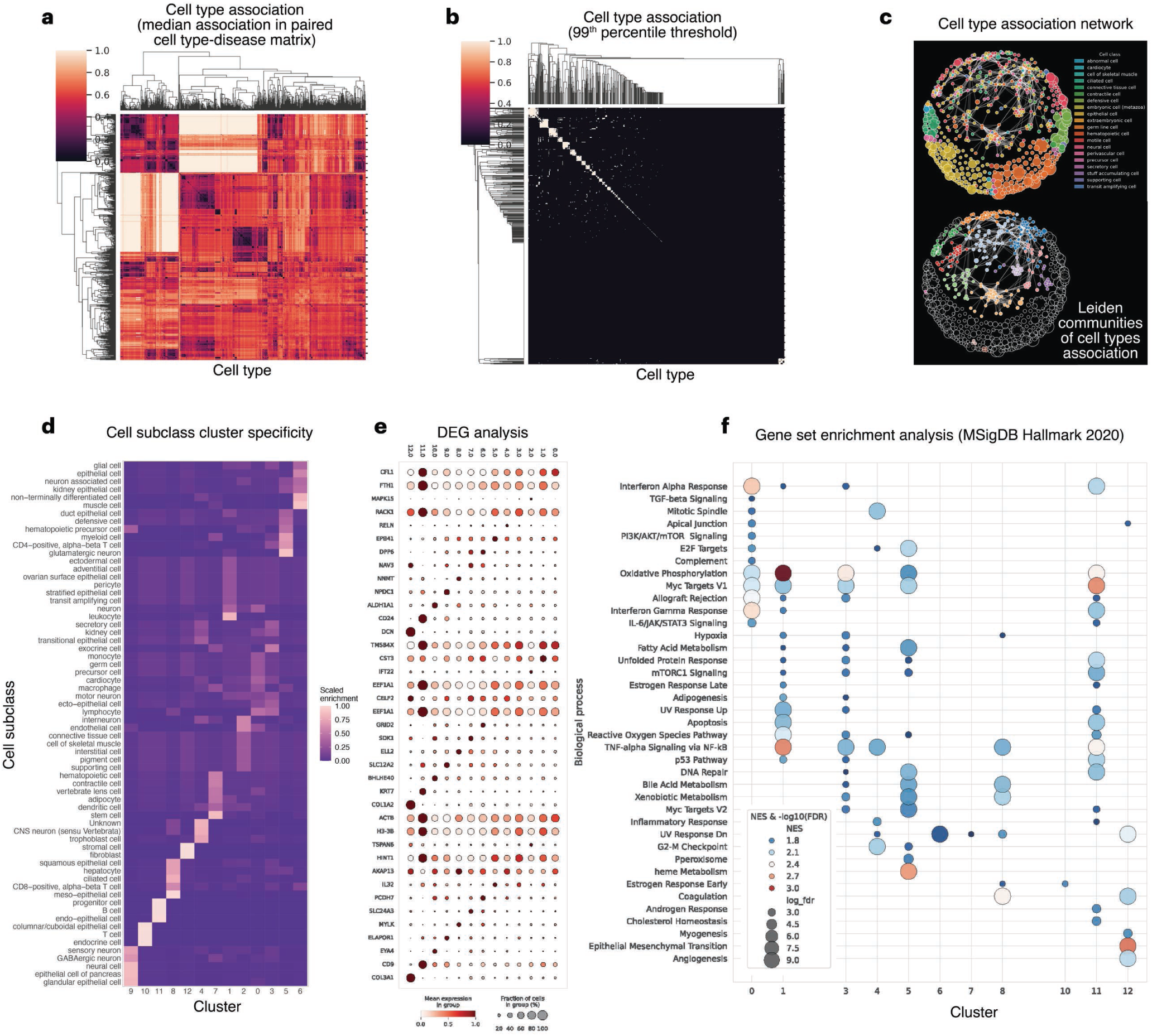
Gene set enrichment analysis of the cell type association network. **a,** Cell type association matrix. The pairwise association matrix of cell type–disease groups was aggregated into a context-agnostic cell type–cell type association matrix by taking the median normalized clade-size value for each pair of cell types across disease contexts. Smaller values indicate stronger co-localization within the global MILK hierarchy. **b,** Thresholded cell type association matrix. Strong cell type associations were retained by applying a stringent percentile threshold to the cell type–cell type association matrix. The resulting binarized matrix was used to define edges in the cell type association network. **c,** Cell type association network. Cell type networks were visualized using an attractive and repulsive forces layout. Nodes denote cell types and edges denote strong associations retained after thresholding. Top, nodes colored by cell class. Bottom, Leiden community detection applied to the thresholded network, with communities containing at least two cell types highlighted by color. **d,** Cell subclass composition of network communities. Cell subclass composition of the Leiden communities identified in c. The matrix was scaled using the Sinkhorn–Knopp algorithm to visualize enrichment patterns across communities. **e,** Differentially expressed genes across cell type communities. Differentially expressed genes were identified for each Leiden community in the cell type association network. Dot size denotes the fraction of cells expressing each gene, and color denotes mean expression. **f,** Gene set enrichment across cell type communities. Gene set enrichment analysis was performed against the MSigDB Hallmark 2020 gene sets collection using ranked differentially expressed genes from each Leiden community. Enriched terms are shown across communities, with dot size indicating the number of associated genes and color indicating normalized enrichment score.

**Supplementary Fig. 5.8.**
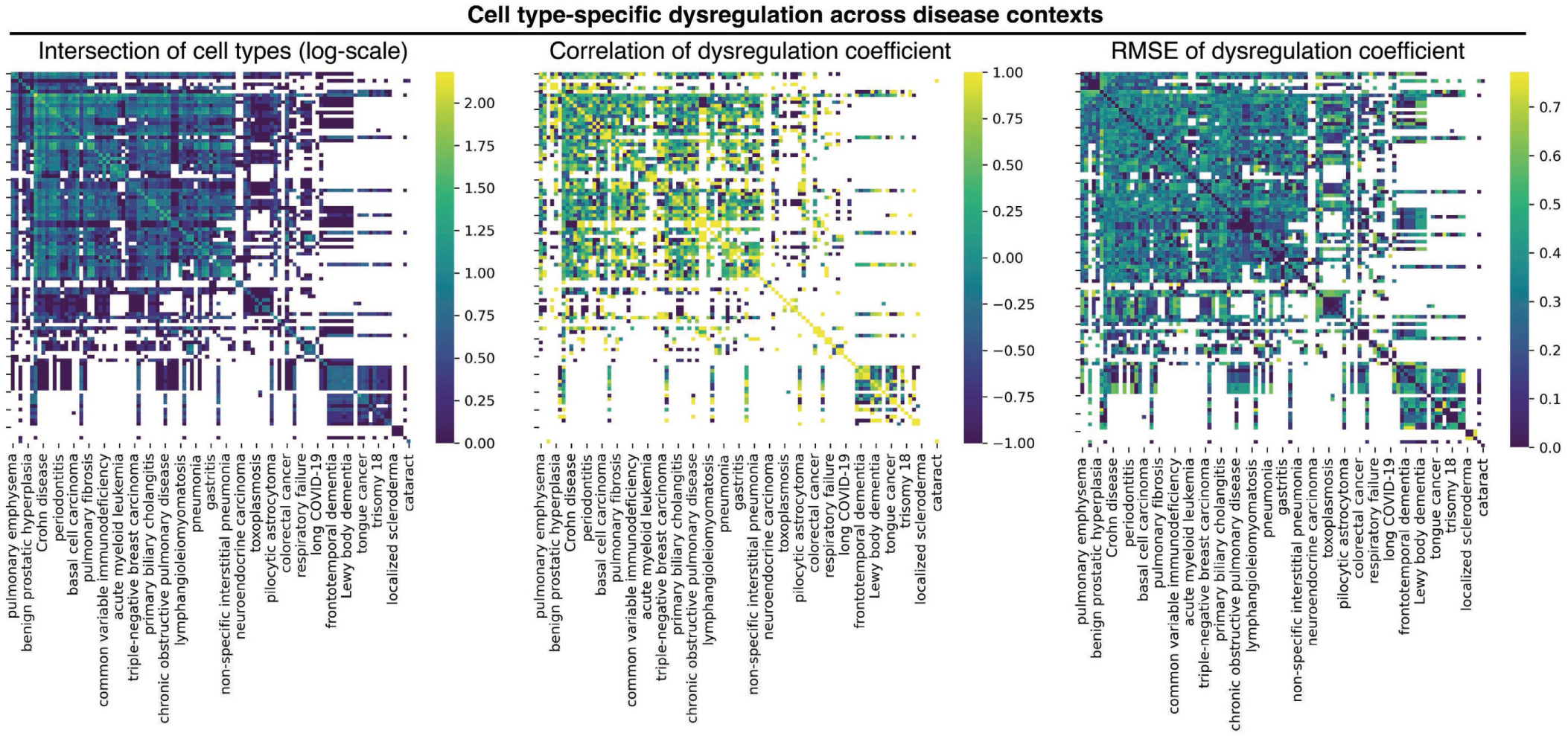
Disease–disease matrices underlying the disease association network. Disease–disease relationships were evaluated by comparing cell type-specific dysregulation coefficients across shared cell types. **a,** Shared cell type coverage. Heatmap showing the log-transformed number of cell types shared between each pair of disease contexts. White entries indicate disease pairs with no shared cell types or insufficient shared cell type coverage for downstream comparison. **b,** Correlation of dysregulation coefficients. Pearson correlations of dysregulation coefficients between disease pairs, calculated across shared cell types for disease pairs with at least two shared cell types. This correlation matrix was used to define edges in the disease association network. **c,** Dysregulation coefficient RMSE. Root mean square error (RMSE) between dysregulation coefficient vectors for each disease pair across shared cell types, providing a complementary measure of absolute differences in dysregulation magnitude.

**Supplementary Fig. 5.9.**
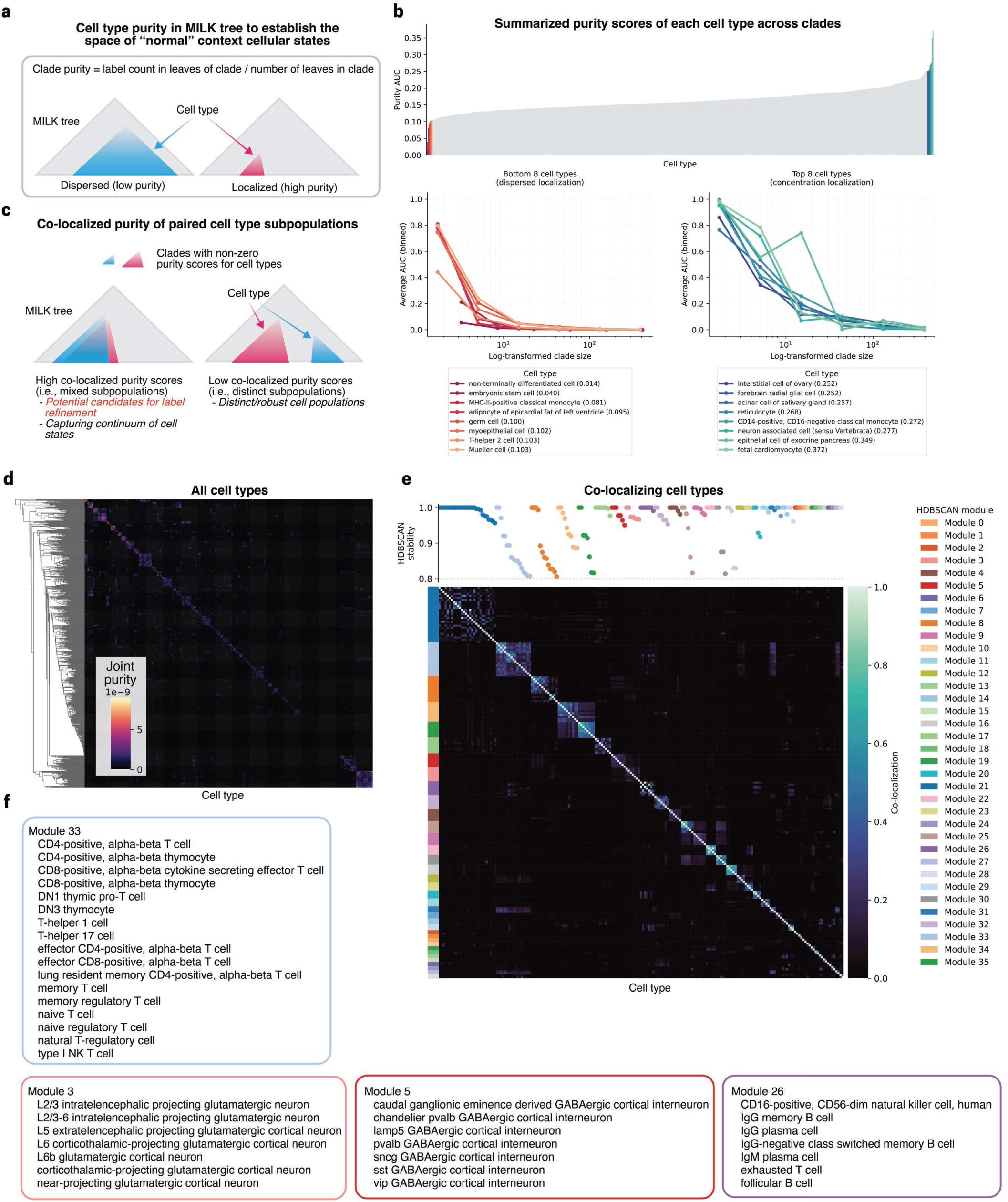
Cell type annotation coherence and refinement candidates in the normal-context MILK hierarchy. Normal-context cells were extracted from the 1 million-cell Geneformer-based MILK hierarchy to evaluate the coherence of cell type annotations in the absence of disease-context perturbation. **a,** Cell type purity in MILK clades. Schematic of clade-level cell type purity, defined as the number of cells carrying a given cell type label divided by the total number of cells in the clade. Higher purity indicates stronger localization of a cell type label within restricted regions of the MILK hierarchy, whereas lower purity indicates broader dispersion across the tree. **b,** Cell type purity across clade sizes. Purity scores were calculated for each cell type across MILK clades and summarized as an area-under-the-curve score across log-transformed clade sizes. Cell types with low purity AUC values show dispersed localization, whereas cell types with high purity AUC values show concentrated localization. The bottom- and top-ranked cell types are highlighted with purity curves across binned clade sizes. **c,** Pairwise co-localized purity of cell type labels. Schematic of joint purity between pairs of cell type labels. For each clade, joint purity was calculated as the product of the two cell type-specific purity values. High joint purity indicates that two cell type labels repeatedly co-localize within the same MILK clades, whereas low joint purity indicates stronger separation across the hierarchy. **d,** Cell type co-localization matrix. Weighted average joint purity scores were calculated across all clades for all pairs of cell type labels, using clade size as the weight. The resulting matrix summarizes pairwise co-localization of cell type annotations in the normal-context MILK hierarchy. **e,** Co-localizing cell type modules. HDBSCAN clustering was applied to the cell type co-localization matrix to identify modules of cell type labels with shared localization patterns. The heatmap shows pairwise co-localization scores, side colors denote HDBSCAN module assignments, and the top track shows HDBSCAN cluster stability for each cell type. **f,** Representative co-localization modules. Cell type labels contained in four representative HDBSCAN modules are shown. These modules may reflect biologically continuous cell states, closely related transcriptional programs, or annotation groups that could benefit from future refinement.

**Supplementary Fig. 5.10.**
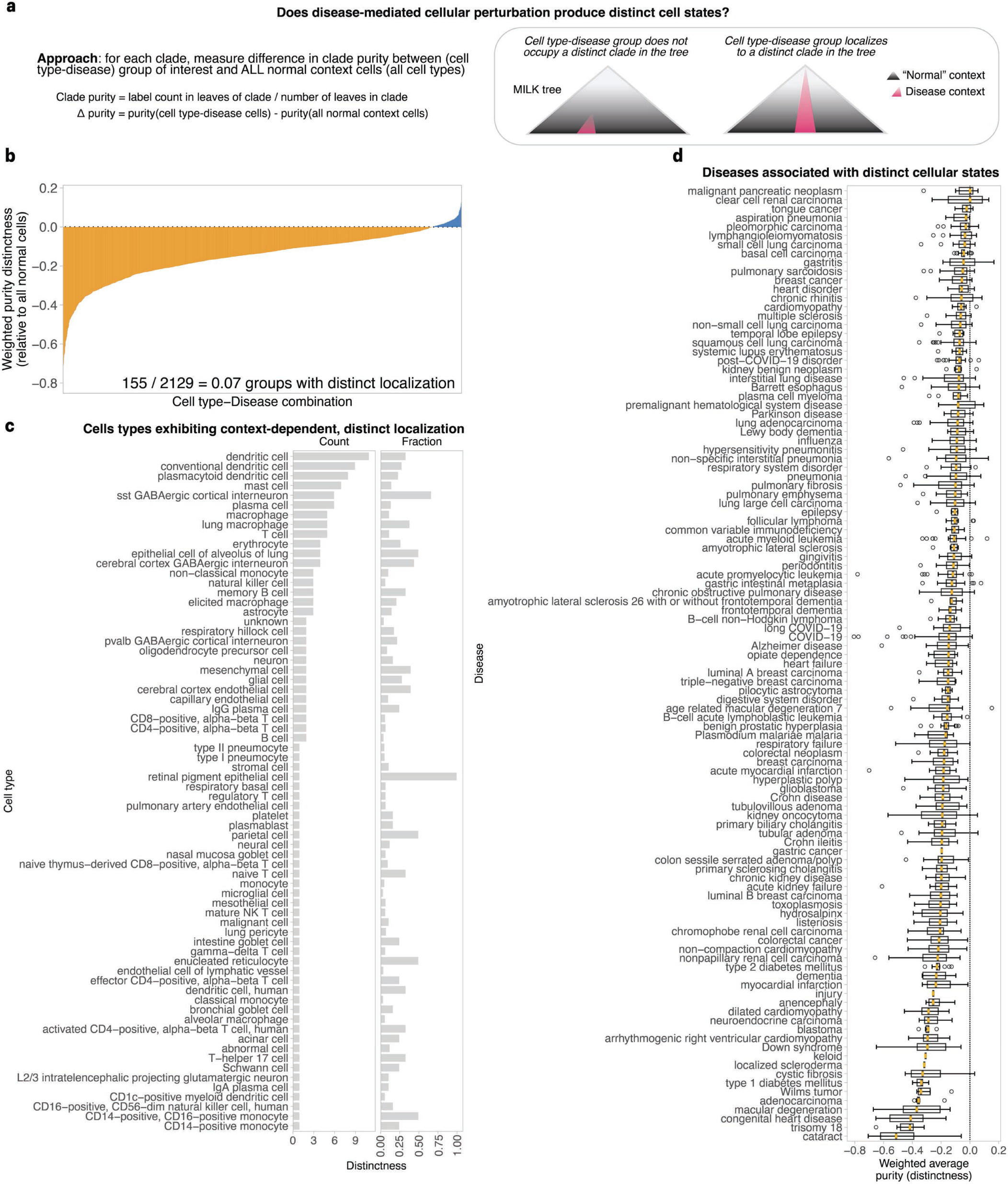
Disease-context localization distinctness relative to normal cellular states. **a,** Disease-context distinctness metric. To assess whether disease-associated cellular perturbations occupy regions of the MILK hierarchy distinct from normal-context cellular states, each cell type–disease group was compared with the pooled population of all normal-context cells. For each MILK clade, distinctness was calculated as the difference between the purity of the cell type–disease group and the purity of all normal-context cells. Positive values indicate preferential localization of the cell type–disease group in clades not broadly occupied by normal-context cells, whereas negative values indicate localization within regions represented by normal-context cell states. **b,** Distribution of disease-context distinctness scores. Clade-level purity differences were summarized for each cell type–disease group as a clade size-weighted average distinctness score. Among 2,129 cell type–disease groups, 155 groups (7%) showed positive distinctness scores, indicating disease-context localization outside the pooled normal-context cell-state space. **c,** Cell types with disease-distinct localization. Cell types with positive distinctness scores in at least one disease context were summarized by the number of disease contexts in which distinct localization was observed and by the fraction of represented disease contexts showing distinct localization. **d,** Disease-level distinctness distributions. Distributions of distinctness scores across cell type–disease groups are shown for each disease context. Boxplots summarize the extent to which cell states associated with each disease localize outside or within the pooled normal-context cellular landscape.

**Supplementary Fig. 6.1.**
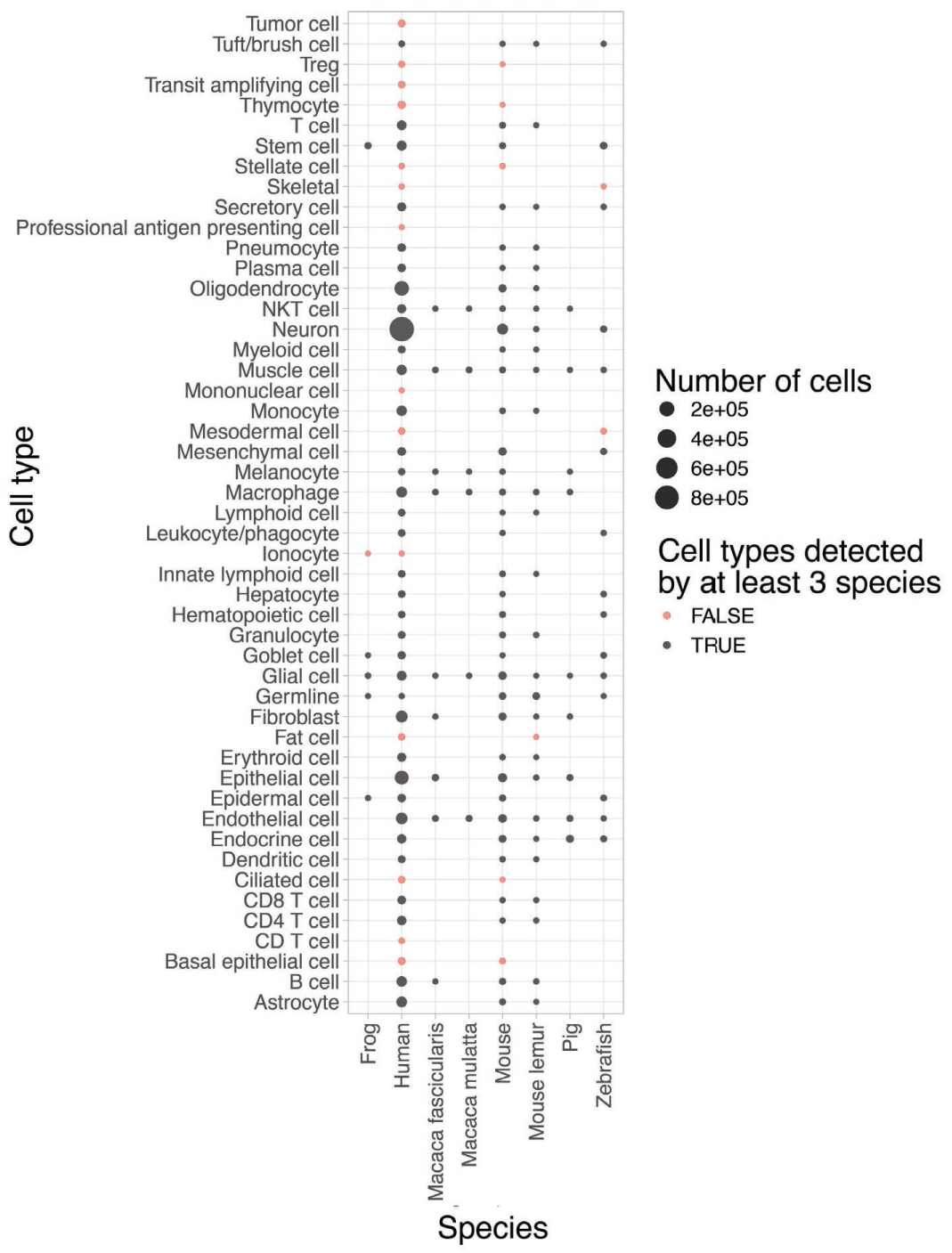
Cell type and species composition of the cross-species scRNA-seq atlas. Overview of coarse cell type representation across the cross-species atlas used for MILK analysis. A representative subset of the Integrated Mega-scale Atlas was obtained, comprising 2,969,114 single-cell UCE embeddings across eight species: frog, human, crab-eating macaque (*Macaca fascicularis*), rhesus macaque (*Macaca mulatta*), mouse, mouse lemur, pig and zebrafish. Coarse cell type labels provided by the original study were used because they enabled comparison of broadly shared cellular identities across species. Dot size denotes the number of cells for each cell type–species pair. Black dots indicate cell types represented in at least three species, whereas red dots indicate cell types represented in fewer than three species.

**Supplementary Fig. 6.2.**
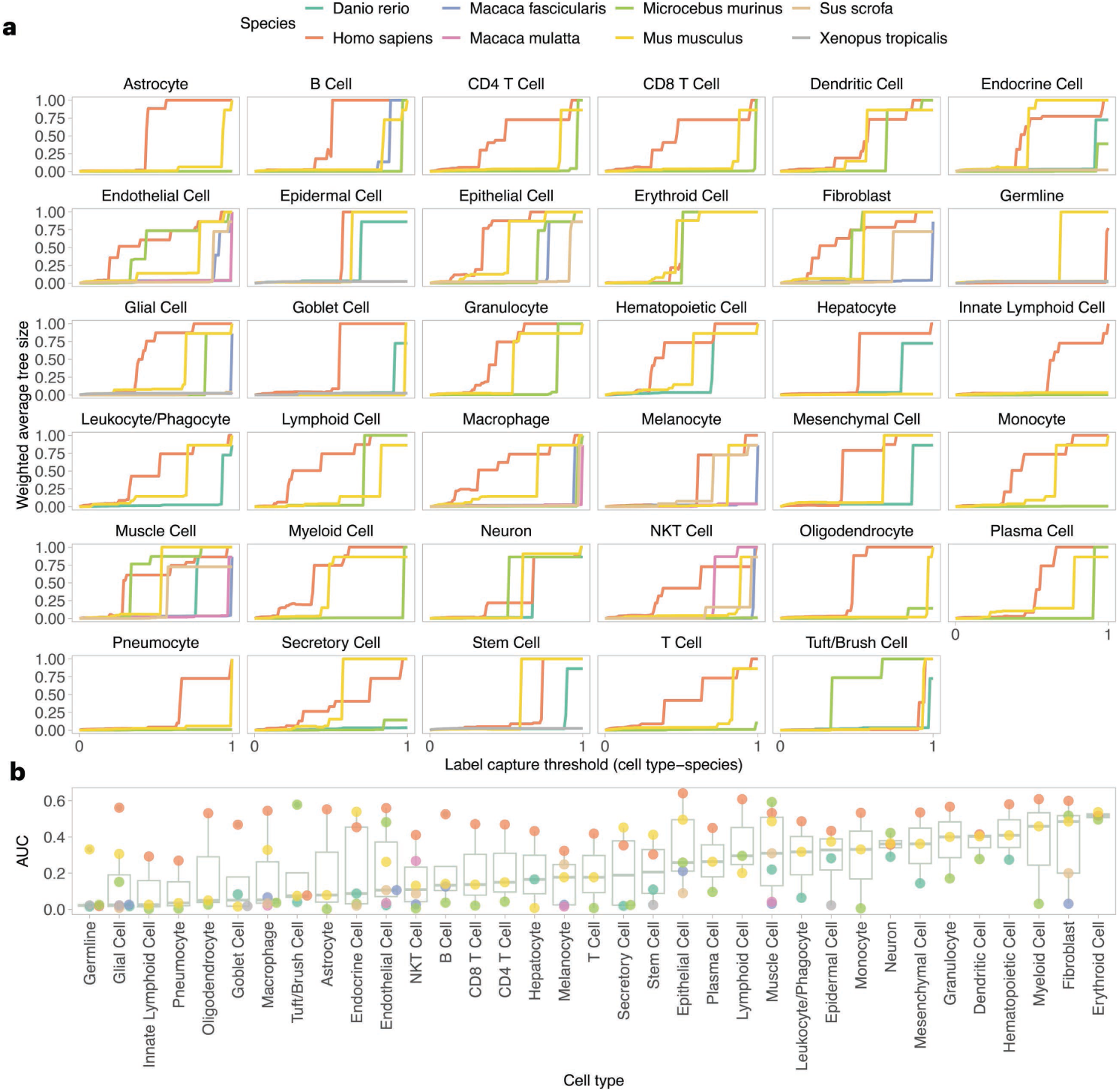
Cell type–species capture scale across the cross-species MILK hierarchy. **a,** Cell type–species label capture curves. For each cell type–species label, the characteristic clade size required to capture the label was calculated across label-capture thresholds. At each threshold, clades capturing at least the specified relative proportion of the target cell type–species population were identified, and the weighted average clade size was calculated using the purity of the target label within each clade as the weight. Clade sizes were normalized by the total number of cells in the MILK tree and plotted as a function of the label-capture threshold. Each panel shows one coarse cell type, with curves corresponding to species in which that cell type was represented. **b,** Cell type-specific capture-scale AUC. Area under the curve (AUC) values were calculated from the label-capture curves in a for each cell type–species label. Cell types were ranked by the median AUC across species. Higher AUC values indicate that larger clades are required to capture the corresponding cell type–species populations, reflecting broader dispersion across the MILK hierarchy, whereas lower AUC values indicate more localized organization.

**Supplementary Fig. 6.3.**
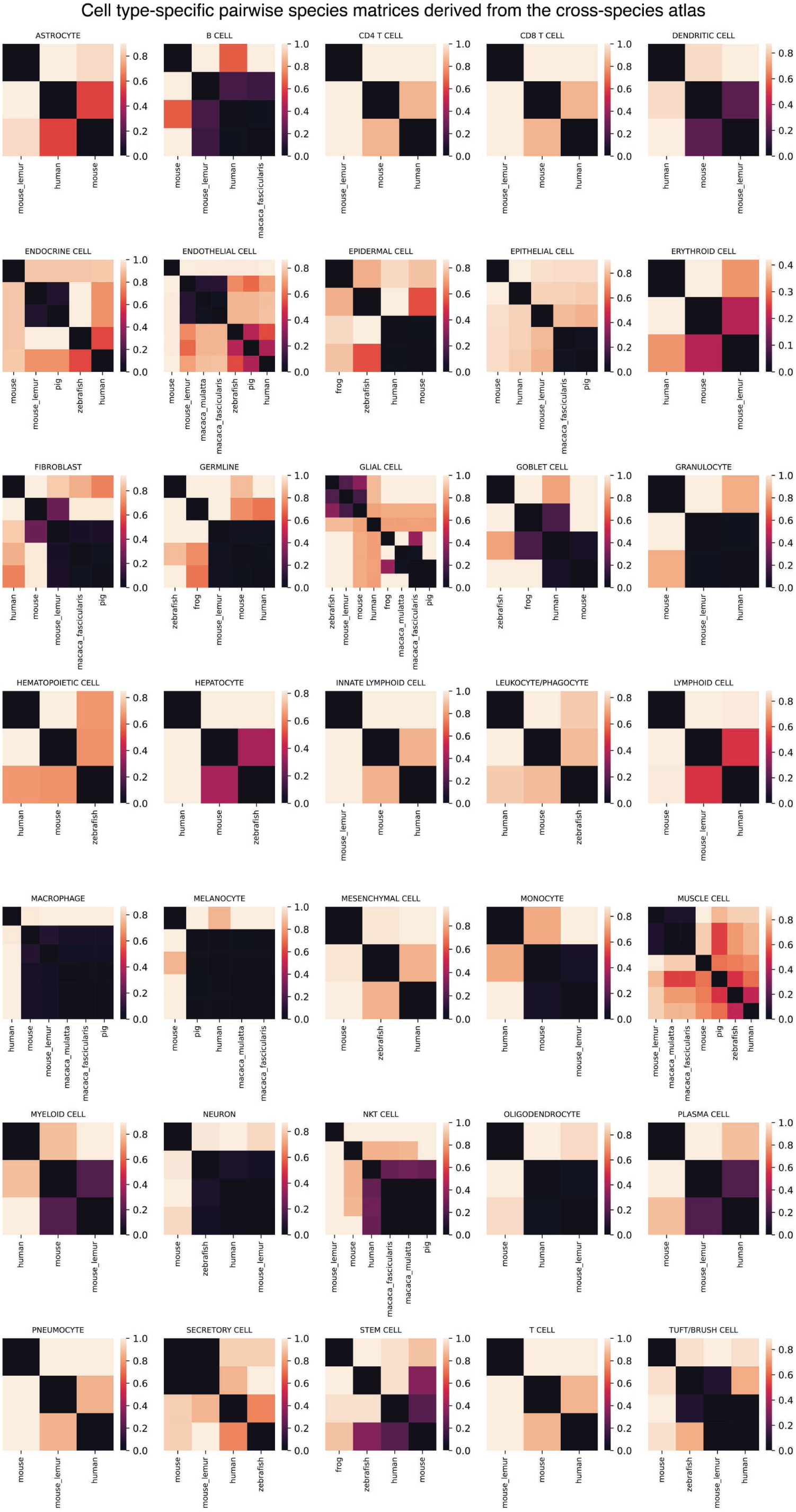
Cell type-specific species association matrices derived from the cross-species MILK hierarchy. An all-by-all pairwise association matrix was calculated for cell type–species labels based on their co-localization within the cross-species MILK tree. For each pair of cell type–species labels, association was quantified as the weighted average normalized clade size required to jointly capture both groups, using the product of their clade-level relative proportions and label purities as weights. Smaller normalized clade sizes indicate stronger co-localization within the MILK hierarchy. From the complete cell type– species association matrix, cell type-specific submatrices were extracted to evaluate species relationships within each cell type. Cell type-specific matrices containing at least three species were retained for downstream reconstruction of species trees and comparison with the reference species phylogeny.

